# Design and evaluation of novel 4-anilinoquinolines and quinazolines EGFR inhibitors in lung cancer and chordoma

**DOI:** 10.1101/545525

**Authors:** Christopher R. M. Asquith, Kaitlyn A. Maffuid, Tuomo Laitinen, Chad D. Torrice, Graham J. Tizzard, Carla Alamillo-Ferrer, Karl M. Koshlap, Daniel J. Crona, William J. Zuercher

## Abstract

Epidermal growth factor receptor (EGFR) inhibitors have been used to target non-small cell lung cancer (NSCLC) and chordomas with varying amounts of success. We have probed several key structural features including an interaction with Asp855 within the EGFR DGF motif and interactions with the active site water network. The EGFR target engagement was then evaluated in an in-cell assay. Additionally, inhibitors were profiled in representative cellular models of NSCLC and chordomas. In addition to a structure activity relationship insights for EGFR inhibtior design, we also identified a compound (**18**) that is the most potent inhibitor (IC_50_ = 310 nM) on the UCH-2 chordoma cell line to date.

---

Cancer is the second leading cause of death globally and is responsible for an estimated 9.6 million deaths in 2018.^1^ Kinases have been successfully utilized as drug targets for the past 30 years, with 38 kinase inhibitors approved by the FDA to date for mainly cancer indications.^2^ One target that has been intensely studied is epidermal growth factor receptor (EGFR). The inhibitors gefitinib and erlotinib provide significant clinical benefit in patients diagnosed with non-small cell lung cancer (NSCLC) (Fig. 1).^3–4^ The subsequent development of lapatinib as a dual EGFR and Her2 inhibitor has extended the clinical utility of EGFR inhibitors to the treatment of Her2 positive breast cancers (Fig. 1).^5^

**Figure 1.**
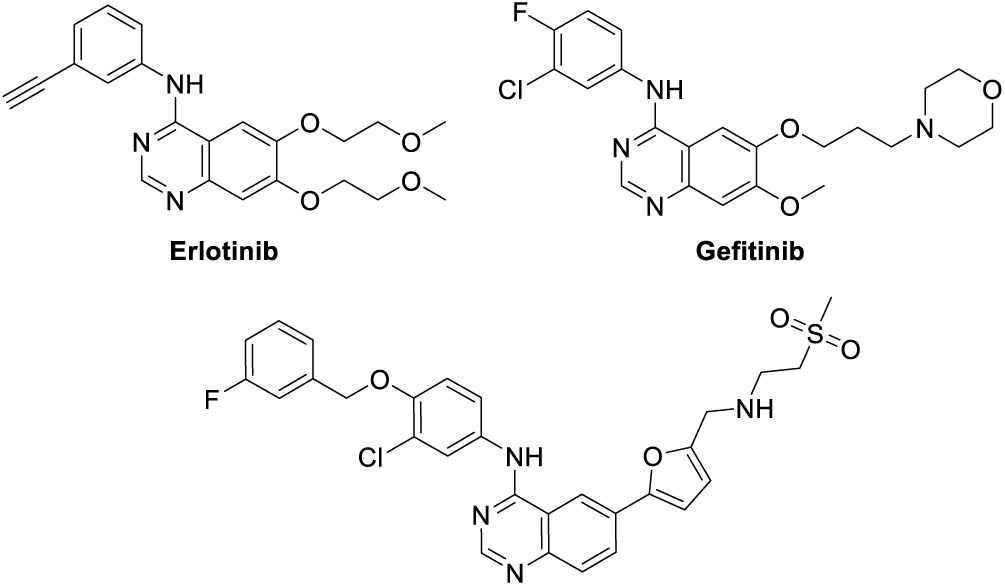
Example structures of clinical EGFR quinazolines.

Therapeutic intervention in the EGFR pathway is not limited to NSCLC and breast cancer. Other cancers show sensitivity to EGFR inhibitors.^6^ These include chordomas, which are rare tumors arising along the bones of the central nervous system and spine.^7^ These tumors are a significant challenge to treat and radical surgery is the preferred course of treatment.^7–8^ EGFR and its ligand, EGF, are highly expressed in chordomas, and copy number gains of EGFR occur in 40% of chordomas. A number of EGFR inhibitors have been identified that are active in cellular models of chordoma, and afatinib is now undergoing phase 2 clinical trials for treatment of chordoma.^9–12^

Kinase inhibitors commonly have off-targets across the kinome that confound the ability to accurately define the mechanism of action leading to induced phenotypes of interest.^13^ The 4-anilino-quinoline and quinazolines scaffold have demonstrated a range of activity profiles across the kinome from highly selective to broadly promiscuous.^14–15^ We were intrigued by lapatinib’s narrower spectrum kinome profile in addition to the longer chained aniline substituent that reduces common off-target activity by accessing a back pocket in the EGFR ATP-binding site. One significant tractable off-target is cyclin-G-associated kinase (GAK) which is frequently observed to bind 4-anilino-quinoline, quinazolines and 3-cyano-quinolines.^16^

We first designed a small series of compounds to probe the hinge region to investigate the influence of the hinge binding moiety while maintaining the simple erlotinib aniline. To determine the effect of small subtle core modification, these compounds were docked into EGFR using the Schrödinger Maestro suite (Fig. 2).^17^

**Figure 2.**
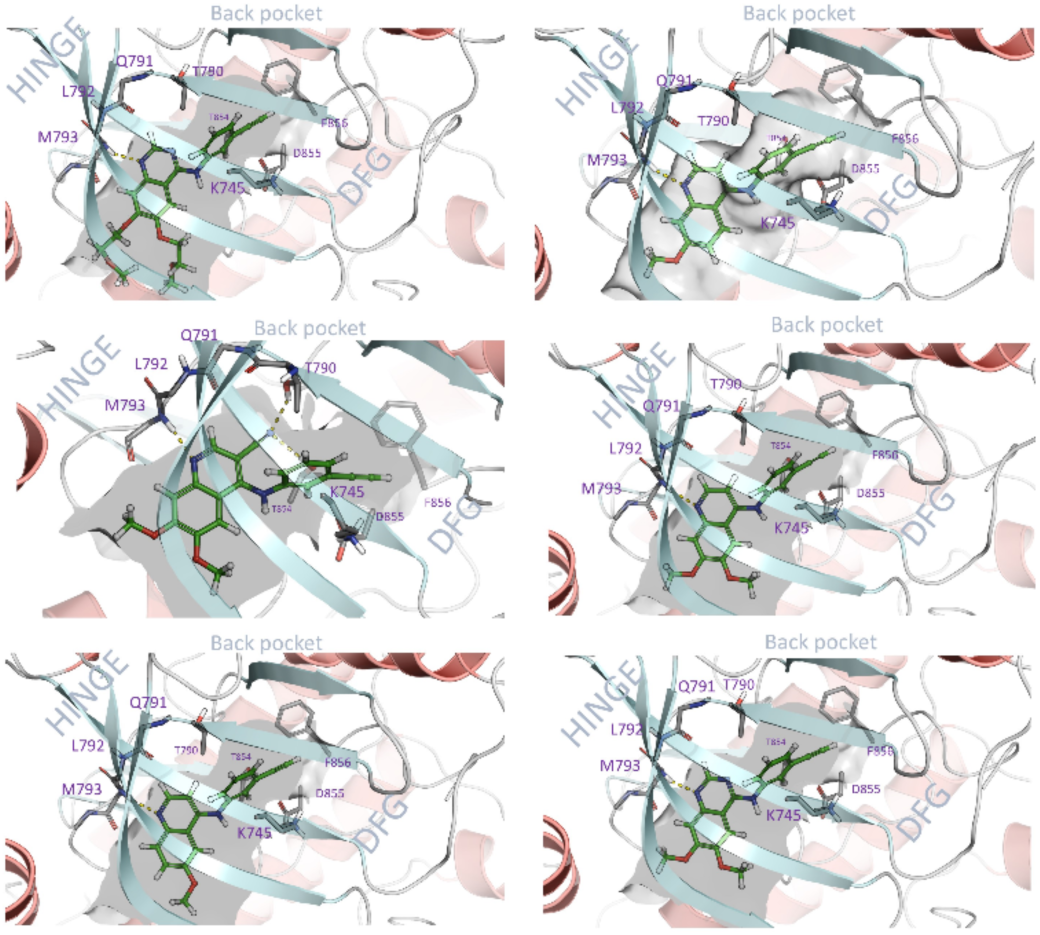
Examples of docked compounds (Left to right - erlotinib, **4**, **7**, **1**, **2**, **3**) in the ATP compteative EGFR binding domain

We prepared several focused arrays of compounds to probe the structure activity relationships of the quinoline/quinazoline series. We synthesized a series of compounds (**1**-**9**, **13**-**15**, **17**-**19**) through nucleophilic aromatic displacement of commercially available 4-chloroquin(az)olines in excellent yields (58-85 %), consistent with previous reports (Sch. 1).^14–15, 18^

These compounds were profiled in an EGFR cellular activity assay, in addition to a lung cancer cell line and several chordoma cell lines (Tab. 1). The 6,7-dimethoxyquinolin-4-amine with the erlotinib 3-ethynylaniline (**1**) showed high potency in the in-cell EGFR phosphorylation assay (IC_50_ = 270 nM), as previously reported.^19^ The data on the chordoma cell lines (UCH-1 and UCH-2) is also consistent with previous reports.^19^ The A431 lung cancer cell line showed good activity at IC_50_ = 1.4 µM and threefold weaker potency on WS-1 normal fibroblast cell viability. The removal of either methoxy group to form the 6-methoxy (**2**) or 7-methoxy (**3**) yielded compounds with more than a 60-fold drop in EGFR in cell potency. This also led to a drop off in cellular potency in the lung cancer cell line and UCH-1, there was a double in potency in UCH-2. However, the potency values of **2** and **3** were still in the low double digit micromolar range.

**Scheme 1.**
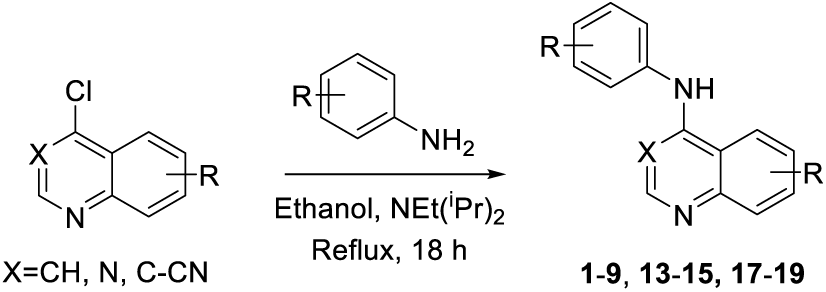
General synthetic procedure

**Table 1.** Results of a small series of compounds to structurally different hinge binders (**1**-**9**)

| Cmpd | R<sup>1</sup> | R<sup>2</sup> | R<sup>3</sup> | EGFR<sup>a</sup><br>IC<sub>50</sub> (μM) | A431 | UCH-1<br>IC<sub>50</sub> (μM)<sup>b</sup> | UCH-2<br>IC<sub>50</sub> (μM)<sup>b</sup> | WS-1 |
| --- | --- | --- | --- | --- | --- | --- | --- | --- |
| **1** | OMe | OMe | CH | 0.27<sup>c</sup> | 1.4 | 0.54 | 42 | 4.6 |
| **2** | OMe | H | CH | 16 | 10 | 6.6 | 16 | >100 |
| **3** | H | OMe | CH | >20 | 3.4 | 9.4 | 17 | >100 |
| **4** | OMe | OMe | N | 0.55<sup>c</sup> | 0.85 | 0.63 | 66 | 15 |
| **5** | OMe | H | N | 0.59 | 2.2 | 1.4 | 47 | >100 |
| **6** | H | OMe | N | 0.53 | 8.0 | 1.9 | 1.2 | >100 |
| **7** | OMe | OMe | C-CN | 1.8<sup>c</sup> | 1.7 | 4.1 | 36 | >100 |
| **8** | OMe | H | C-CN | 3.5 | >100 | 19.6 | >100 | >100 |
| **9** | H | OMe | C-CN | 1.1 | 44 | 4.1 | 40 | >100 |

<sup>a</sup>ProKinase In-cell assay (n=1), <sup>b</sup>(n=3)

The switch to the quinazoline (**4**) showed a similar potency range to **1** with a slight increase in potency in A431 cells. The removal of either methoxy (**5** and **6**) had no impact on in cell EGFR activity but did reduce activity in all 3 cancer cell lines with no effect in WS-1. The 3-cyano quinoline hinge binder showed a marked drop off in cell EGFR activity. The 6,7-dimethoxy analog (**7**) showed similar potencies to the mono-methoxy quinazolines (**5** and **6**). It was then surprising that the removal of either methoxy (**8** and **9**) reduced the anti-proliferative effect seen in other analogs (**7**), suggesting the involvement of other targets.

With the results of the small focused series in hand, we then started to modify the aniline scaffold, with the aim of establishing an internal hydrogen bond, not only to form a pre-organized structure but also to form an interaction with Asp855 as in Figure 3. In tandem we looked at the water network in EGFR (Fig. 4) and found that by adding a methyl group to the pendent benzyl we were able to increase the ligand efficiency of our model system.^19^

**Figure 3.**
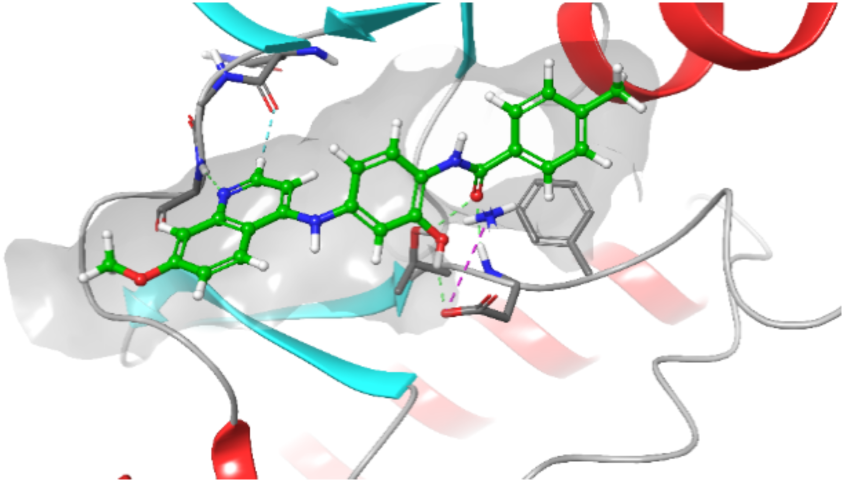
Docking of **15** with an alcohol substitution into EGFR

**Figure 4.**
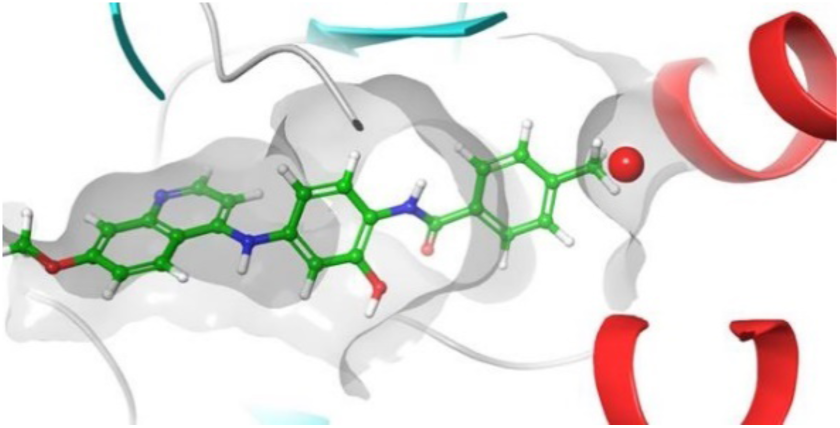
WaterMap simulation of **15** in EGFR

**Scheme 2.**
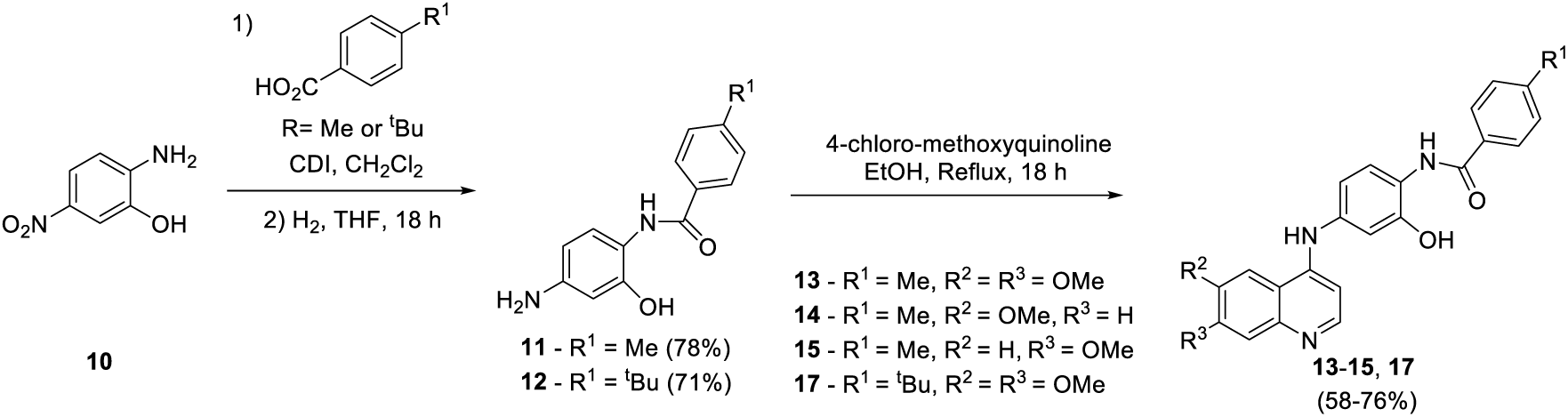
Synthetic procedure for **13**-**15** and **17**

**Scheme 3.**
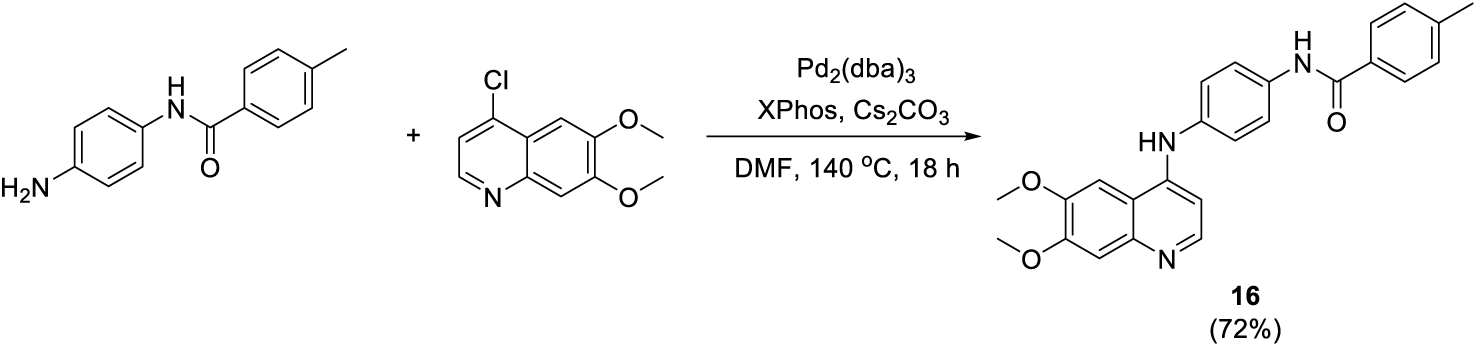
Synthetic procedure for **16**

We prepared several additional compounds (**13**-**18**) where the anilines were not readily available as the ones in Scheme 1, following a three-step protocol from commercially available 2-amino-5-nitrophenol (**10**) as starting material. **10** was coupled with CDI to furish an amide bond, followed by H_2_ reduction to give intermediates **11** and **12** (Sch. 2). S_N_Ar reactions of intermediates **11** and **12** under reflux provided compounds **13**-**15** and **17** in good yields (58-76%).^18^ The des-hydroxy compound (**16**) proved inaccessible *via* a nucleophilic aromatic displacement (even up to 150 °C, DMF, DIPEA, 18 h) and required Buchwald-Hartwig conditions (Sch. 3) to produce **16** in good overall yield (72%).^15^

The 6,7-dimethoxyquinolin-4-amine with the lapatinib derived hydroxy amide aniline (**13**) showed double digit micromolar potency in the in-cell EGFR phosphorylation assay and moderate activity in the A431 cell line and good activity in both chordoma cell lines tested. Removal of one of the methoxy groups (**14** and **15**) had the opposite effect to the previous set (**1**-**9**) and led to an increase in activity in the chordoma cell lines with no change in the A431 anti-proliferative effect.

Interestingly our hypothesis of including the alcohol proved to be pivotal for activity, with the removal of the alcohol (**16**) removing the bulk of the activity previously observed in **13**-**15**. We also considered increasing the bulk *tert*-butyl on the pendant benzyl (**17**) to more fully occupy the displaced water pocket. This only led to additional molecular weight with no potency gain compared with **13**. It was surprising that the *para*-methyl benzylic ether substitution (**18**) with no alcohol showed no in-cell EGFR activity and a limited effect in A431 cells. However, in the two chordoma cell lines there was a sharp increase in activity. **18** is one of the most potent compounds seen to data with an IC_50_ of 330 nM and 310 nM for UCH-1 and UCH-2 respectively. This result is more impressive considering that UCH-2 is typically less sensitive to compound treatment. However **18** does show some observed toxicity in WS-1. The removal of the benzyl in **19** had a similar effect to removing the alcohol in **16** with most activity lost, but moderate potency in A431 was still observed.

We were also interested in the conformation of **13** to see if there would be a rigidity imparted by the alcohol onto the pendant arm structure that was not observed in **16**. The small molecule crystal structure of **13** was solved as a monoclinic structure with a 27:73 to the pre-organization to the internal seven membered ring system (Fig 5).^20^ This system would likely be more ordered, however under our crystalisation conditions the alcohol forms a 2.99Å hydrogen bond with the chloride ion. This significant electrostatic component within the lattice acts as an anchor, hindering further pre-organisation.

**Figure 5.**
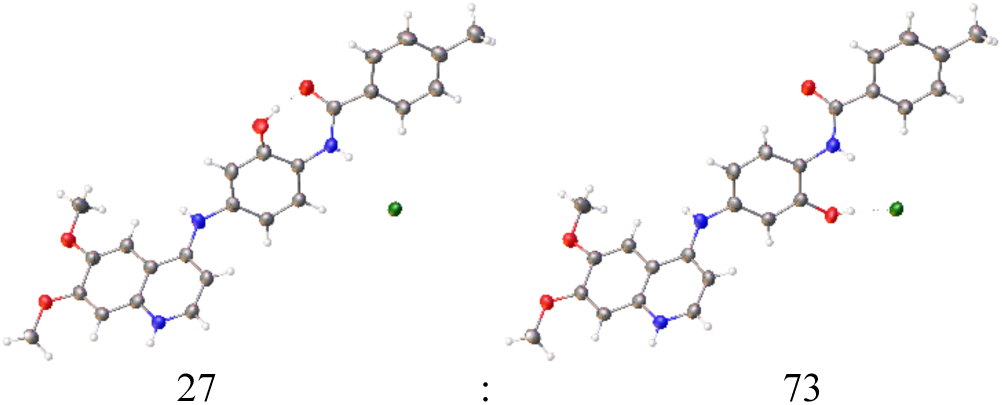
Small molecules crystal structure of **13**

**Table 2.** Matched pair comparison of benzyloxyaniline

| Compound | R<sup>1</sup> | R<sup>2</sup> | R<sup>3</sup> | R<sup>4</sup> | EGFR<sup>a</sup> | A431 | UCH-1 | UCH-2 | WS-1 |
| --- | --- | --- | --- | --- | --- | --- | --- | --- | --- |
|  |  |  |  |  | IC<sub>50</sub> (uM) | EC<sub>50</sub> (uM) |  |  |  |
| **13** | OMe | OMe | OH | **A** | 14 | 1.6 | 1.6 | 2.4 | 1.4 |
| **14** | OMe | H | OH | **A** | 14 | 1.8 | 0.93 | 1.4 | 1.4 |
| **15** | H | OMe | OH | **A** | 6.3 | 1.7 | 0.49 | 0.68 | 0.86 |
| **16** | OMe | OMe | H | **A** | >20 | 21 | 14 | 21 | >100 |
| **17** | OMe | OMe | OH | **B** | 13 | 1.8 | 1.2 | >100 | 1.9 |
| **18** | OMe | OMe | H | **C** | >20 | 1.5 | 0.33 | 0.31 | 1.1 |
| **19** | OMe | OMe | H | **D** | >20 | 3.4 | 15 | 15 | 3.8 |

<sup>a</sup>ProKinase In-cell assay (n=1), <sup>b</sup>(n=3)

EGFR inhibitors have been used to target NSCLC and chordomas, with variation in efficacy across inhibitors and increasing resistance to inhibitors, particularly in the case of NSCLC.^21^ We have highlighted a series od modifications investigating the effect of key structural features on the quin(az)oline scaffold. These modifications can be used to enhance or reduce EGFR activity and generally have a pronounced effect on cellular potency.

One of the key results observed was with the removal of one the methoxy groups from **1** leading to a significant drop off in in-cell EGFR and anti-proliferative effects. Interestingly this is not observed in the other quinazoline and 3-cyanoquinoline templates. This was also less significant when looking at the extended aniline structure of **13**-**16** where the alcohol interaction with Asp855 and conformational rigidity were more significant. It was also clear that the benzyl substitution had a significant contribution to the activity despite not been in the key hinge binding interaction. The most surprising result was compound **18**, which showed limited EGFR activity but the most potent anti-proliferative effect in both UCH-1 and UCH-2 chordoma cell lines. Despite some toxicity observed in WS-1 cells, **18** could prove to be an interesting starting point for further investigations in chordomas and NSCLC and highlights the complexity of both cancer biology and target engagement.

## Acknowledgments

The SGC is a registered charity (number 1097737) that receives funds from AbbVie, Bayer Pharma AG, Boehringer Ingelheim, Canada Foundation for Innovation, Eshelman Institute for Innovation, Genome Canada, Innovative Medi-cines Initiative (EU/EFPIA) [ULTRA-DD grant no. 115766], Janssen, Merck KGaA Darmstadt Germany, MSD, Novartis Pharma AG, Ontario Ministry of Economic Development and Innovation, Pfizer, São Paulo Research Foundation-FAPESP, Takeda, and Wellcome [106169/ZZ14/Z]. We also thank CSC - IT Center for Science Ltd. Finland for the use of their facilities, software licenses and computational resources. We are grateful Dr. Brandie Ehrmann for LC-MS/HRMS support provided by the Mass Spectrometry Core Laboratory at the University of North Carolina at Chapel Hill.

## Supplementary Material

Supplementary material

