## Supplementary Information for "Design and evaluation of novel 4-anilinoquinolines and quinazolines EGFR inhibitors in lung cancer and chordoma"

**Contents**

1.1. Compound Characterization Spectra (**2**-**9** & **13**-**19**)

1.2. Smiles and Labbook codes for final compounds

1.3. EGFR in cell phosphorylation curves and method

1.4. IC50 curves for A431, UCH-1, UCH-2 and WS-1

1.5. Mass Spectrometry method

1.1. **Compound Characterization Spectra (2-9 & 13-19)**

***N*-(3-ethynylphenyl)-6-methoxyquinolin-4-amine** (**2**)

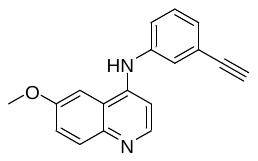

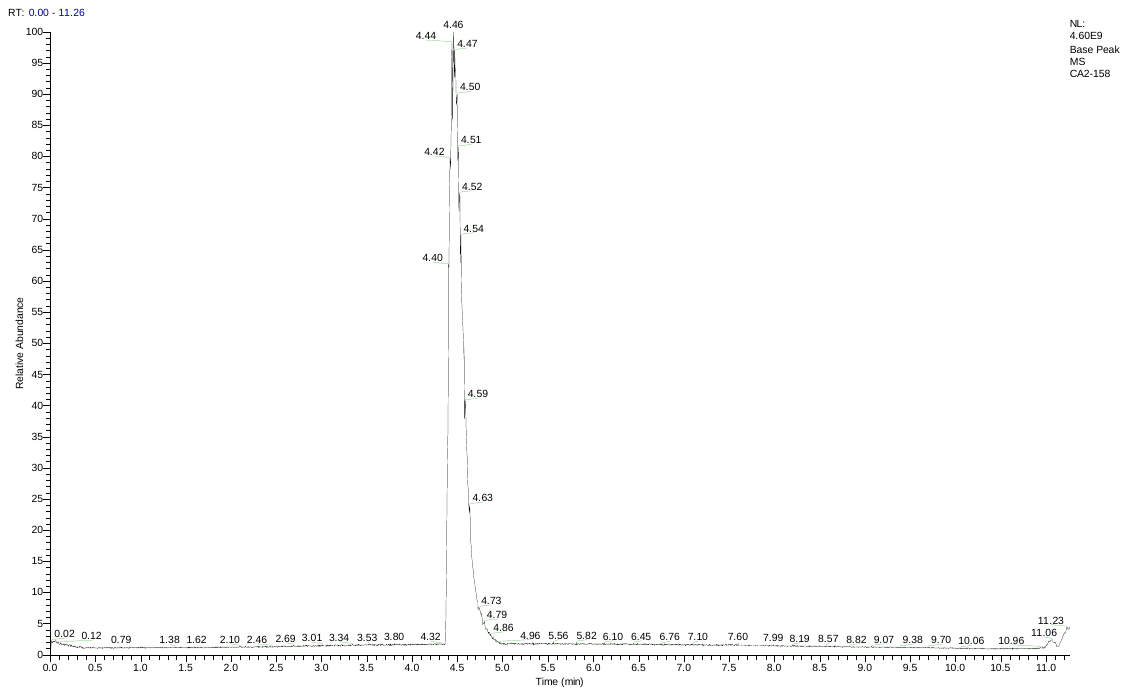

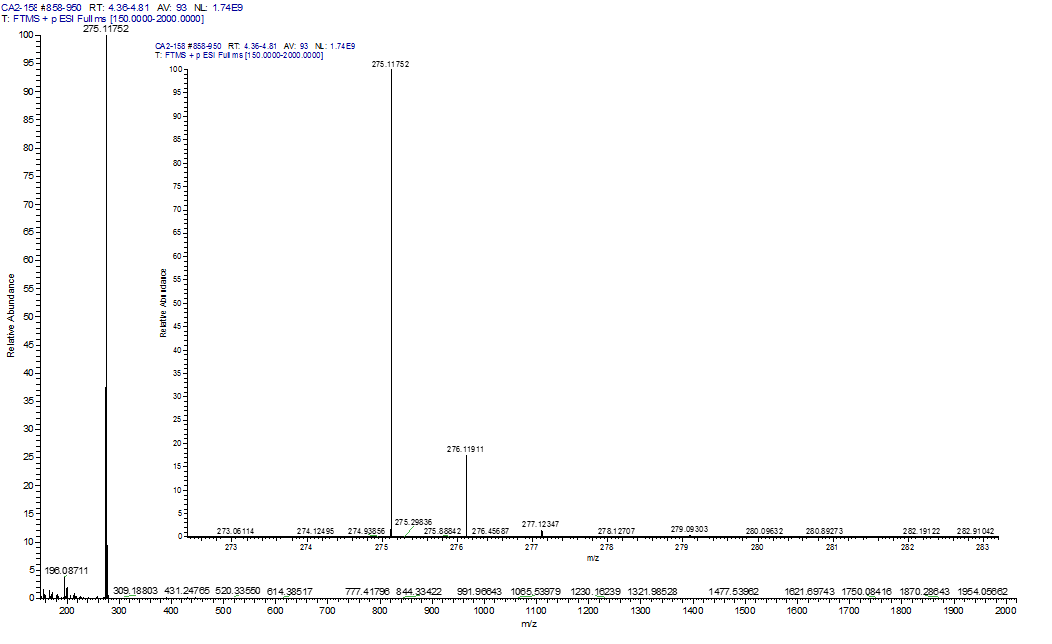

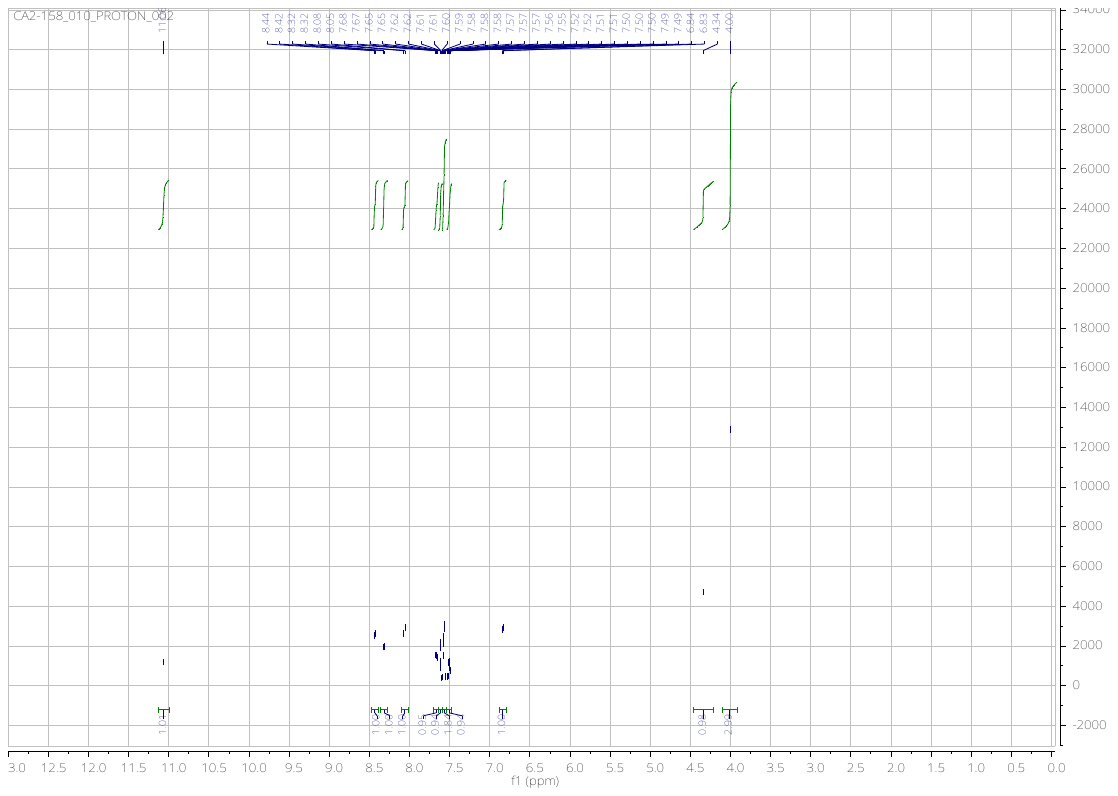

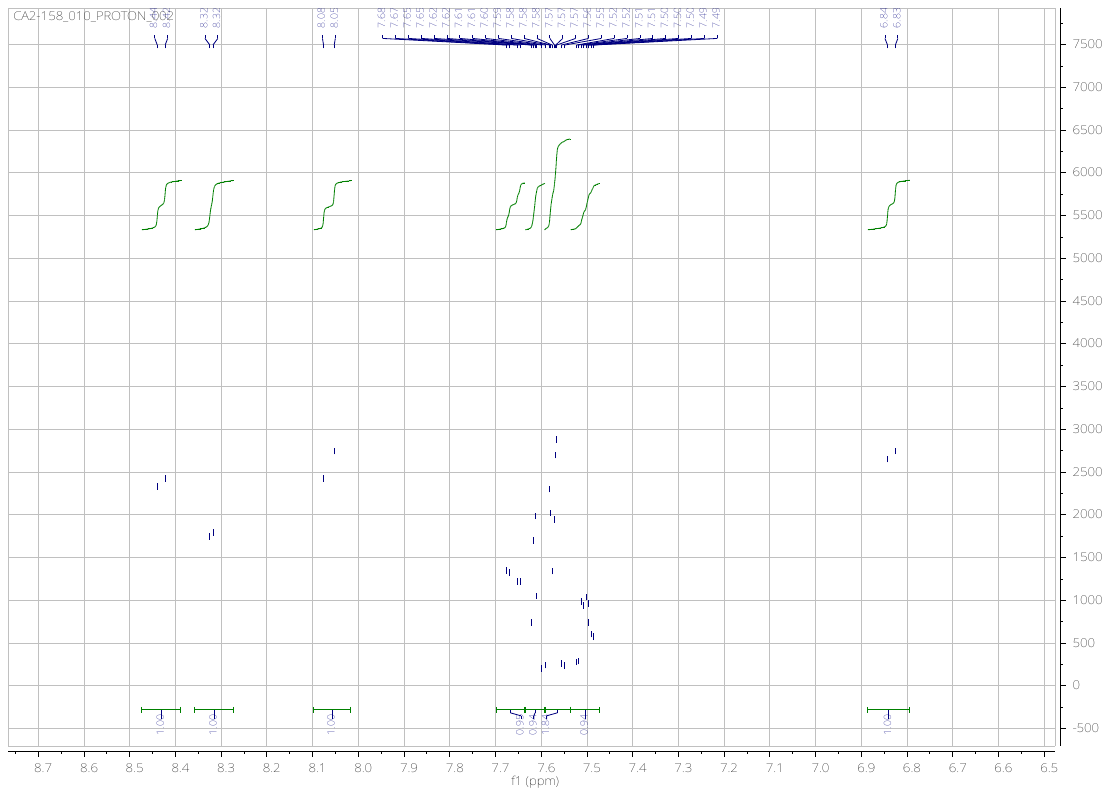

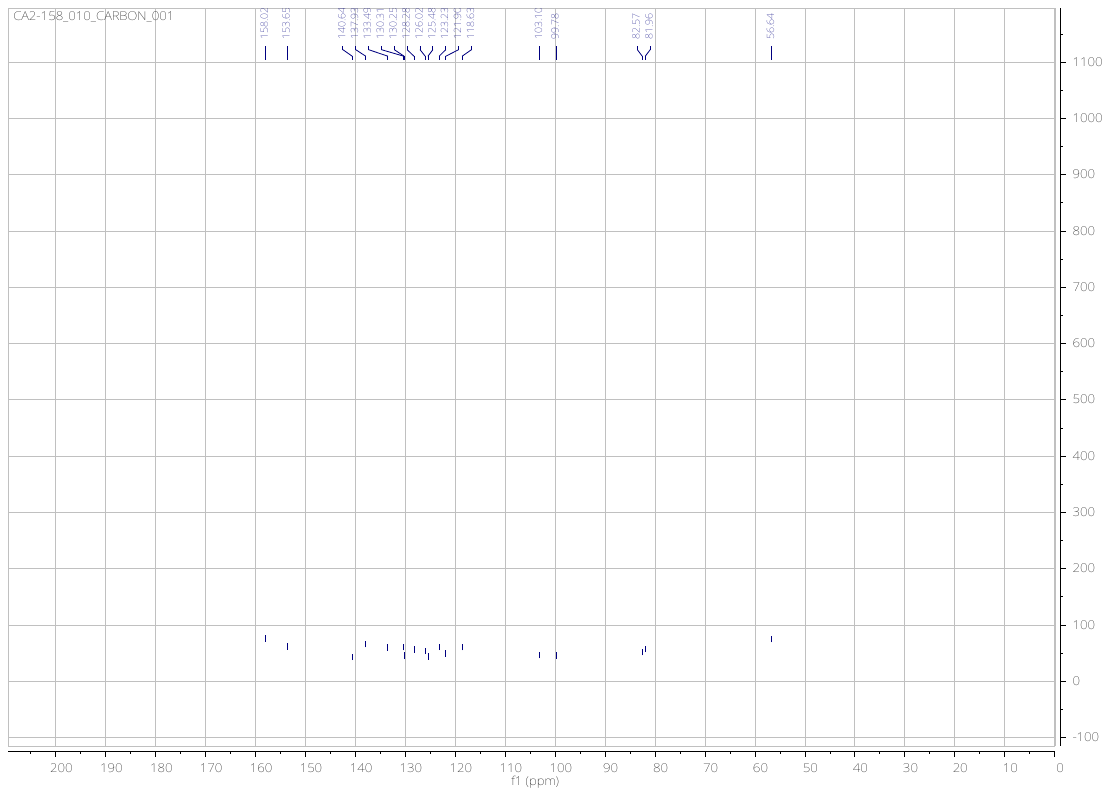

***N*-(3-ethynylphenyl)-7-methoxyquinolin-4-amine** (**3**)

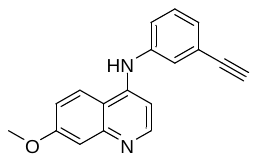

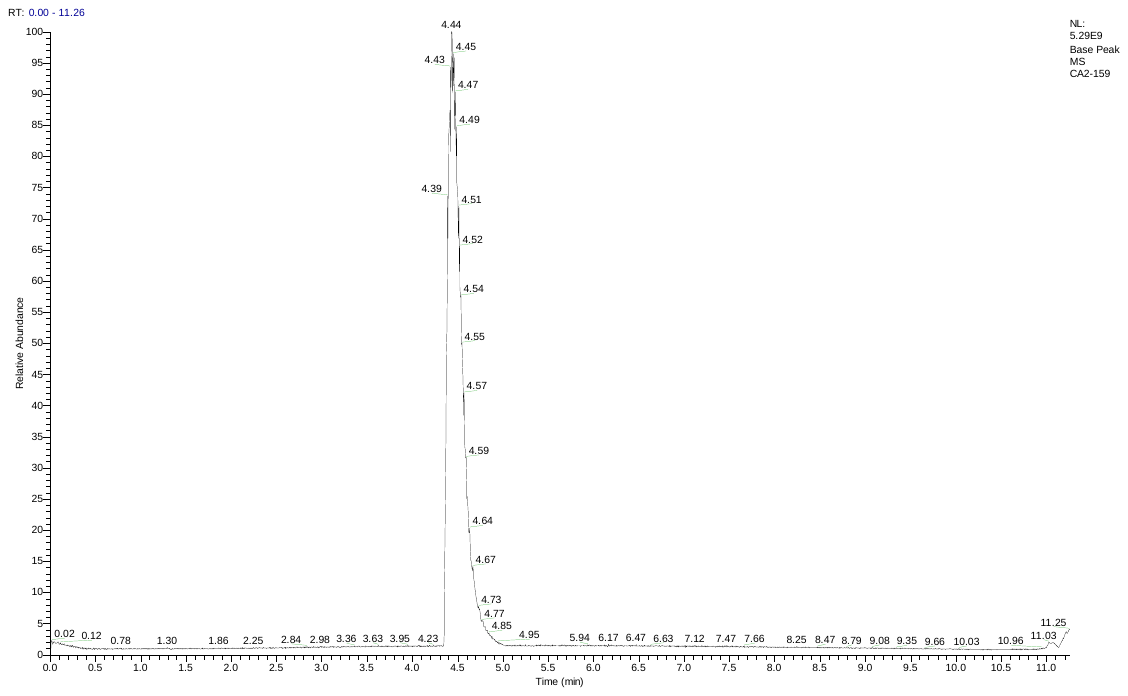

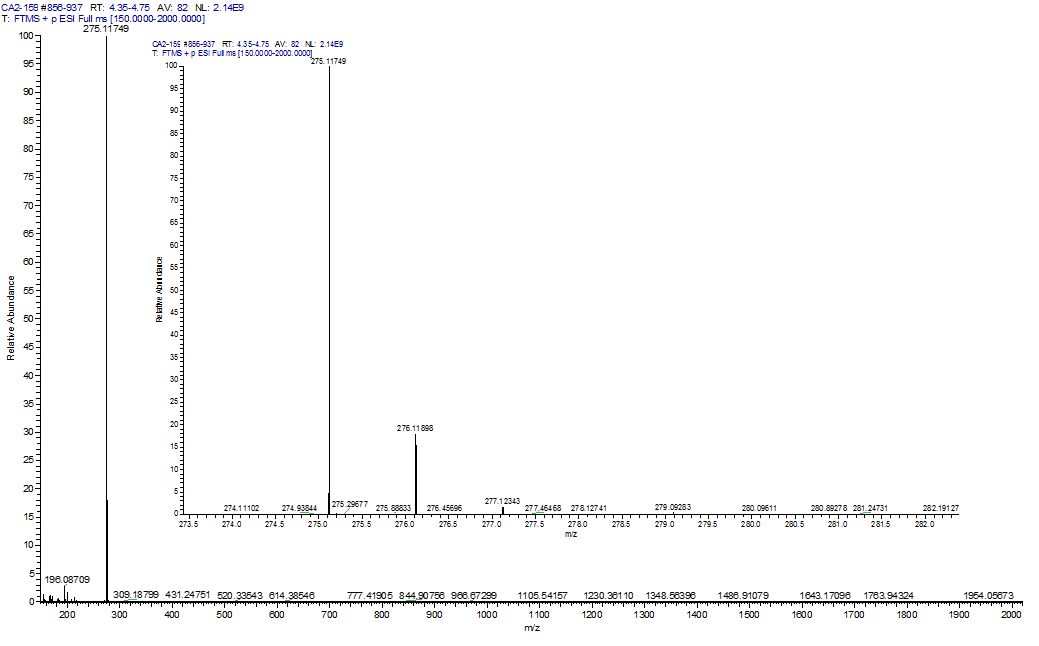

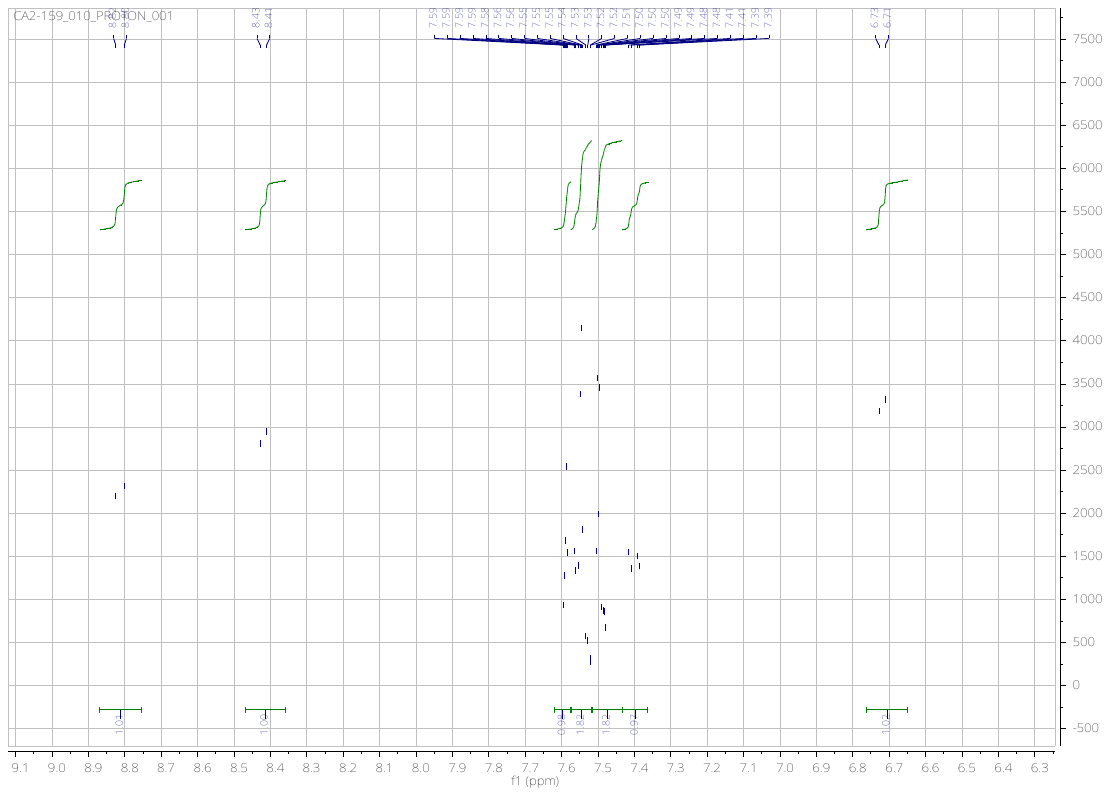

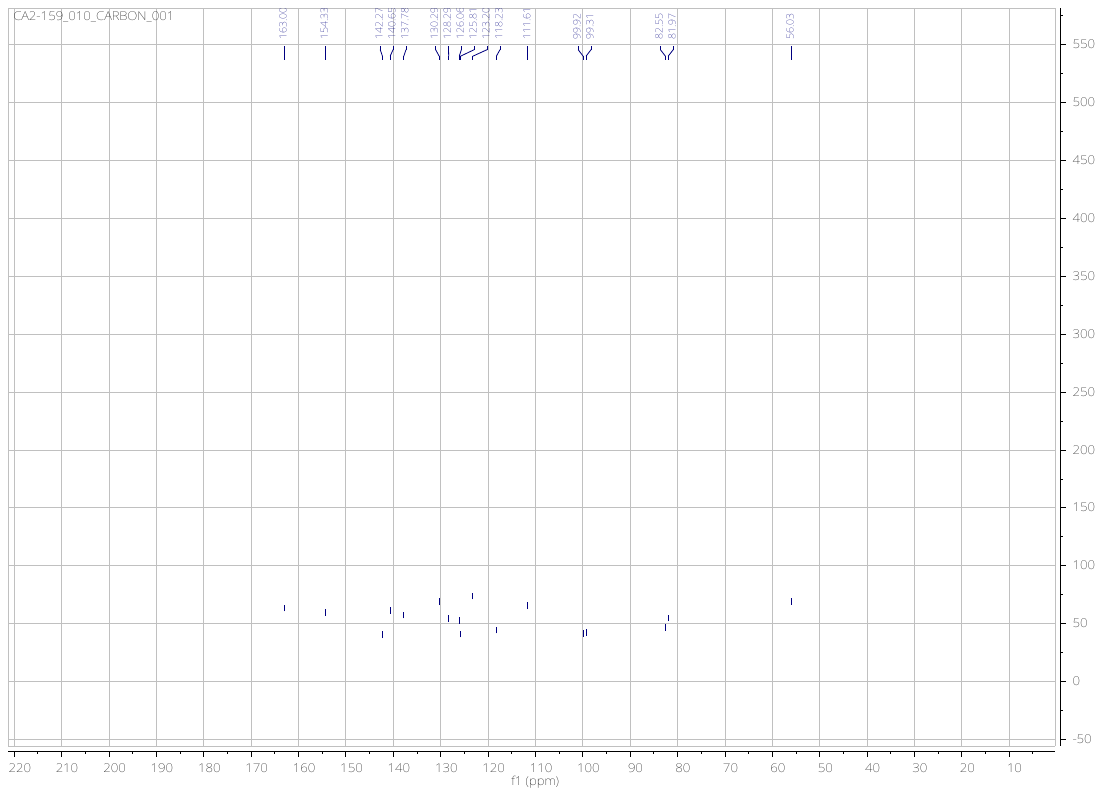

***N*-(3-ethynylphenyl)-6,7-dimethoxyquinazolin-4-amine** (**4**)

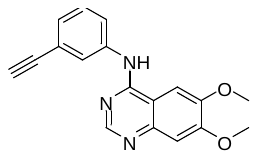

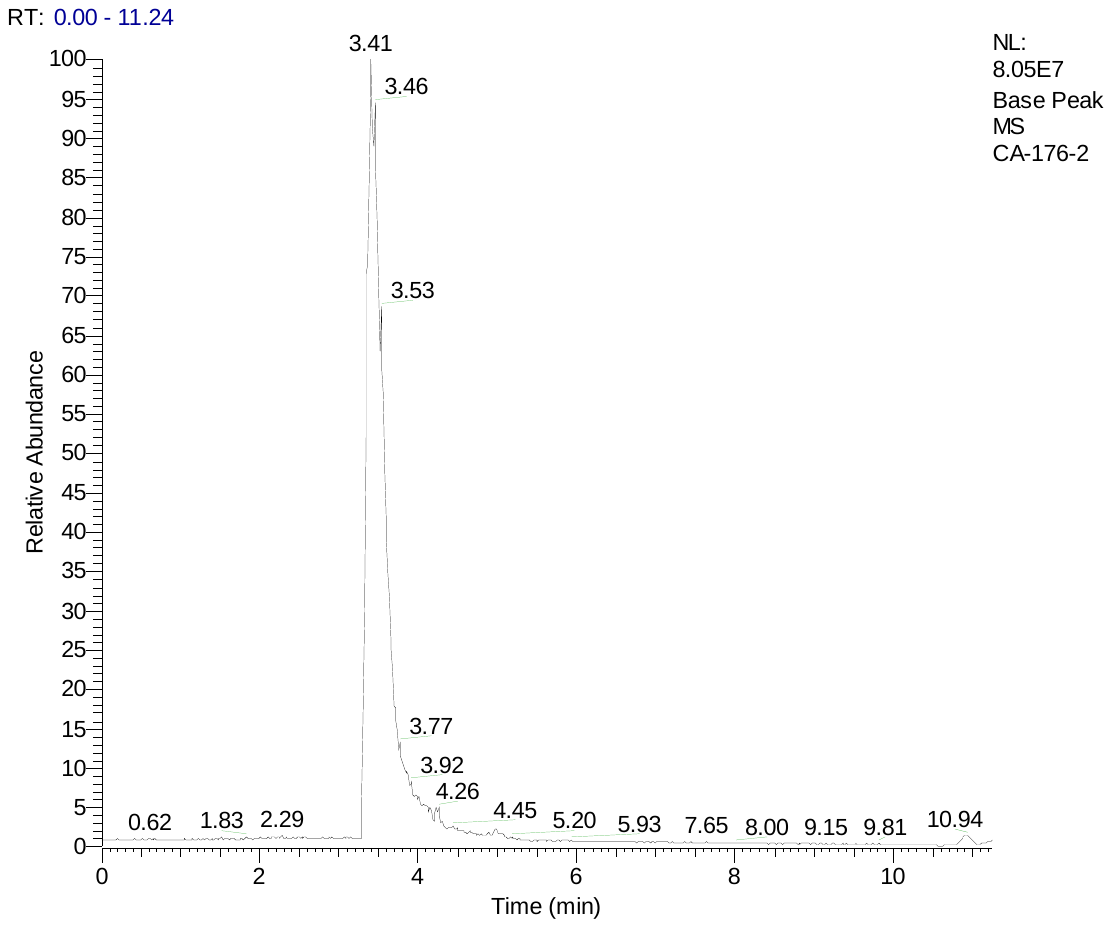

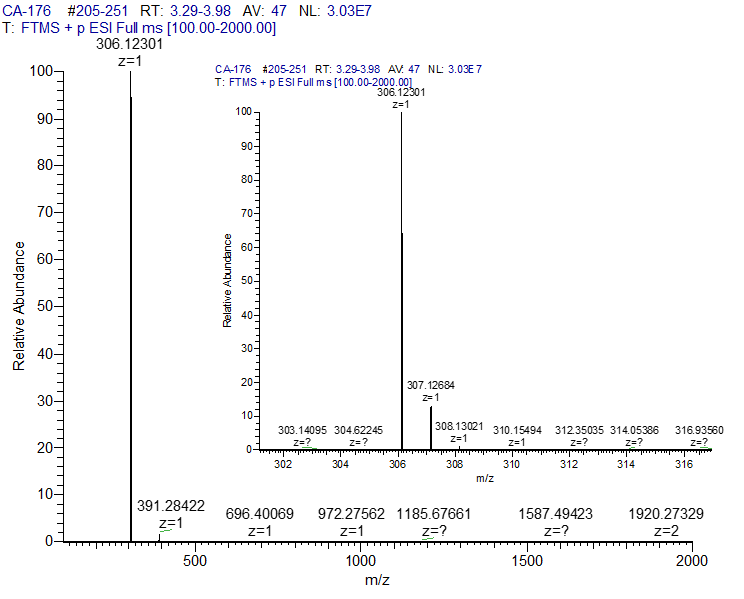

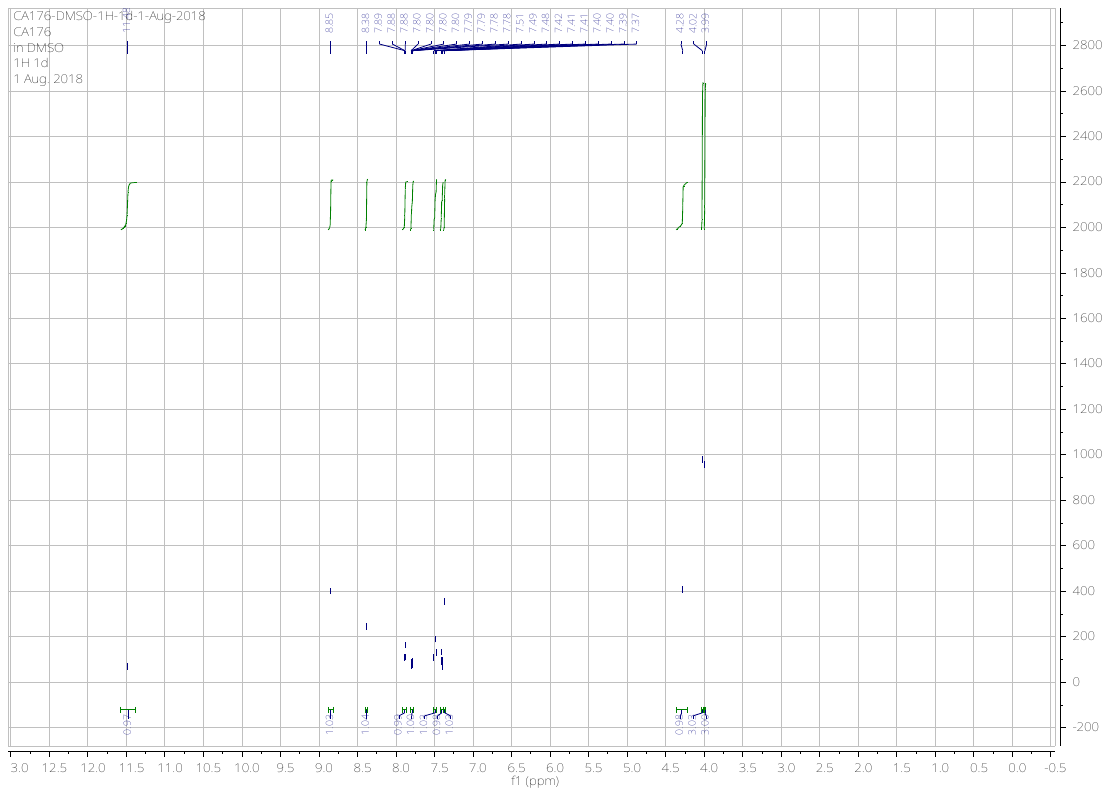

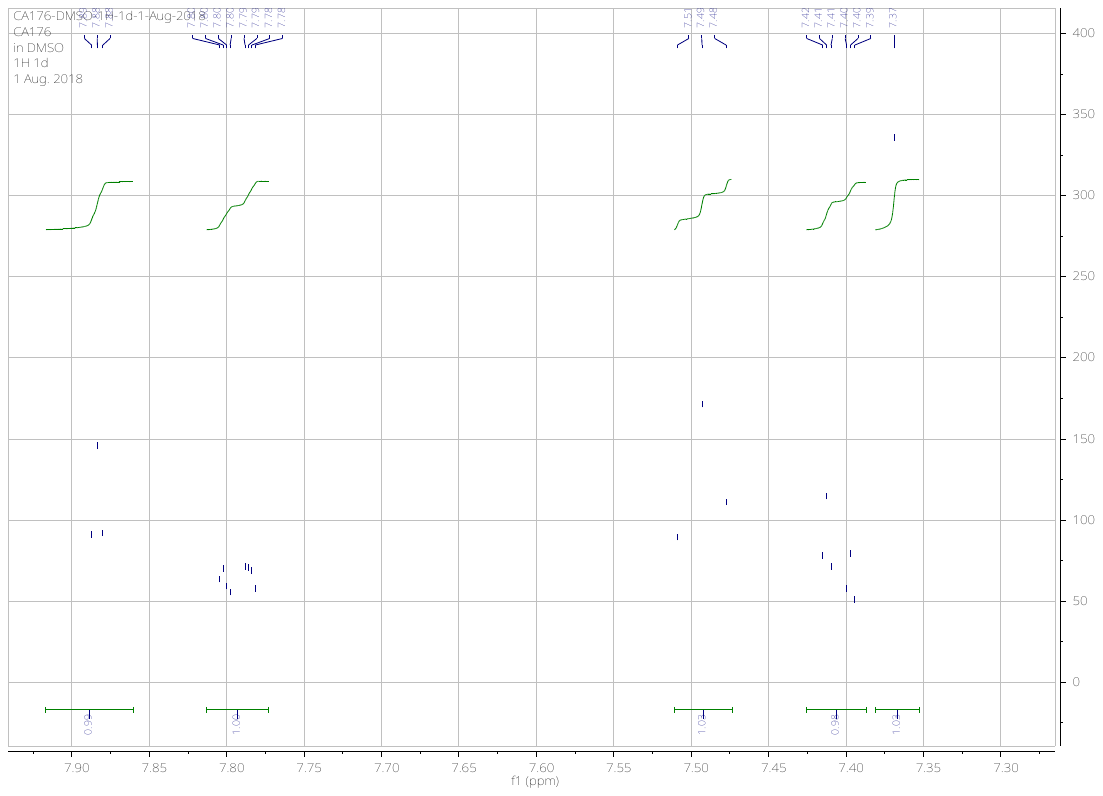

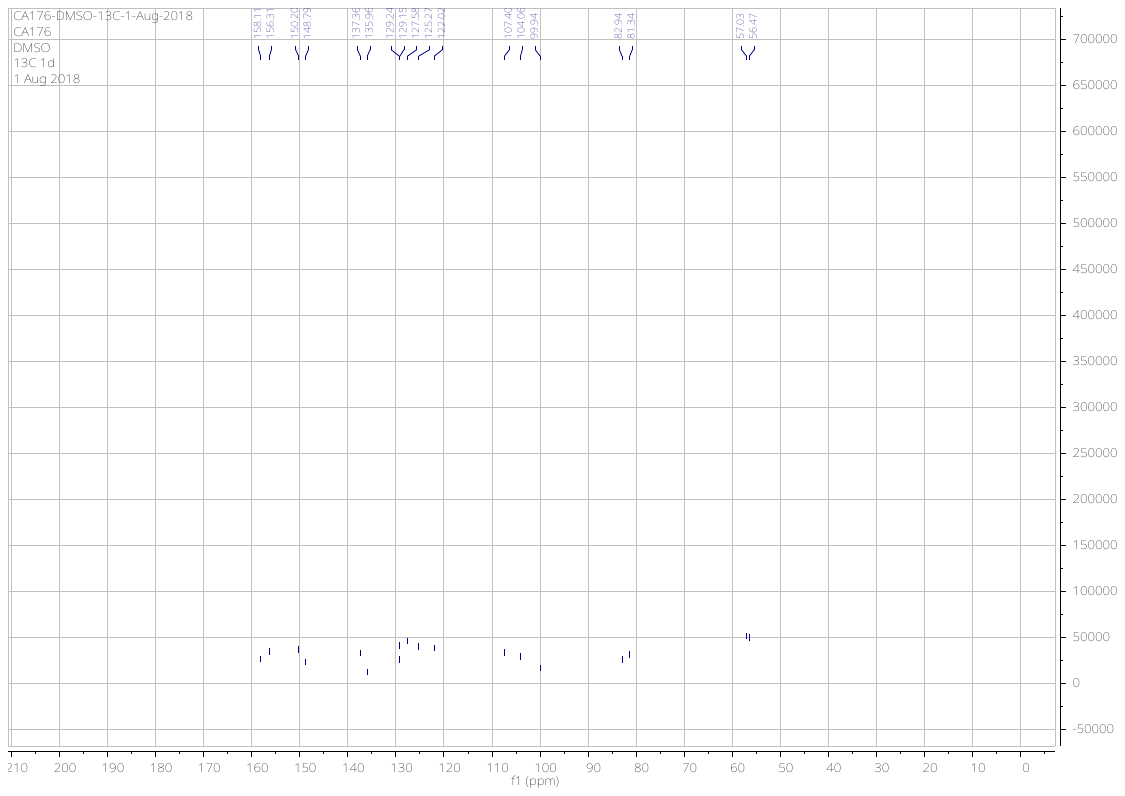

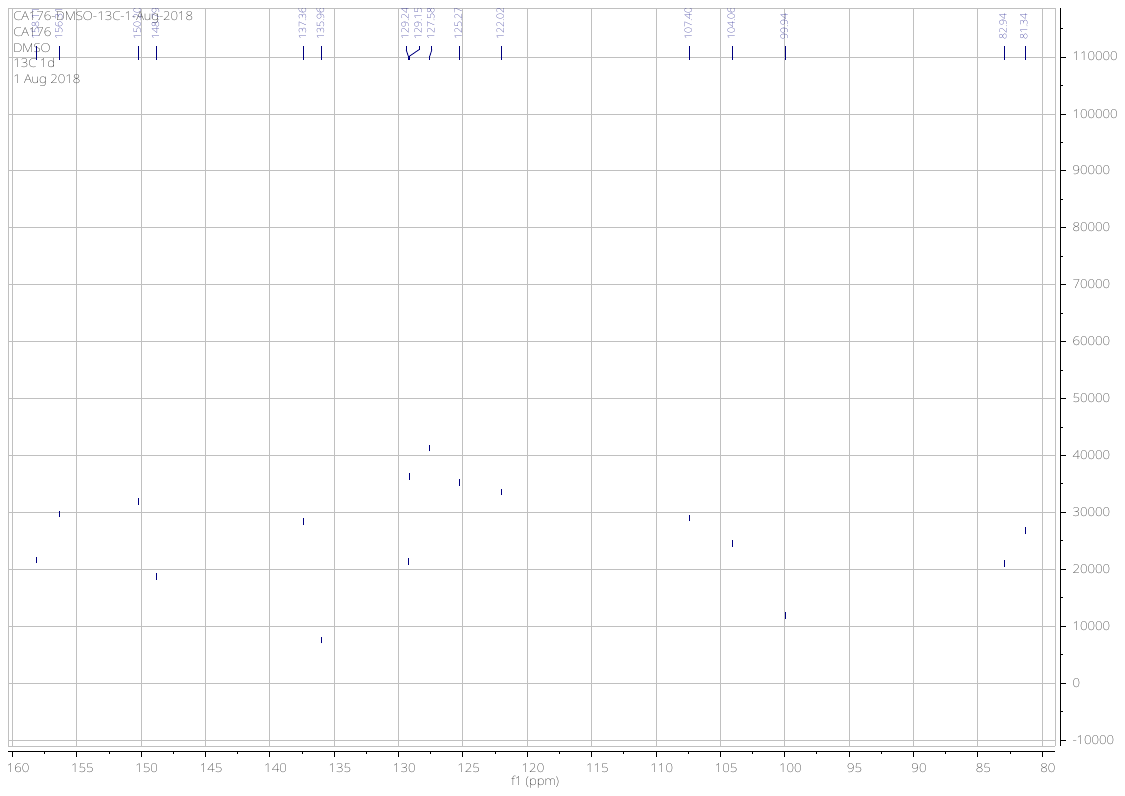

***N*-(3-ethynylphenyl)-6-methoxyquinazolin-4-amine** (**5**)

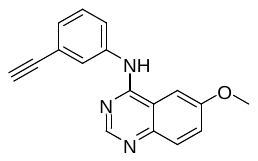

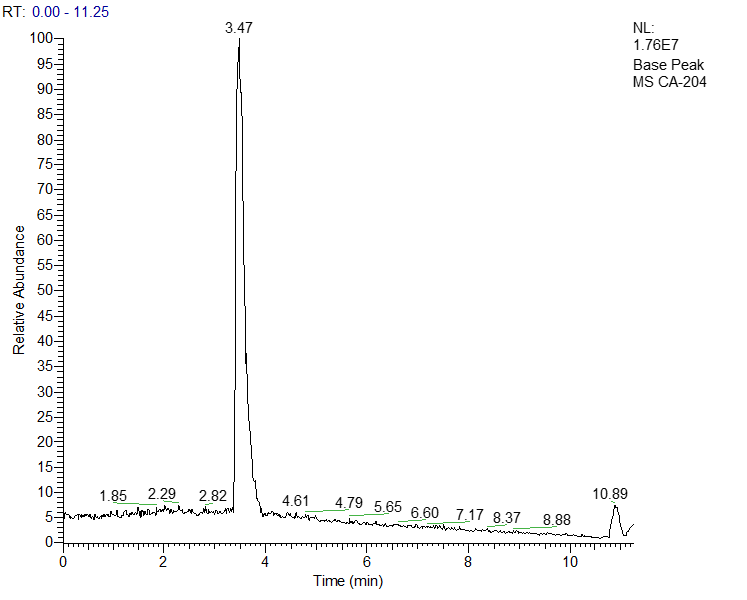

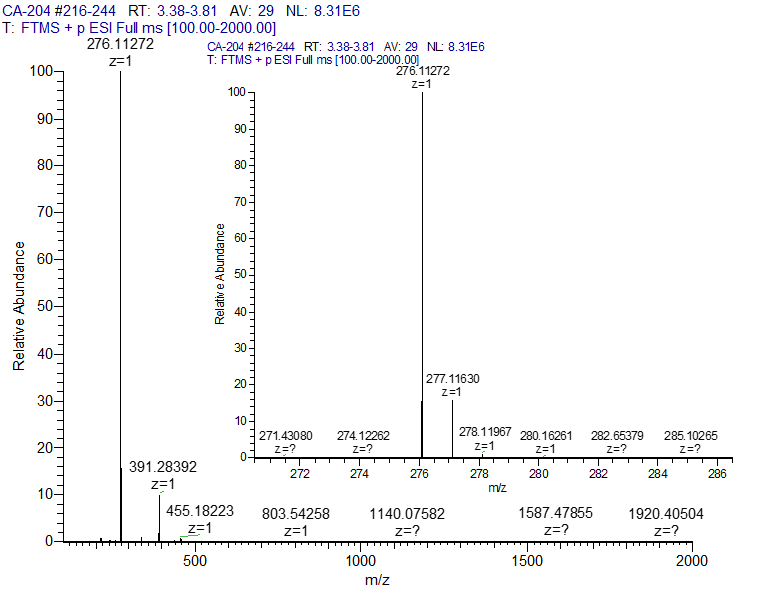

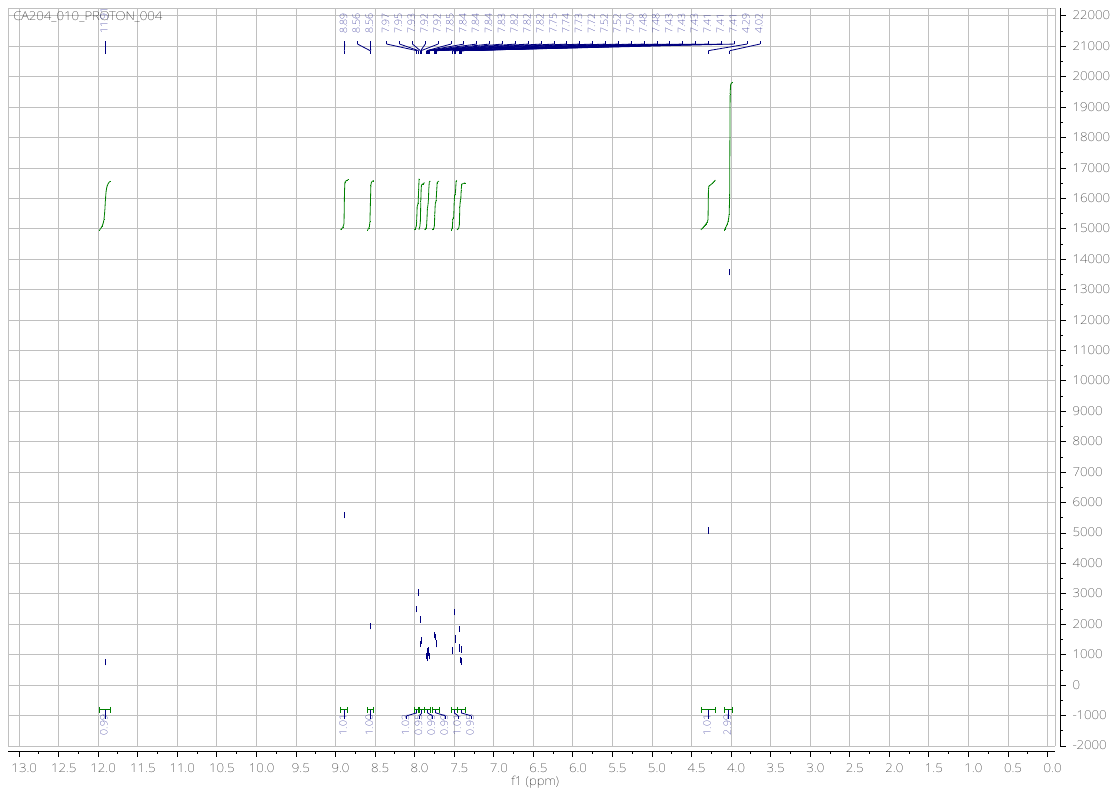

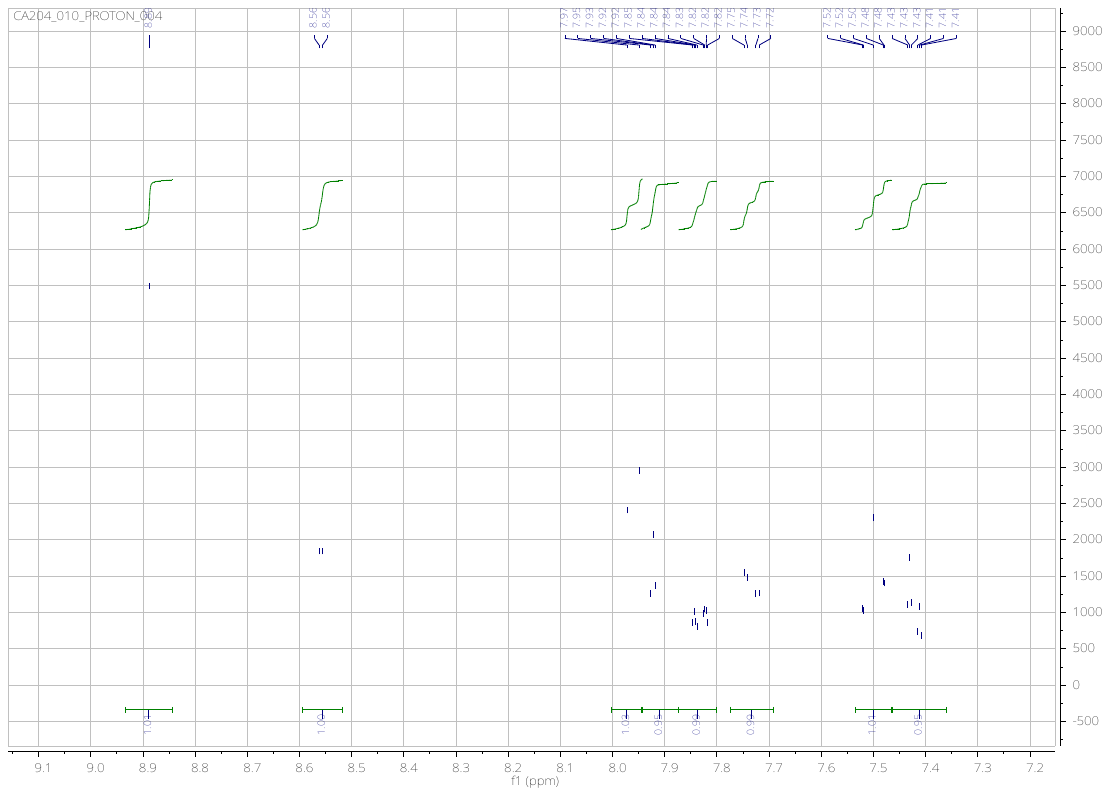

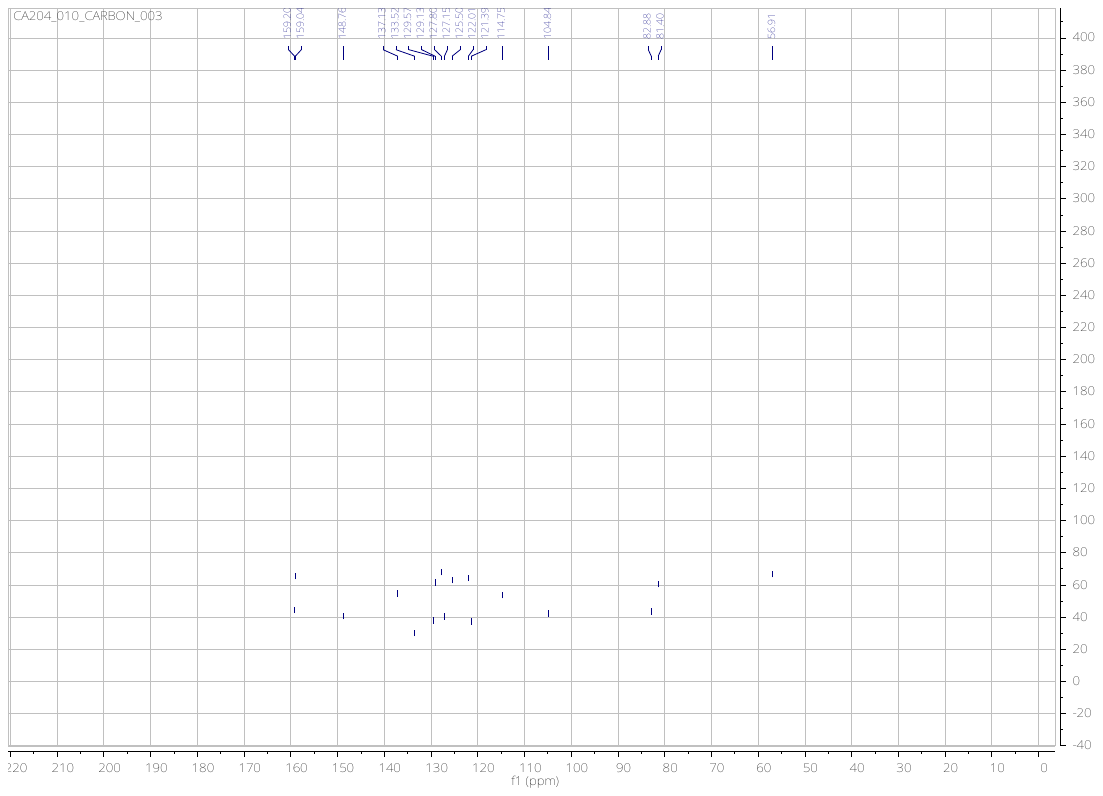

***N*-(3-ethynylphenyl)-7-methoxyquinazolin-4-amine** (**6**)

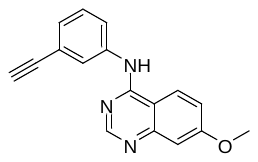

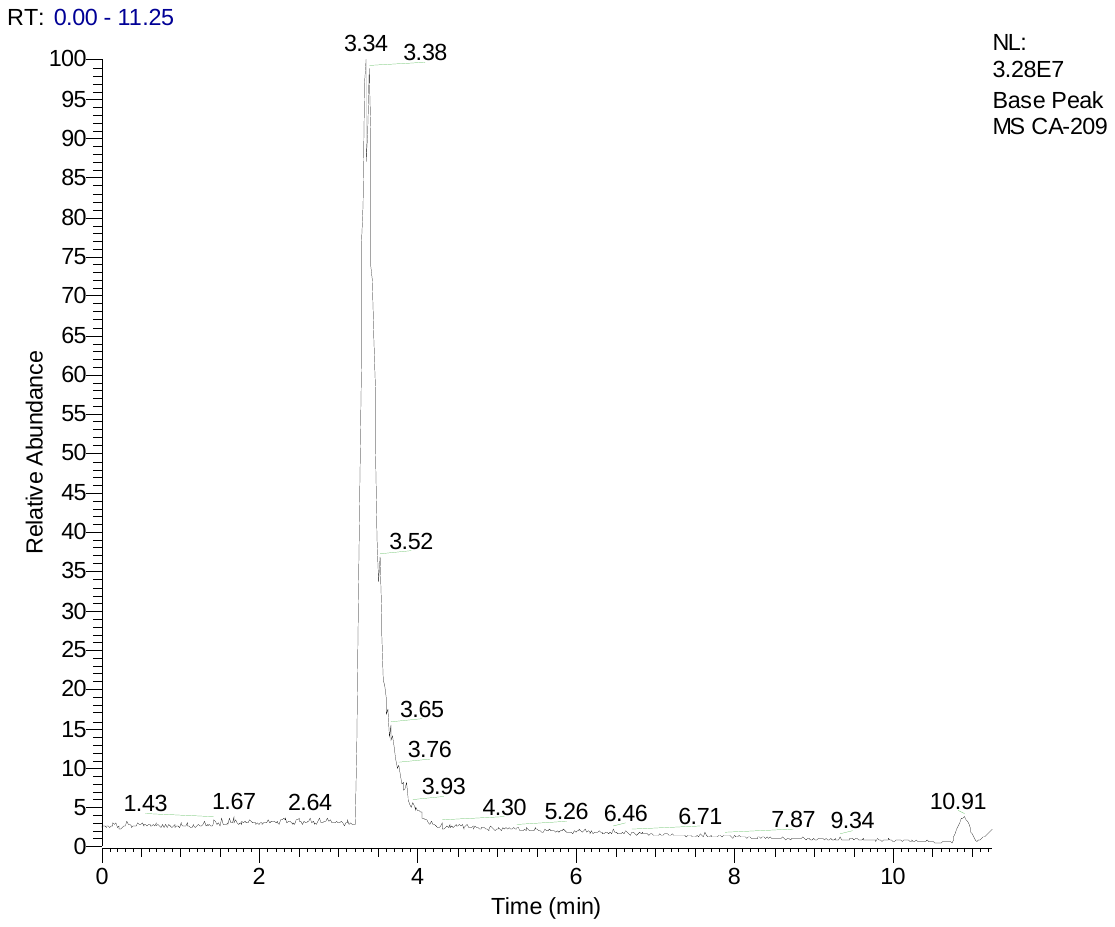

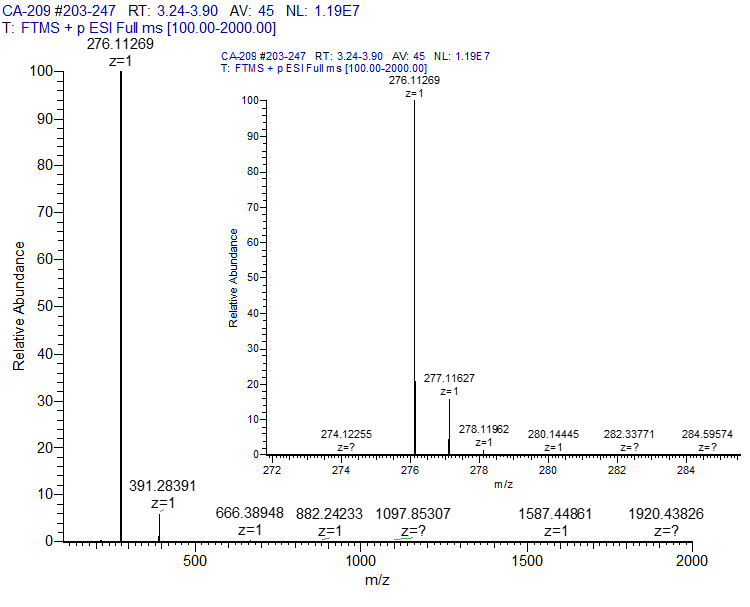

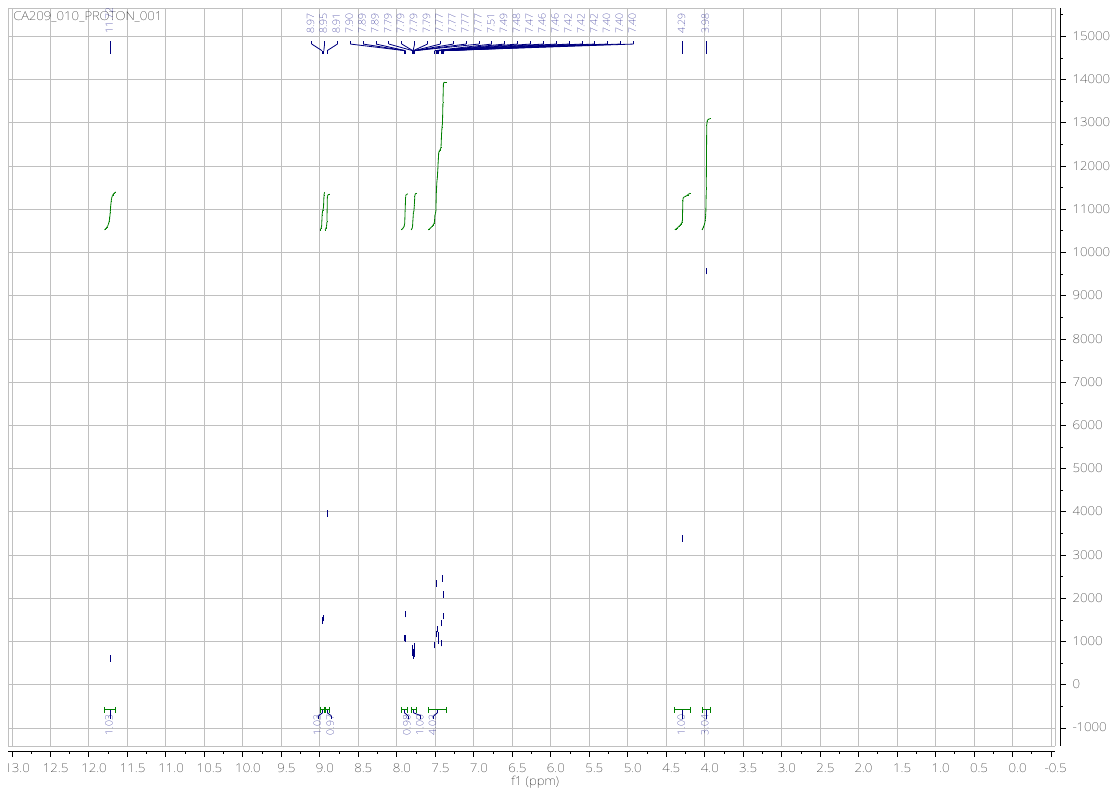

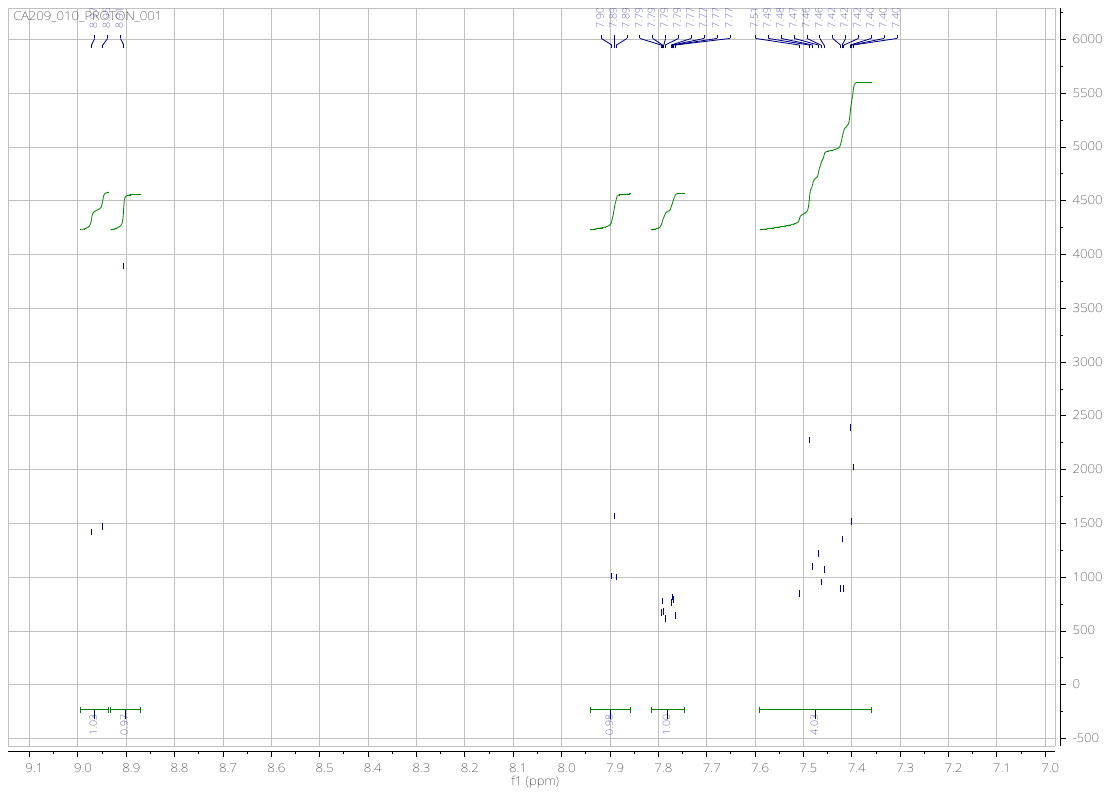

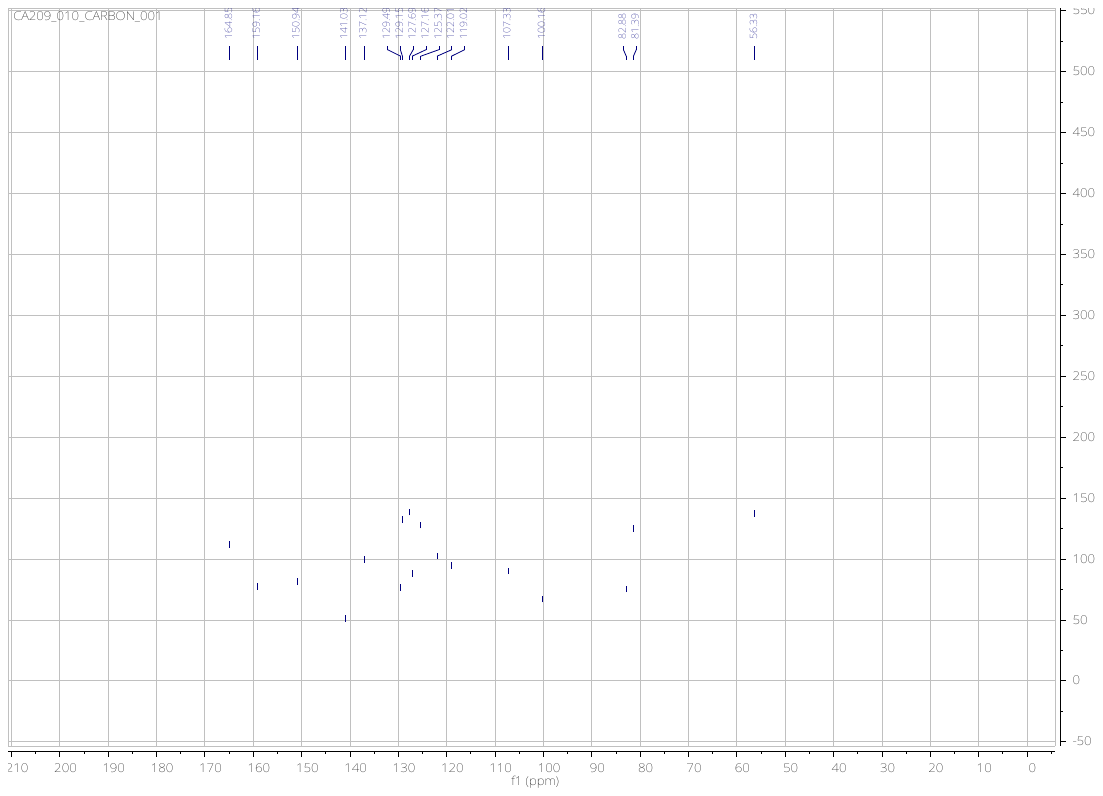

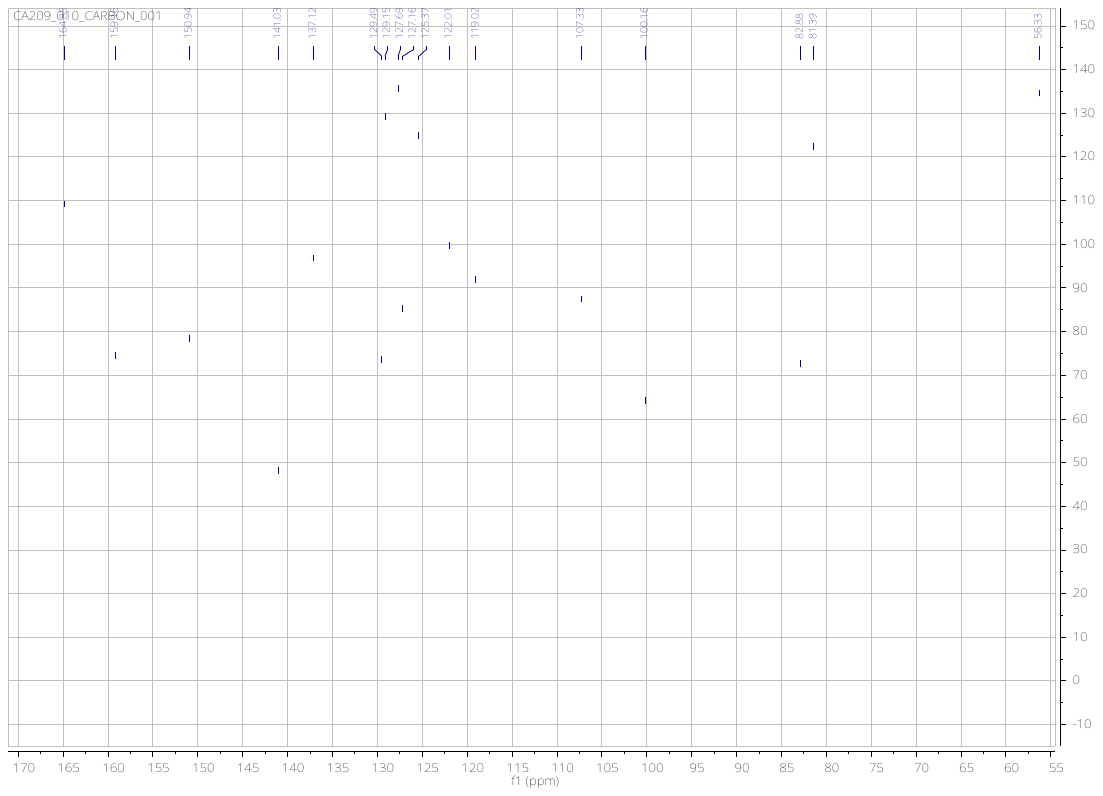

**4-((3-ethynylphenyl)amino)-6,7-dimethoxyquinoline-3-carbonitrile** (**7**)

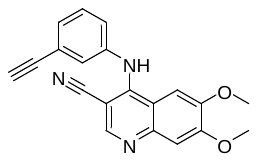

**LC Chromatogram**

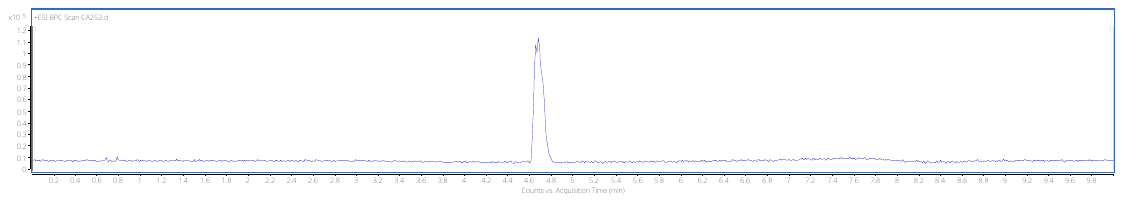

**Full Scan**

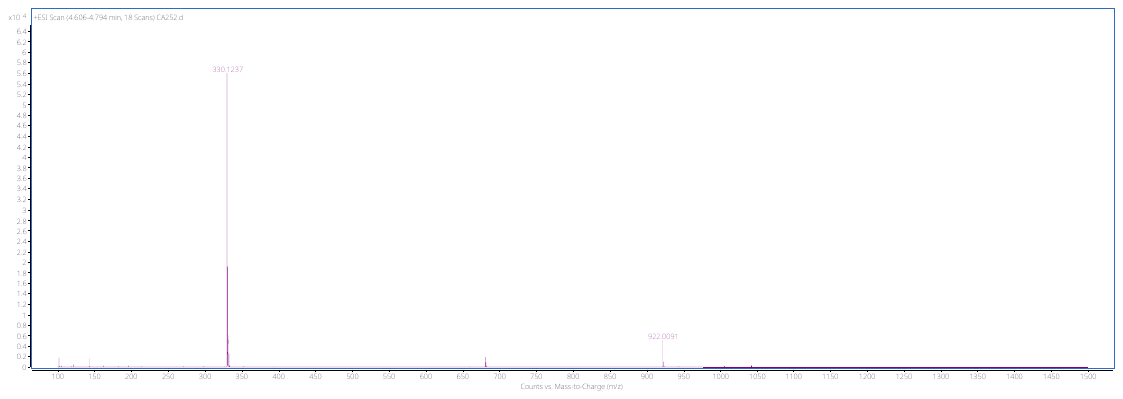

**Full** **Scan at t_r_ = 4.7 min**

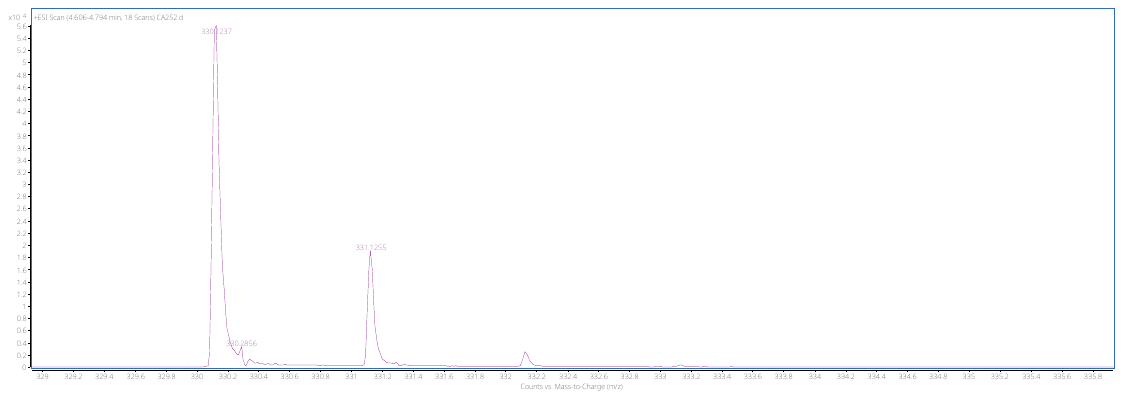

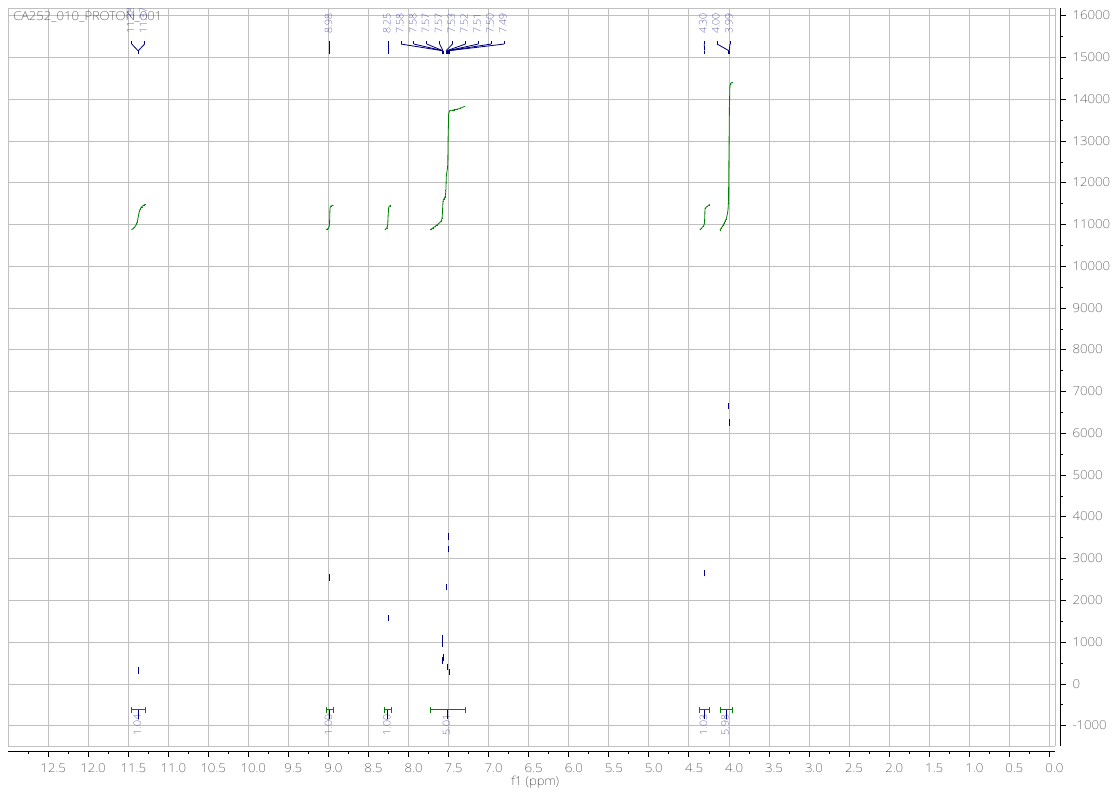

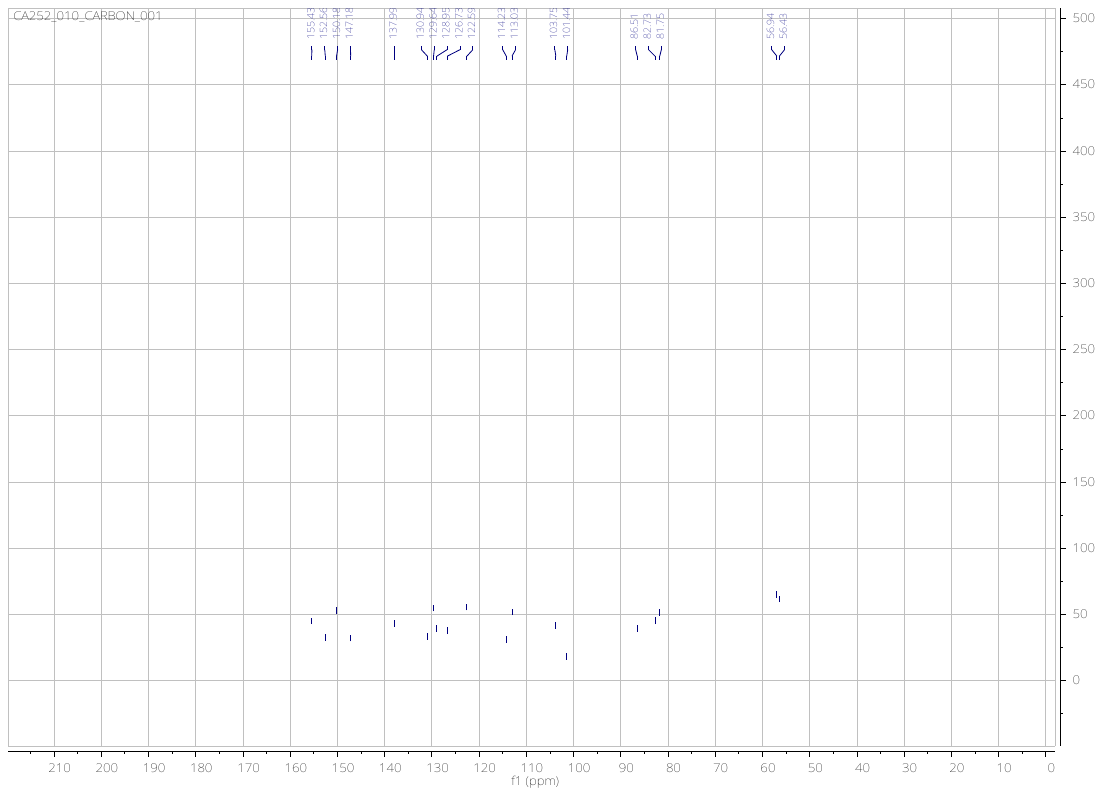

**4-[(3-ethynylphenyl)amino]-6-methoxyquinoline-3-carbonitrile** (**8**)

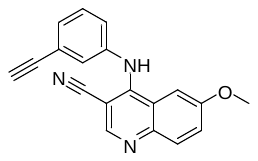

**LC Chromatogram**

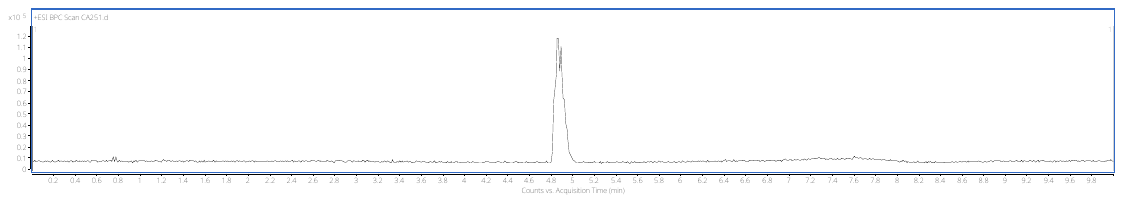

**Full Scan**

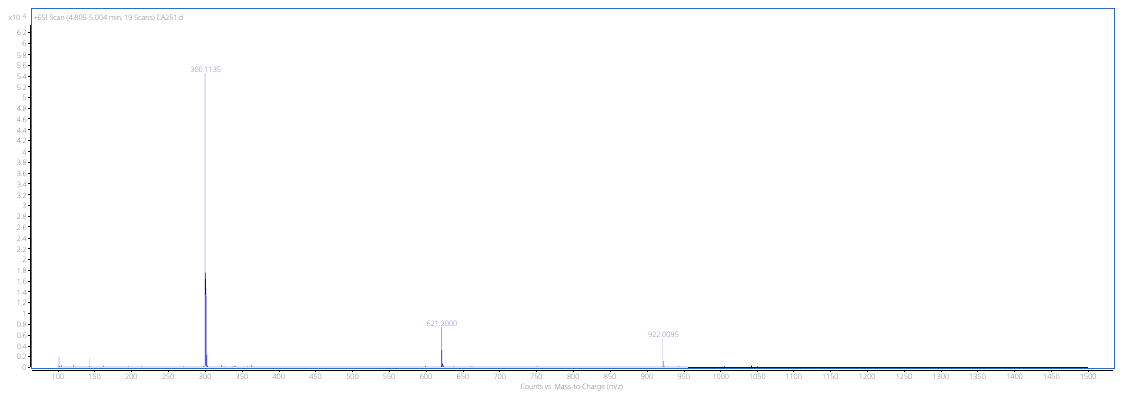

**Full Scan at t_r_ = 4.9 min**

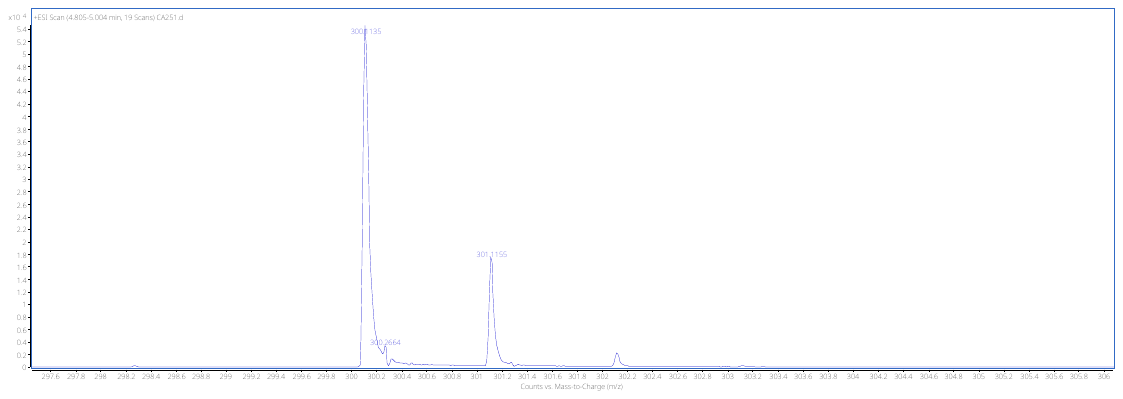

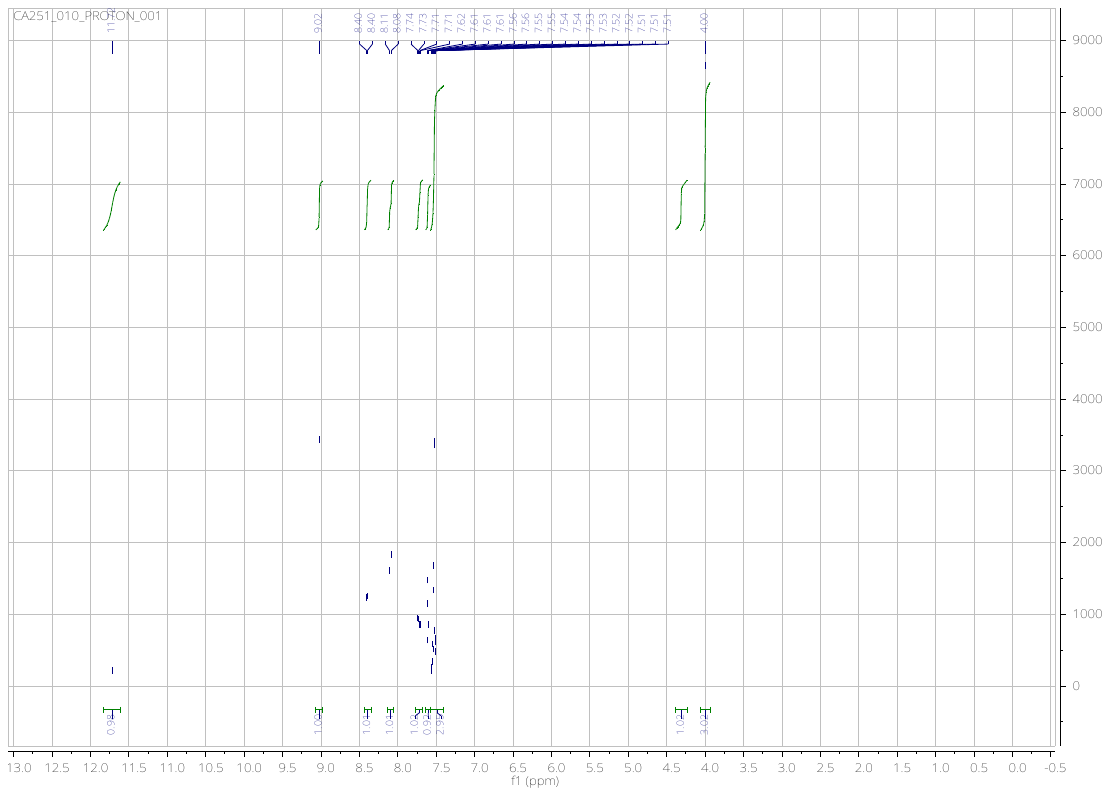

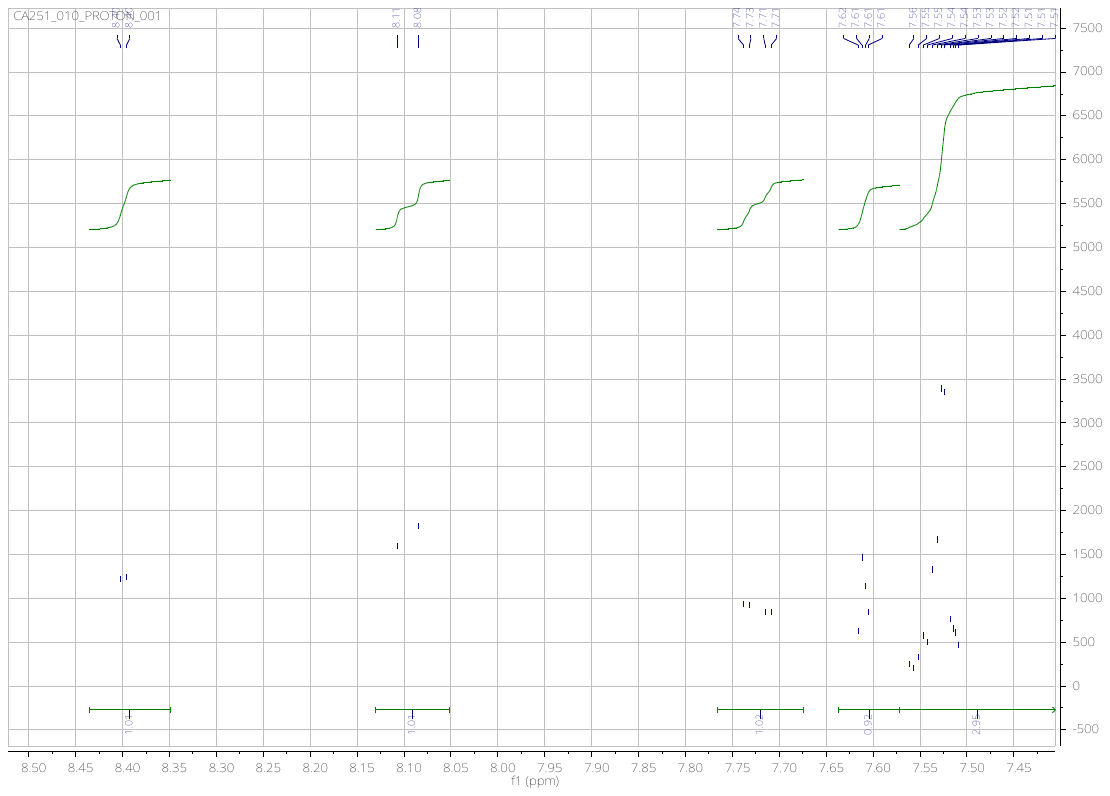

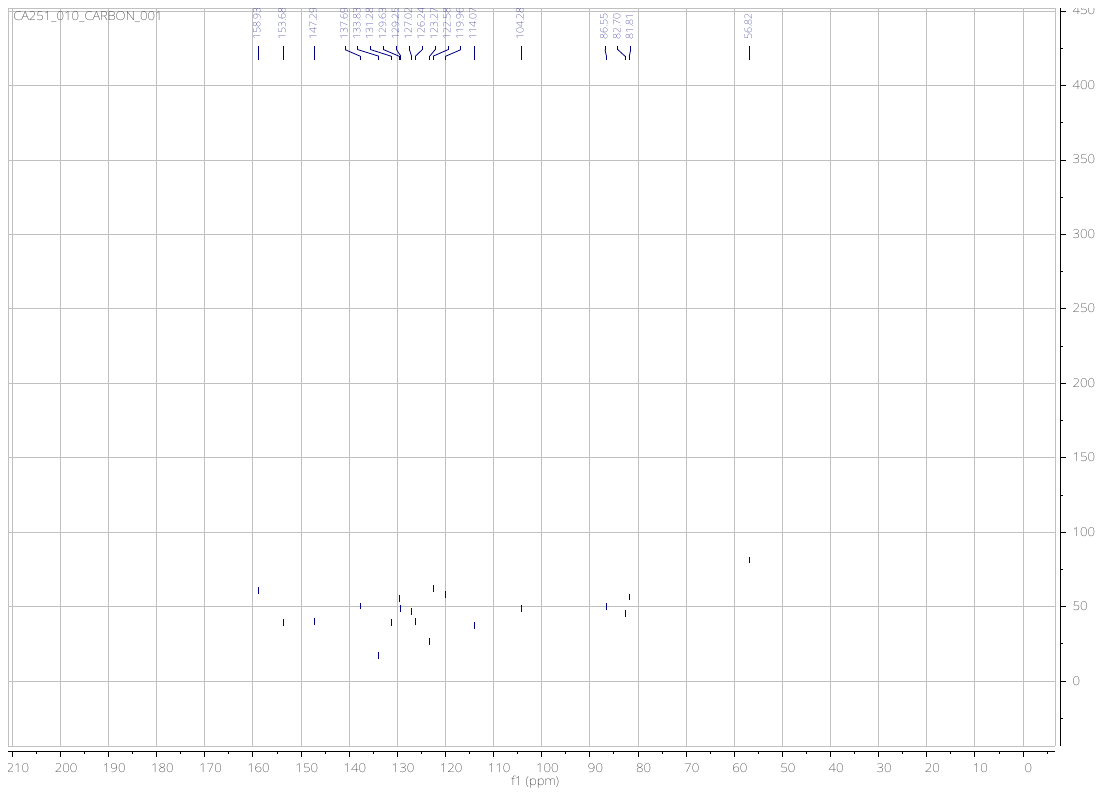

**4-[(3-ethynylphenyl)amino]-7-methoxyquinoline-3-carbonitrile** - **(9)**

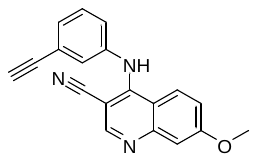

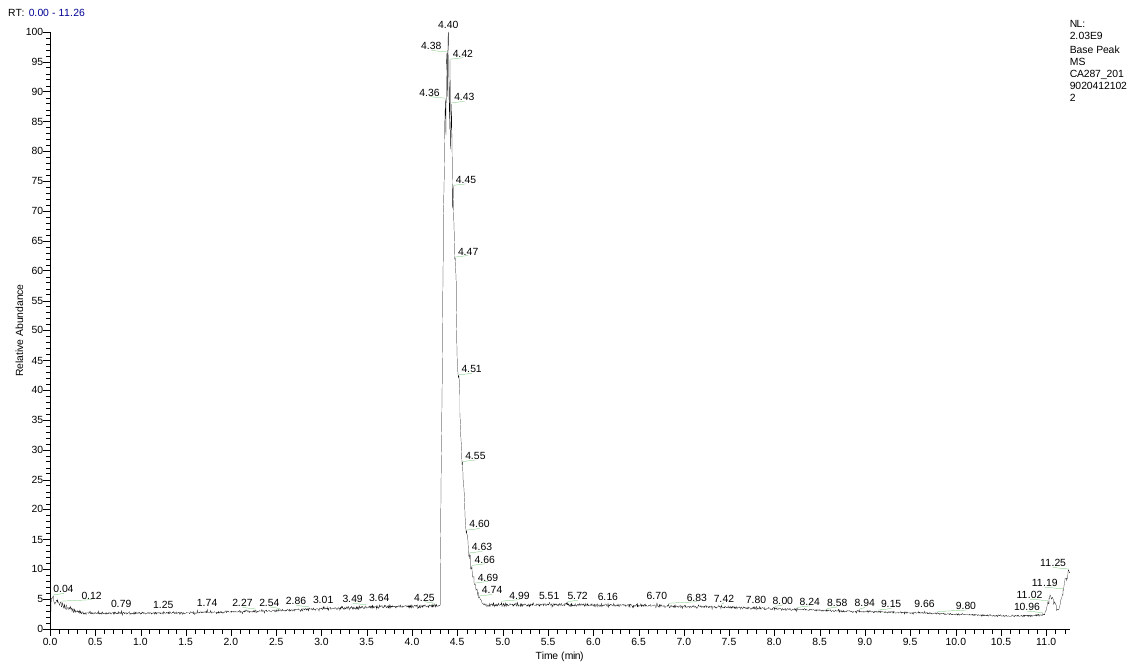

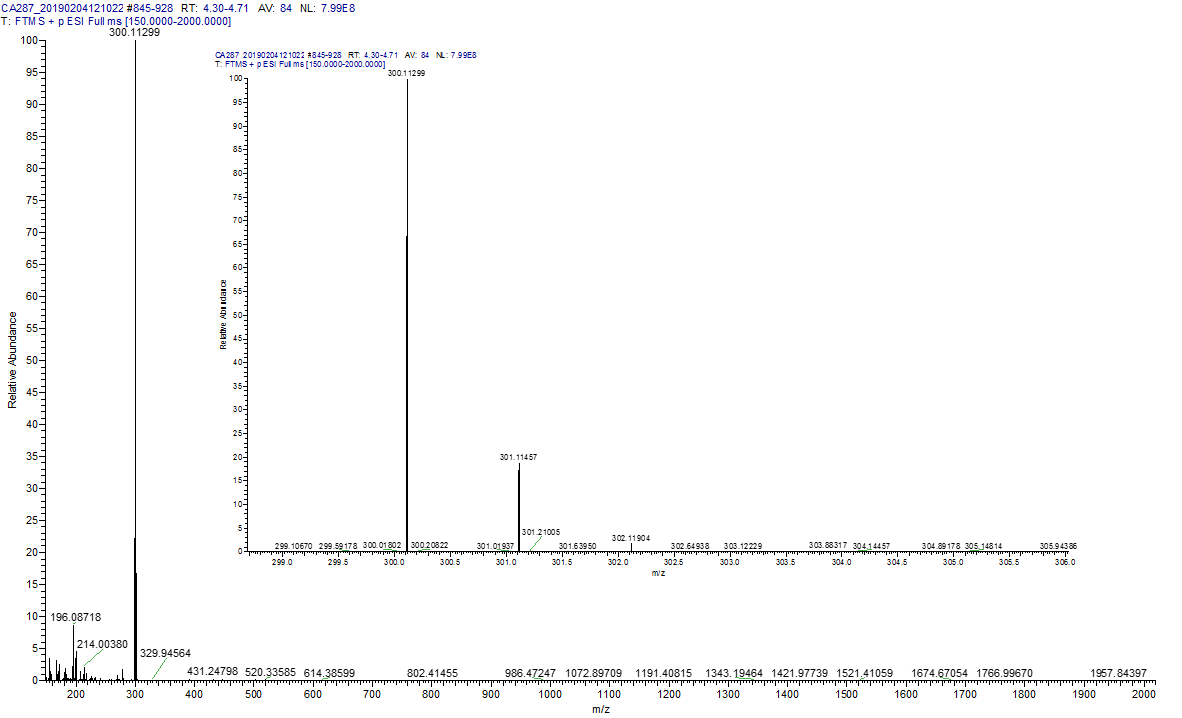

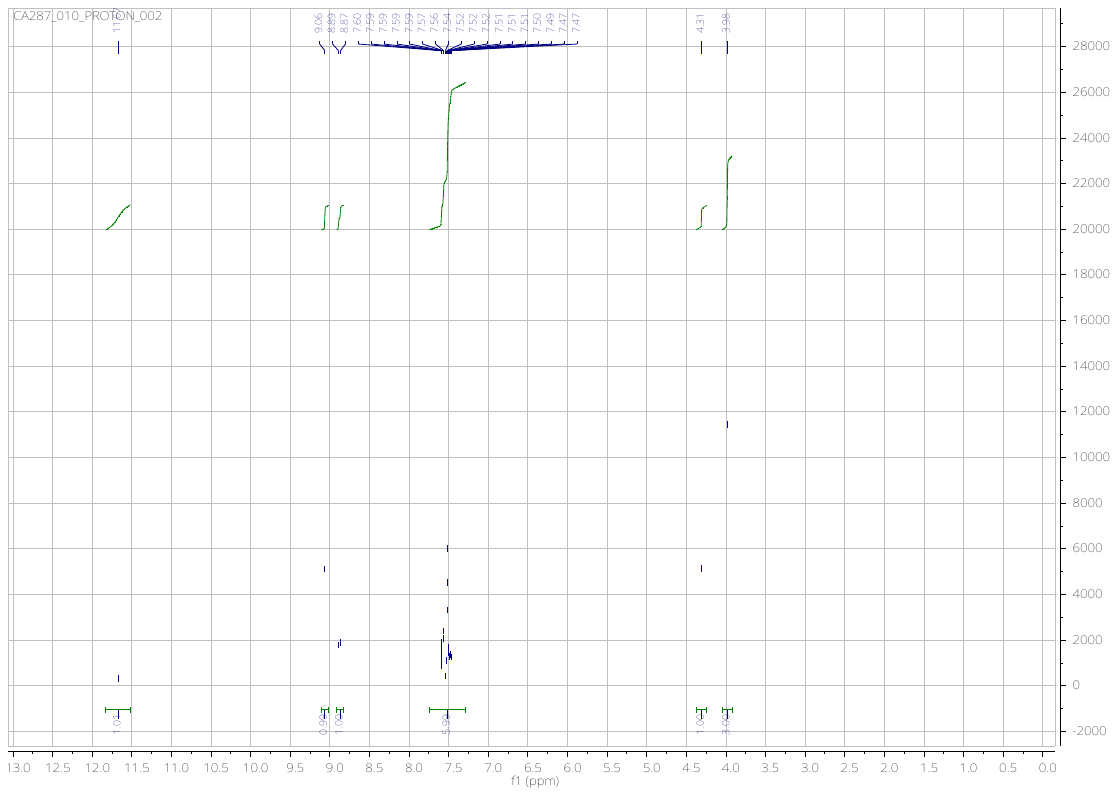

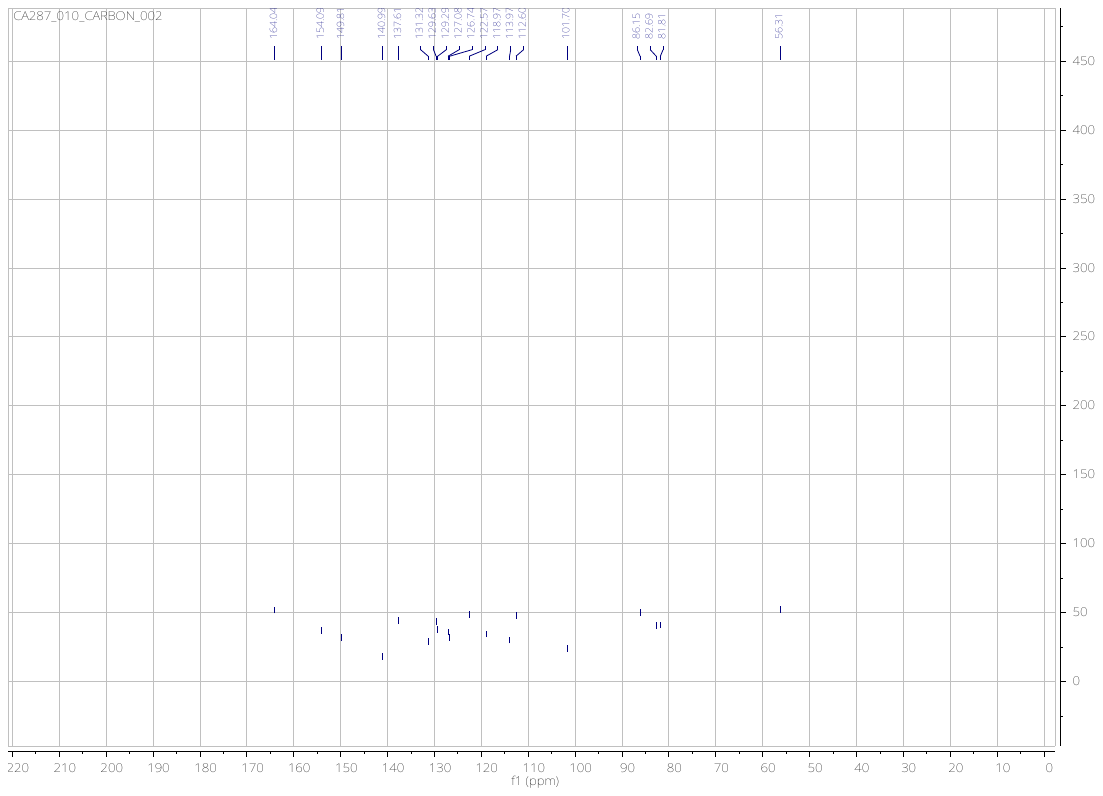

***N*-{4-[(6,7-dimethoxyquinolin-4-yl)amino]-2-hydroxyphenyl}-4-methylbenzamide** (**13**)

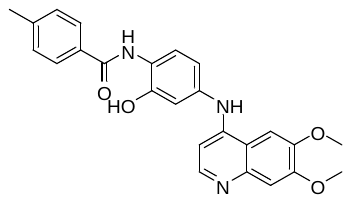

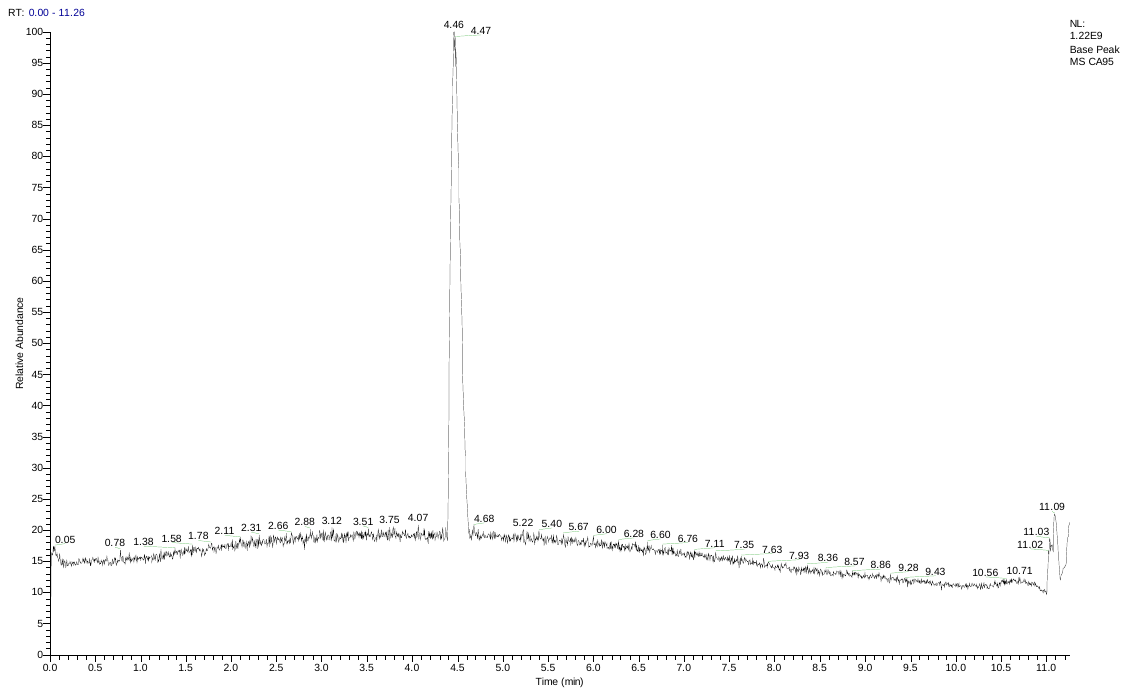

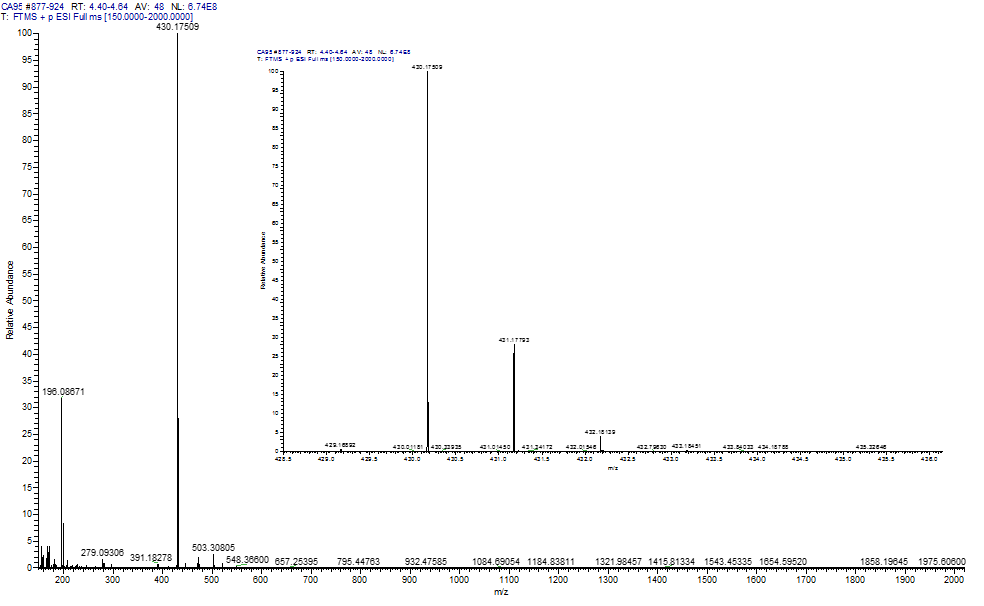

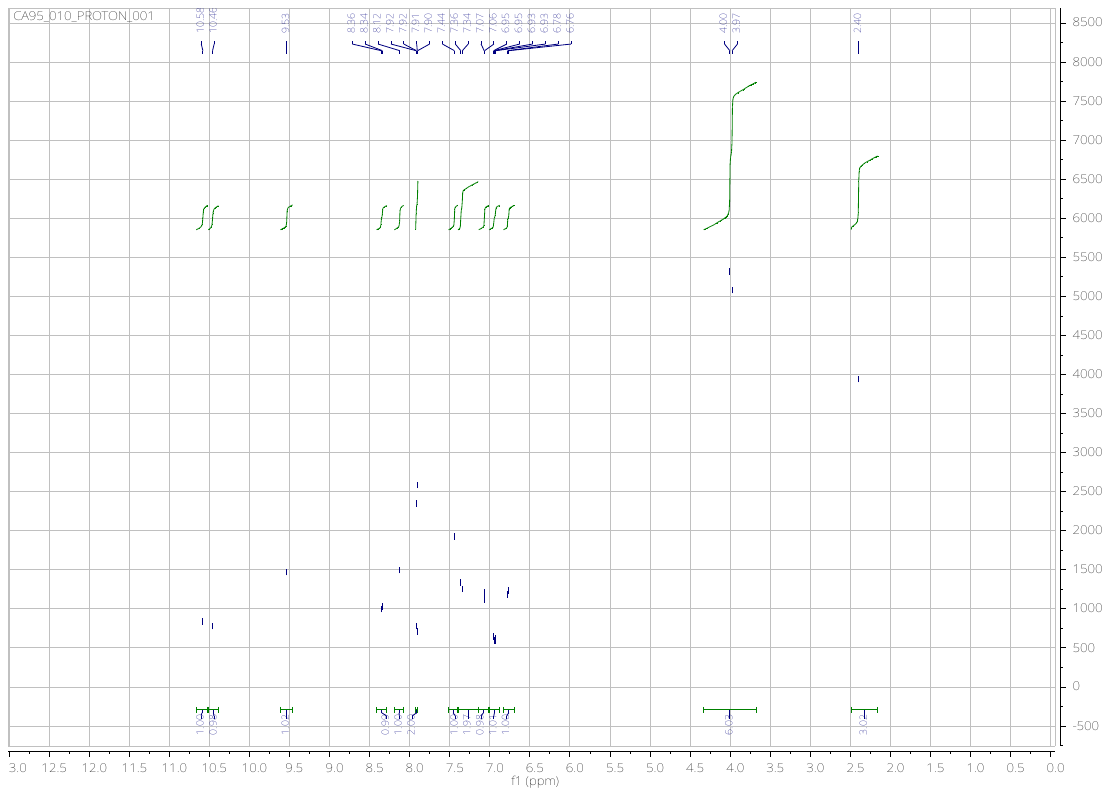

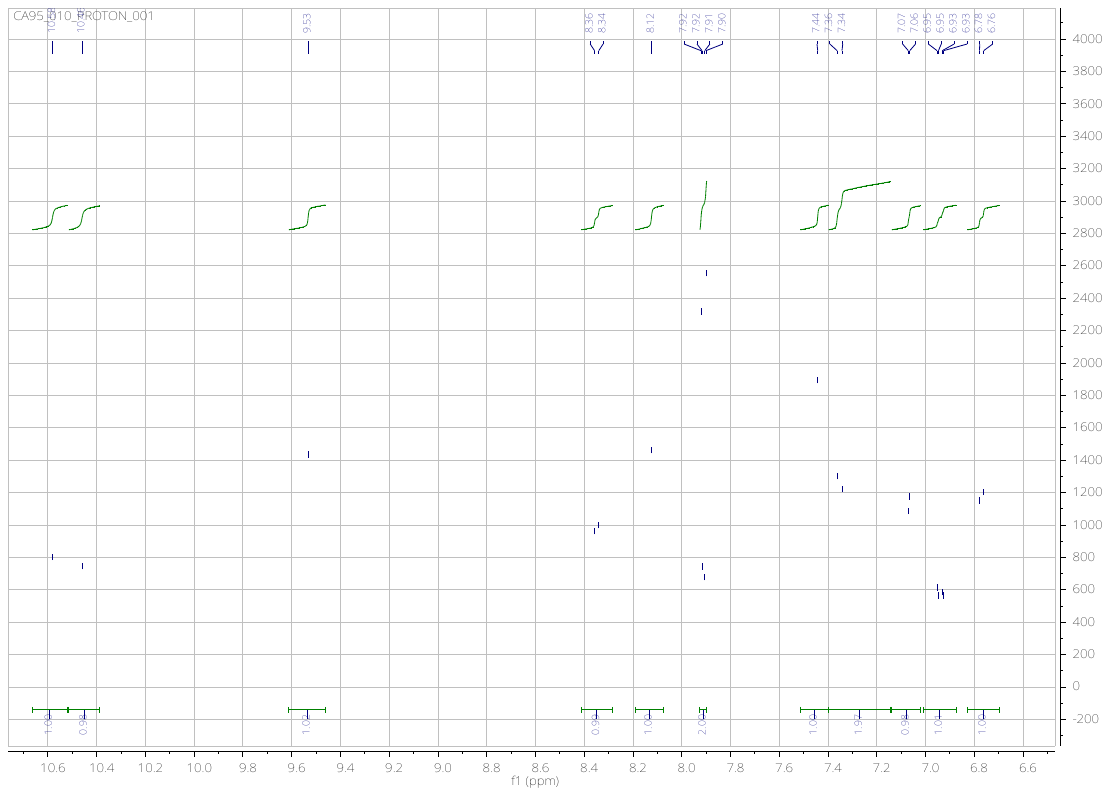

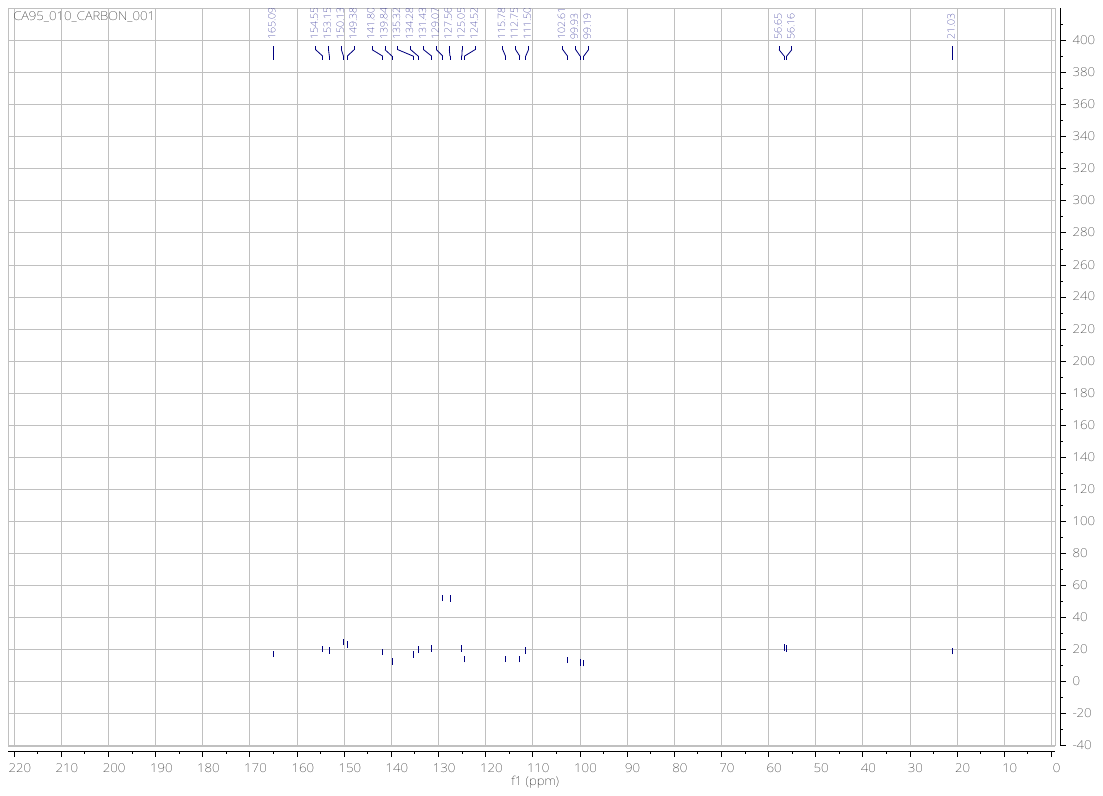

***N*-(2-hydroxy-4-((6-methoxyquinolin-4-yl)amino)phenyl)-4-methylbenzamide** (**14**)

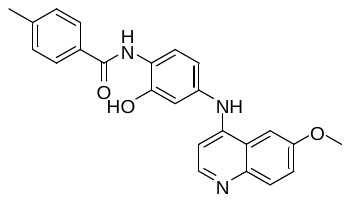

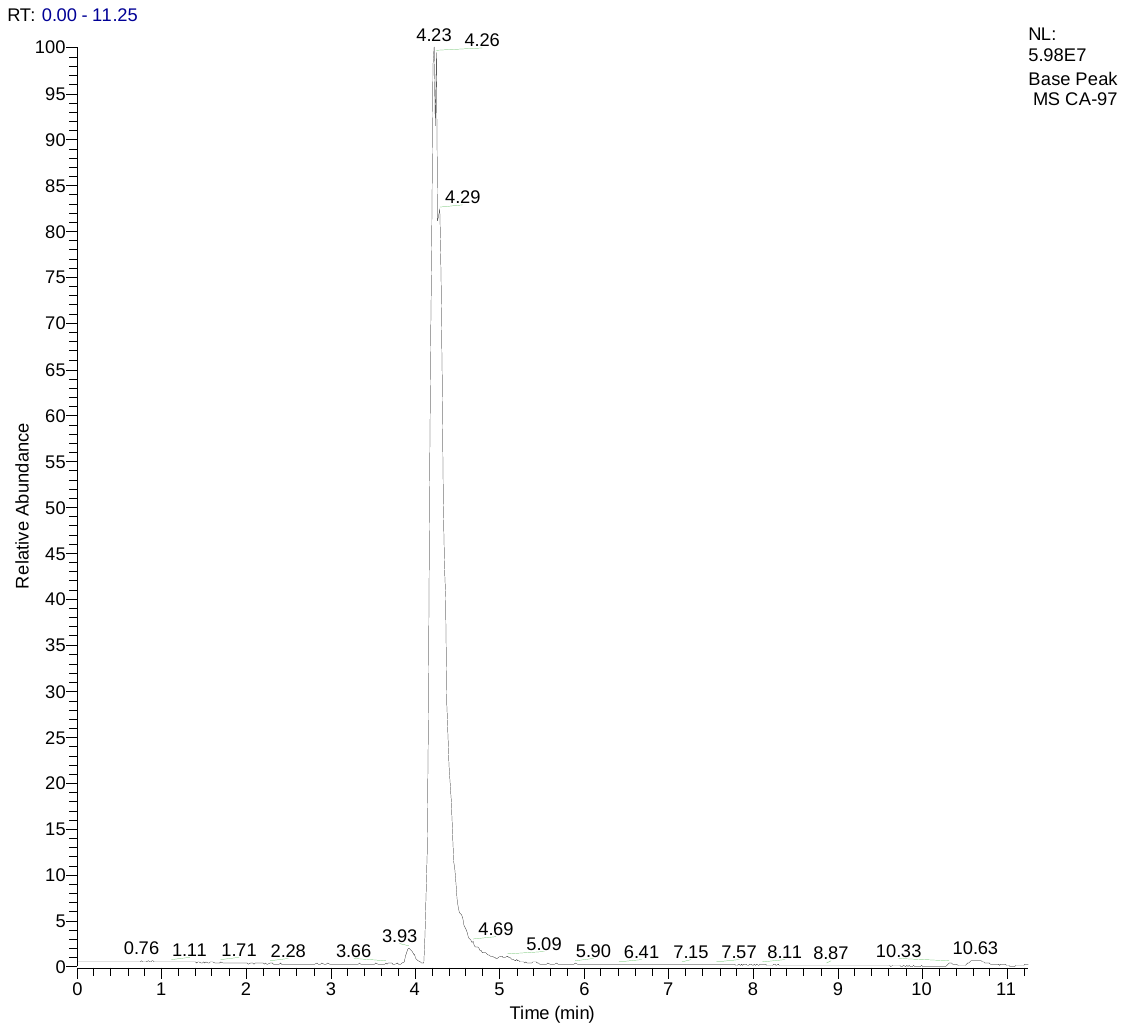

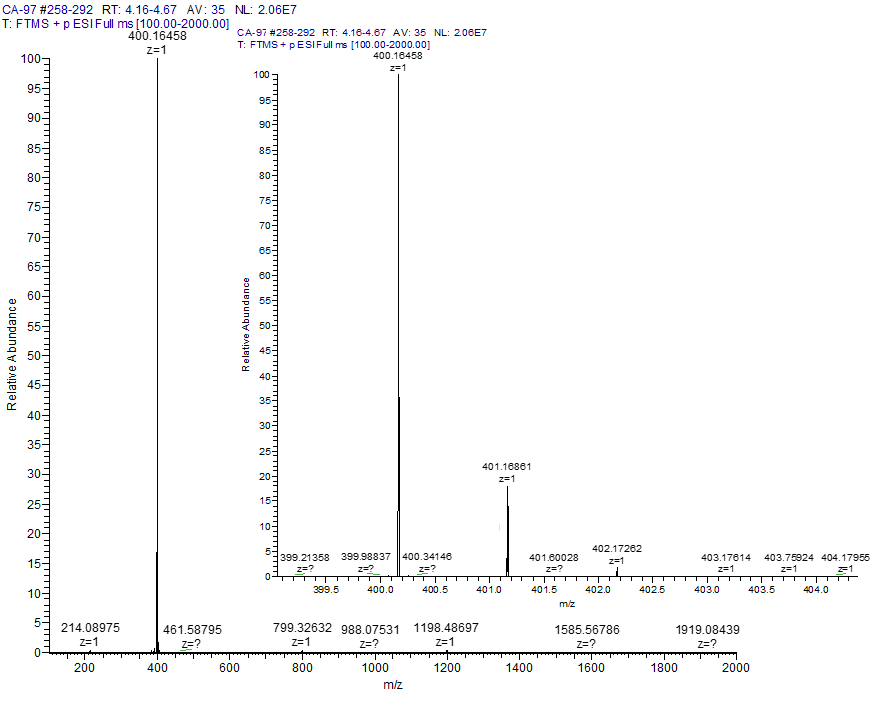

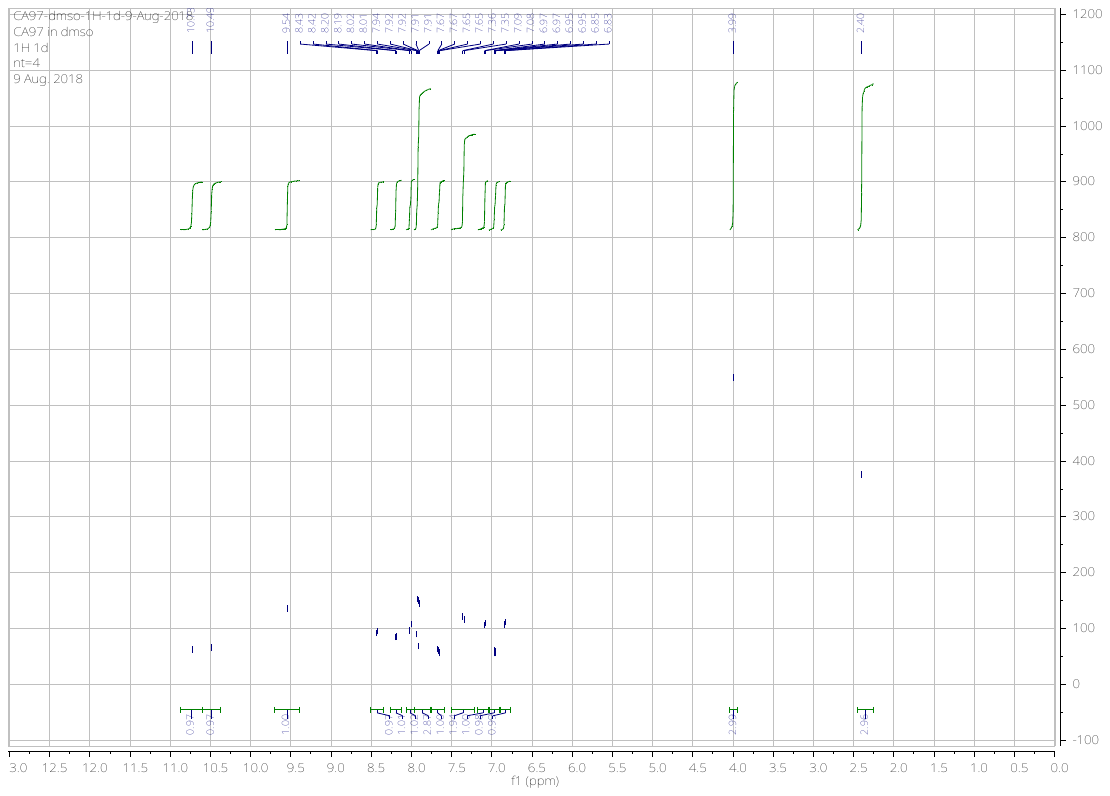

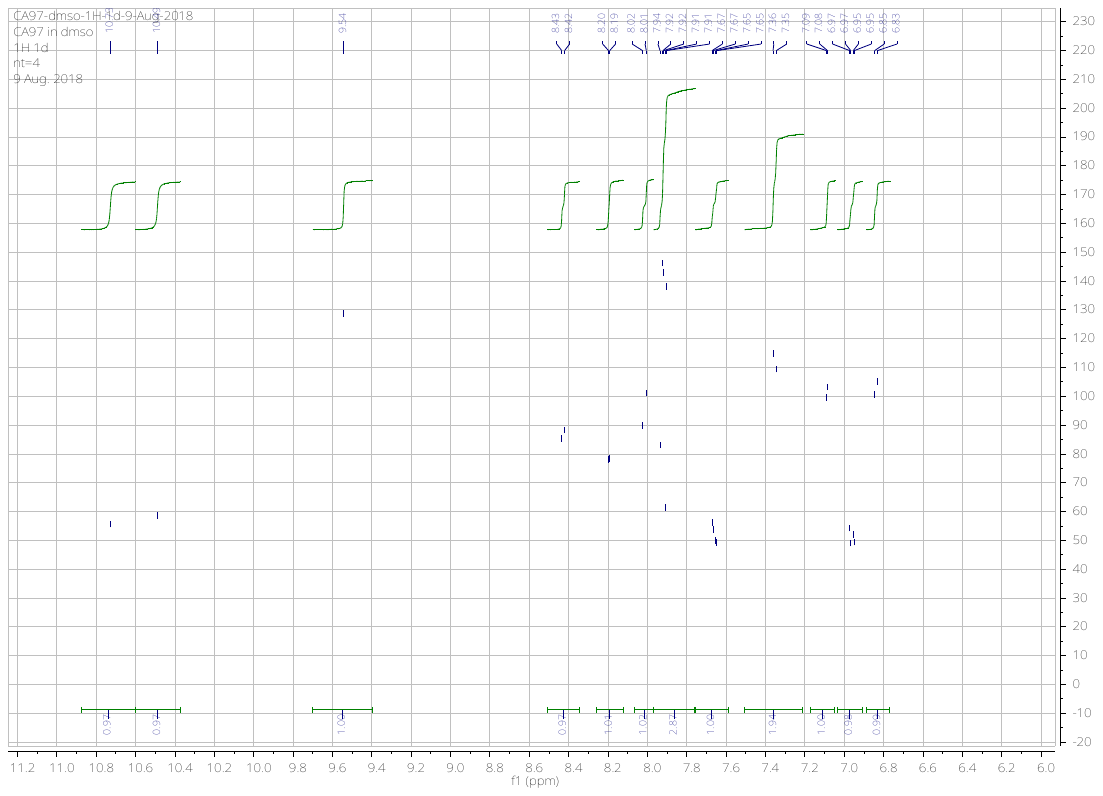

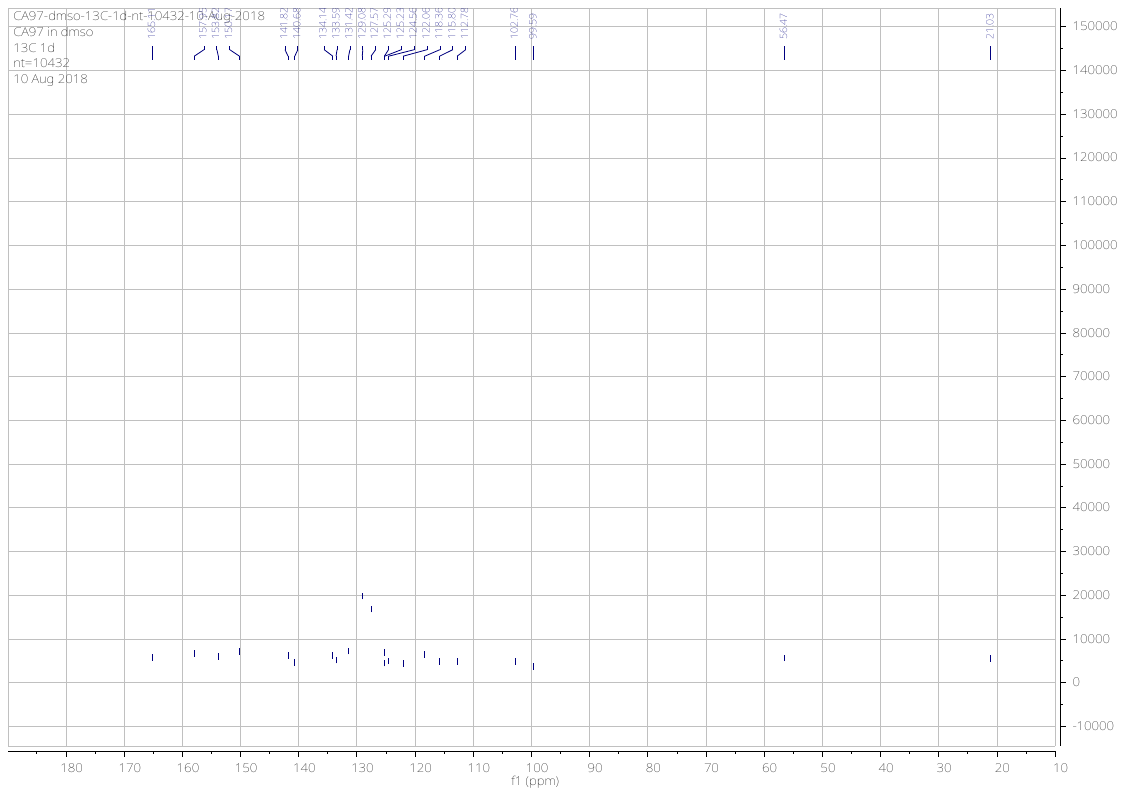

***N*-(2-hydroxy-4-((7-methoxyquinolin-4-yl)amino)phenyl)-4-methylbenzamide** (**15**)

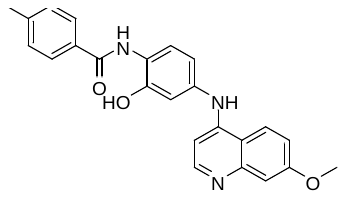

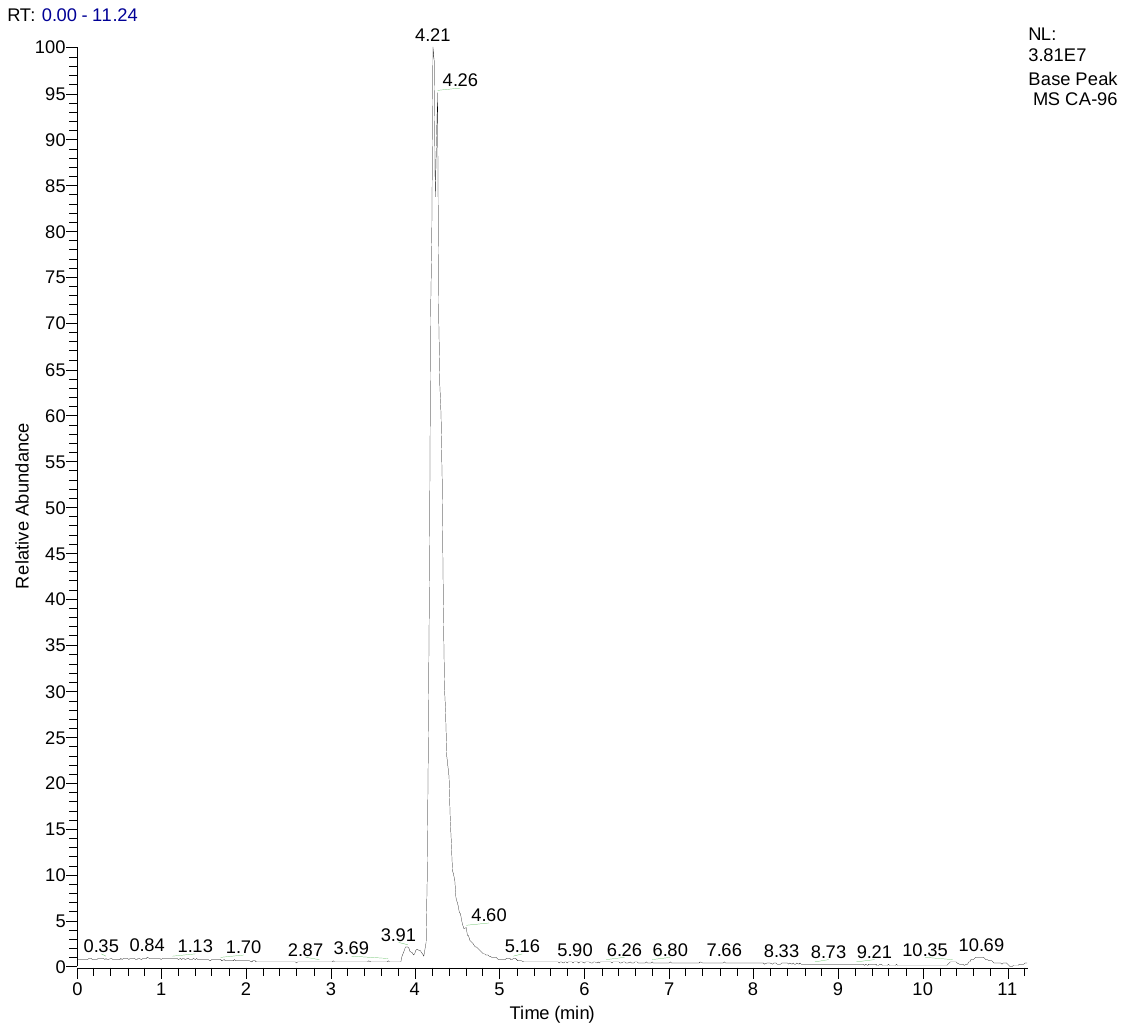

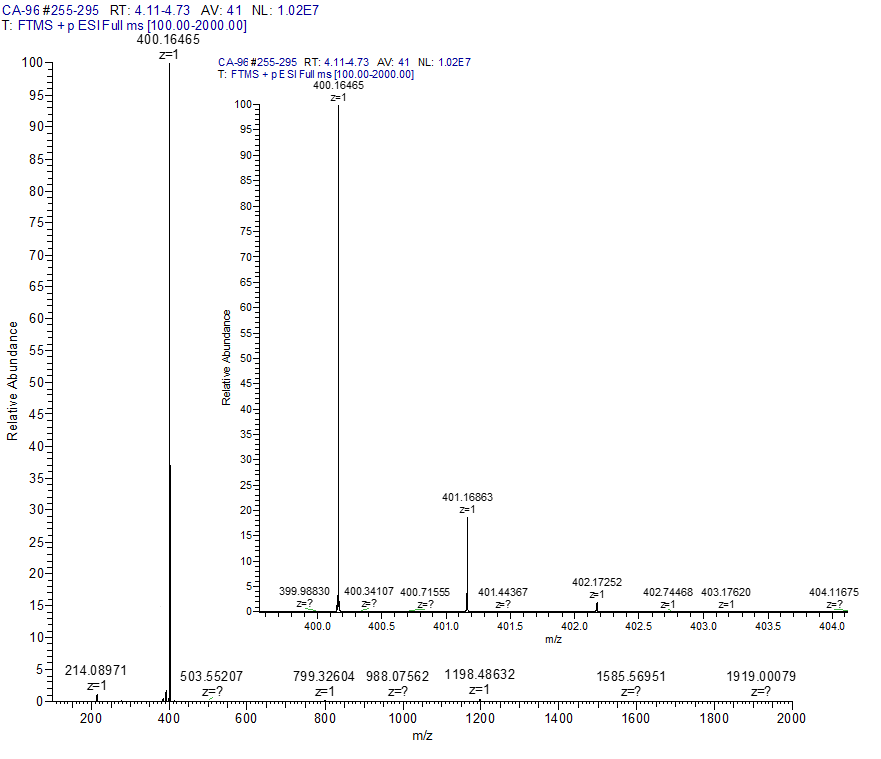

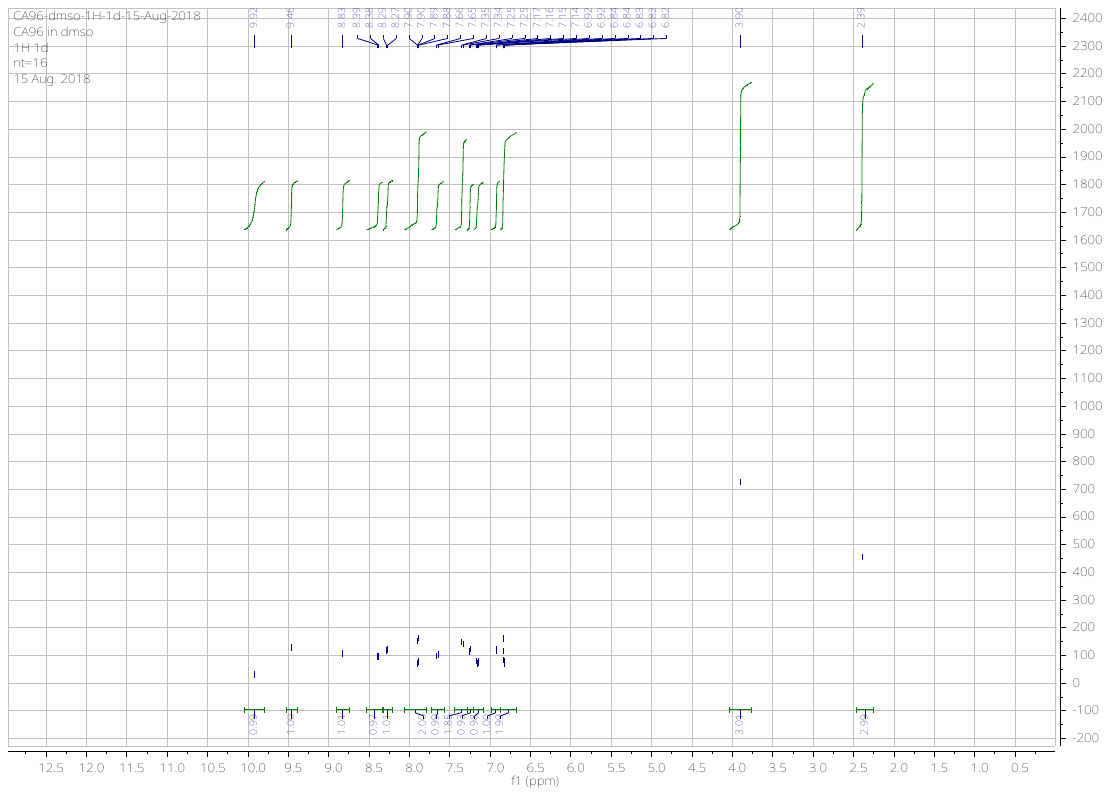

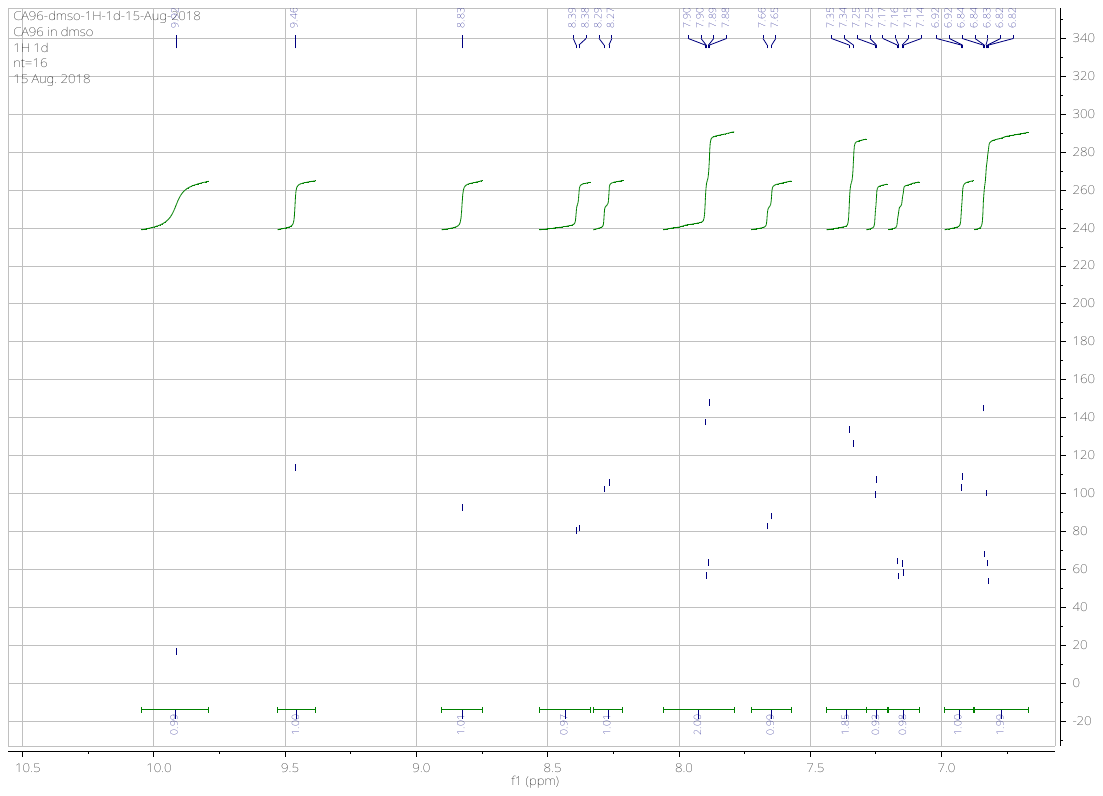

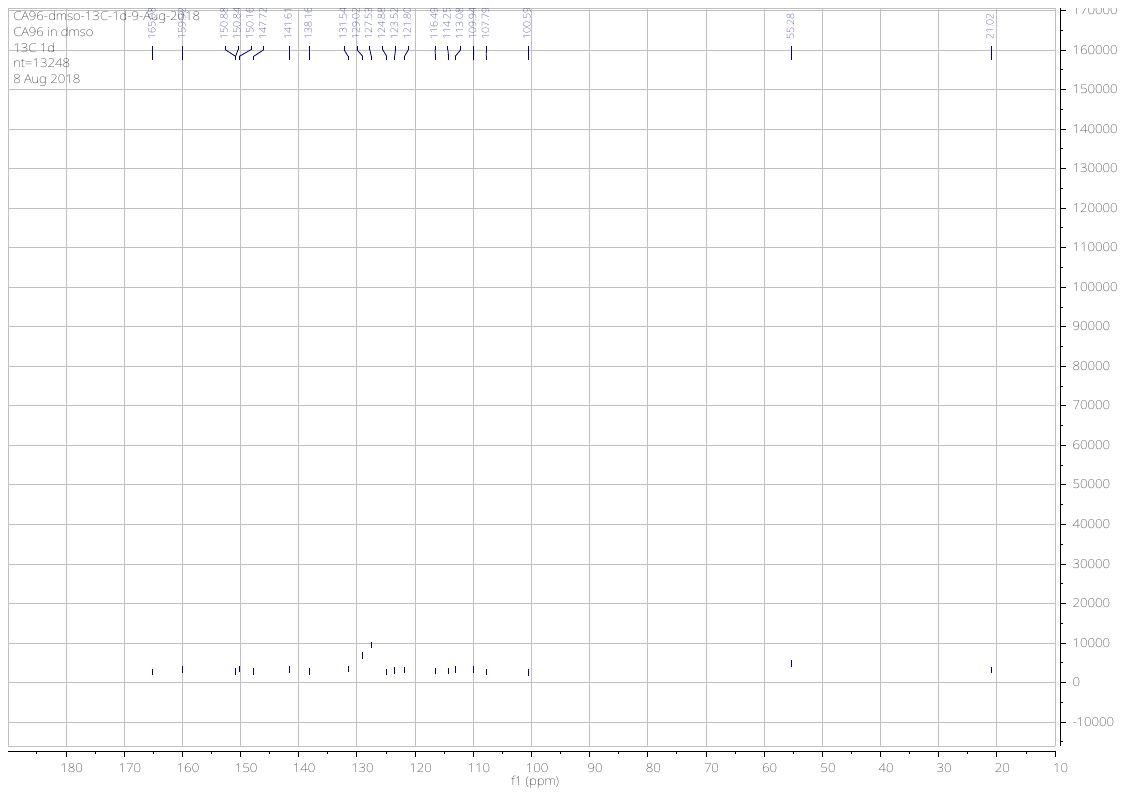

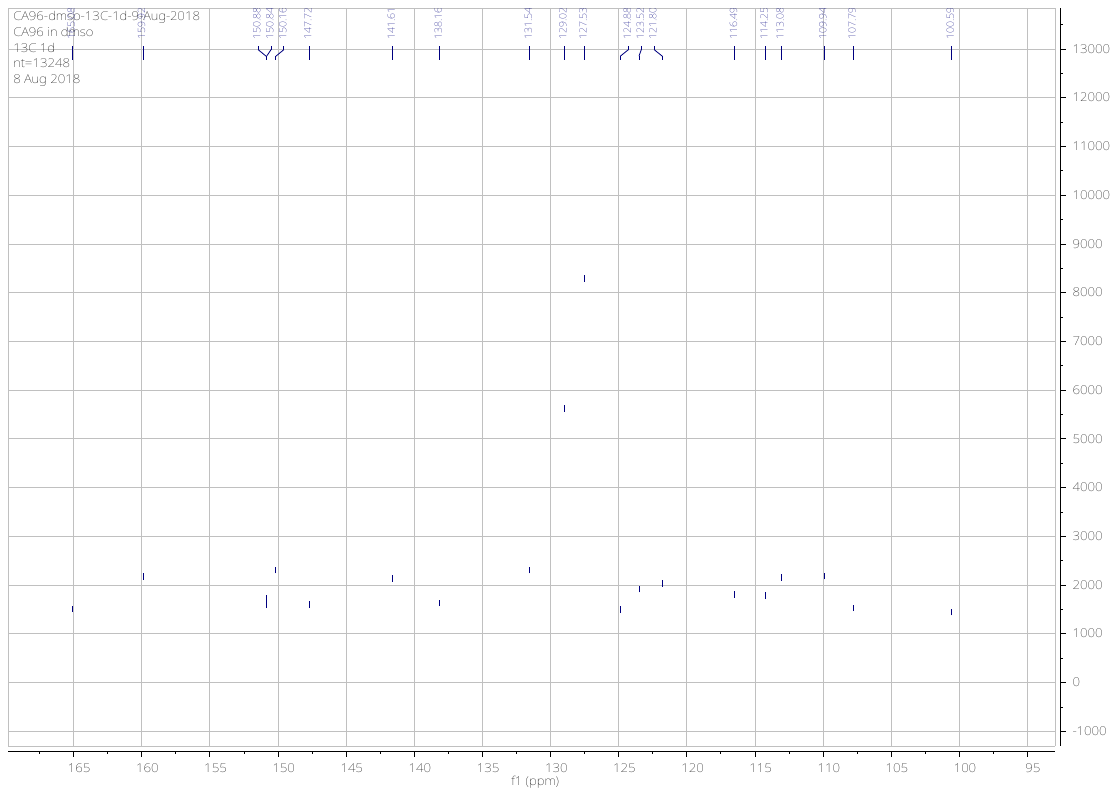

***N*-(4-((6,7-dimethoxyquinolin-4-yl)amino)phenyl)-4-methylbenzamide** (**16**)

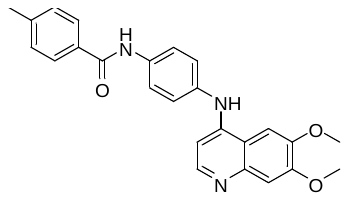

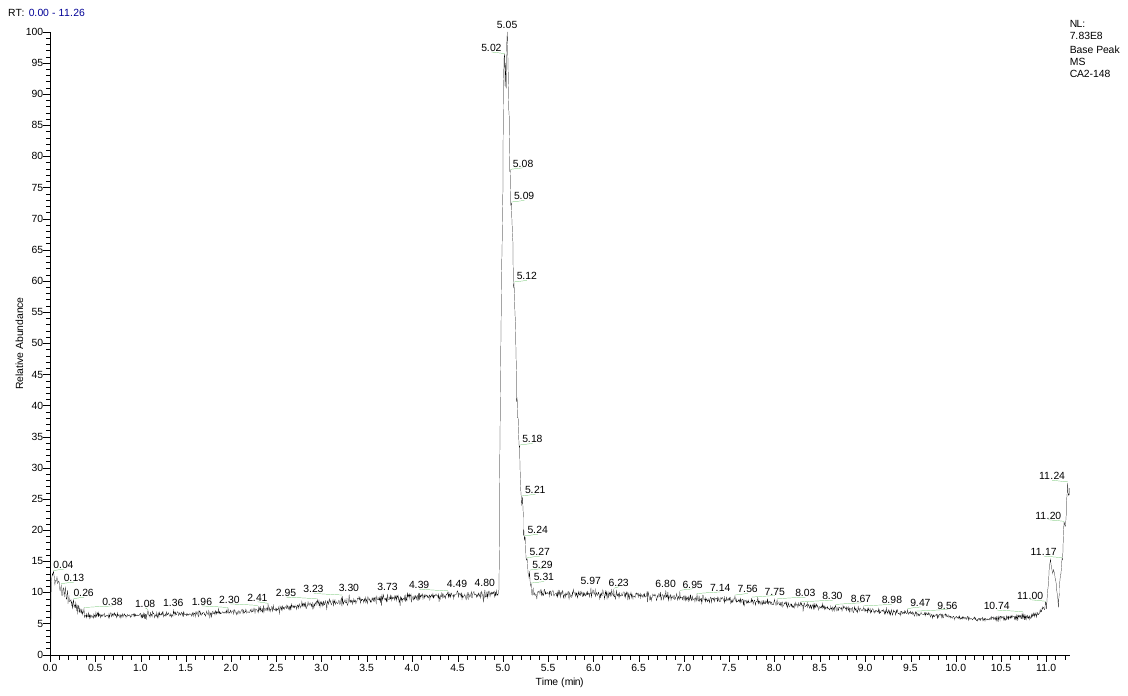

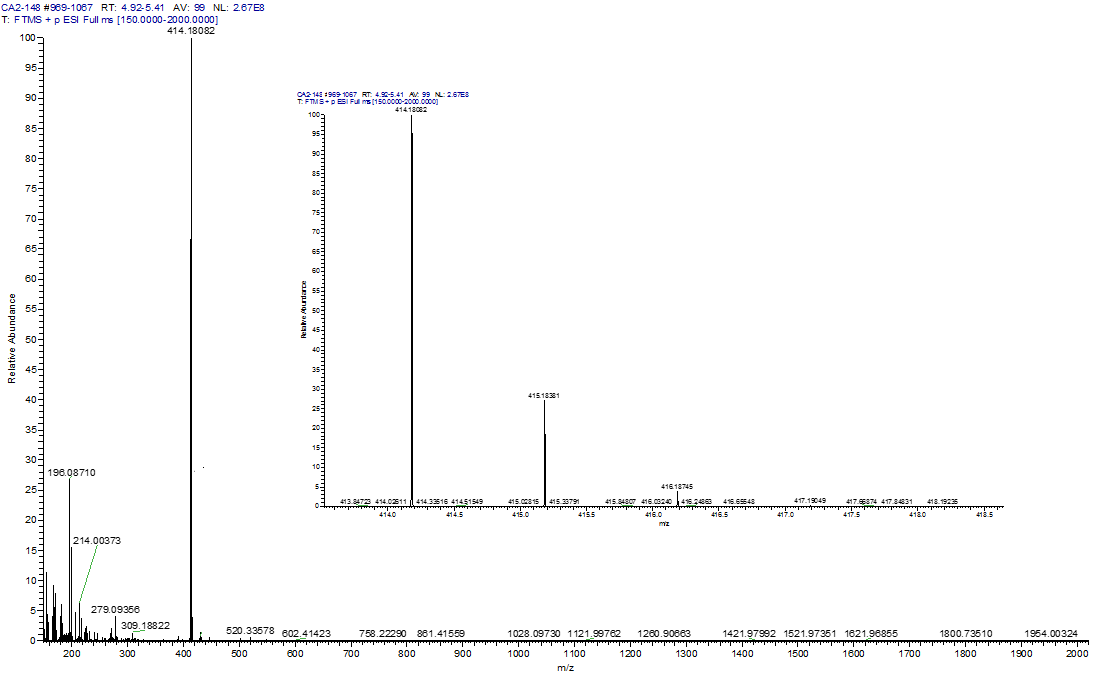

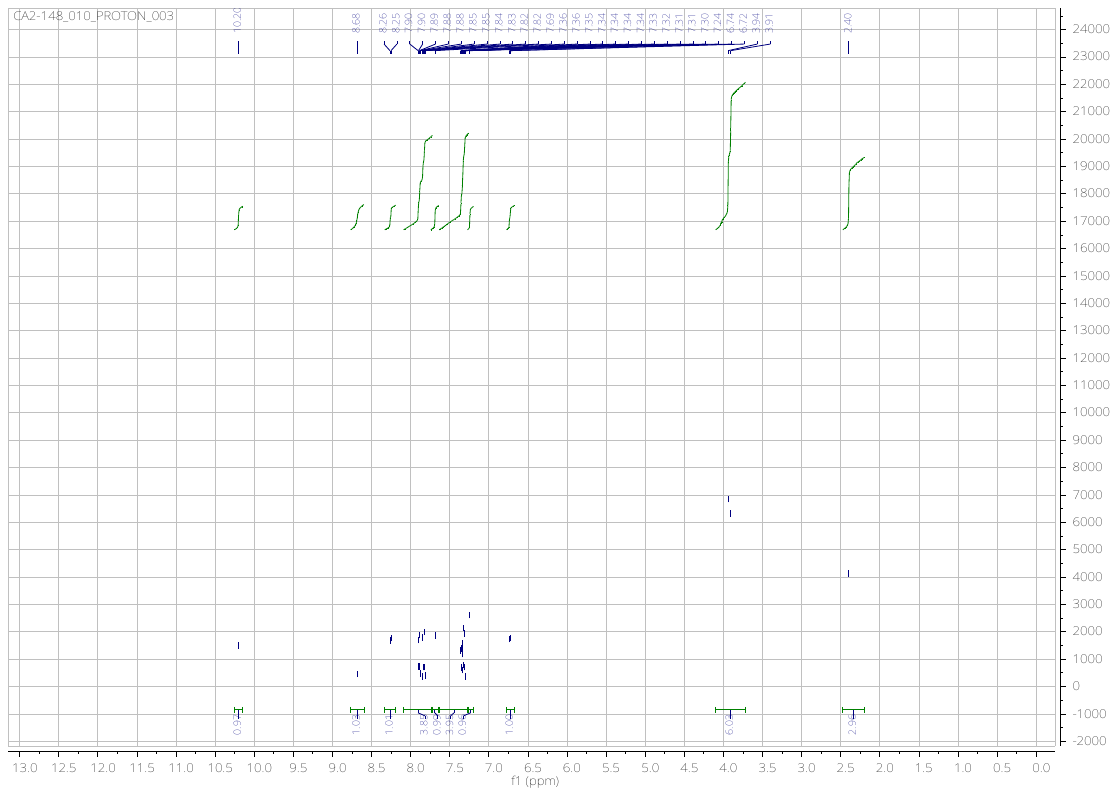

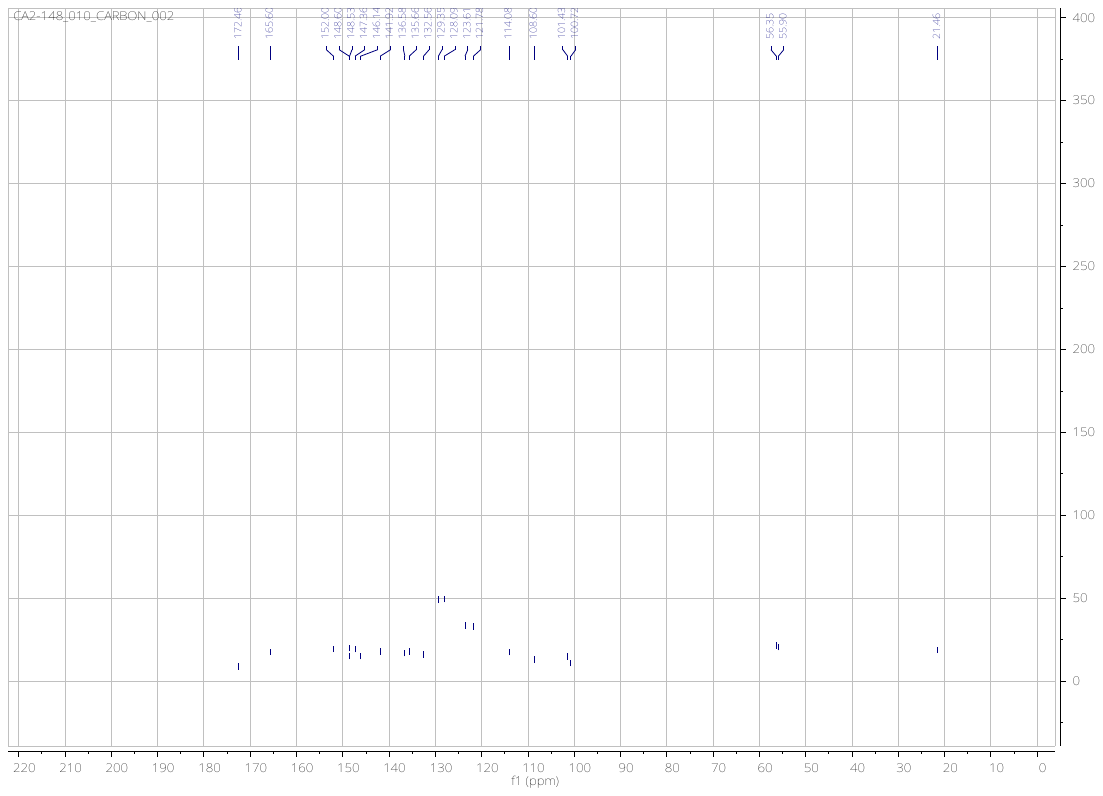

**4-(*tert*-butyl)-*N*-(4-((6,7-dimethoxyquinolin-4-yl)amino)-2-hydroxyphenyl)benzamide** (**17**)

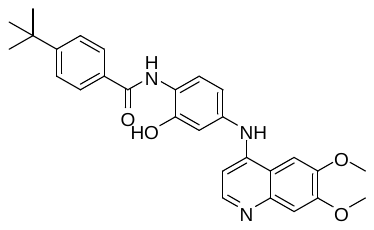

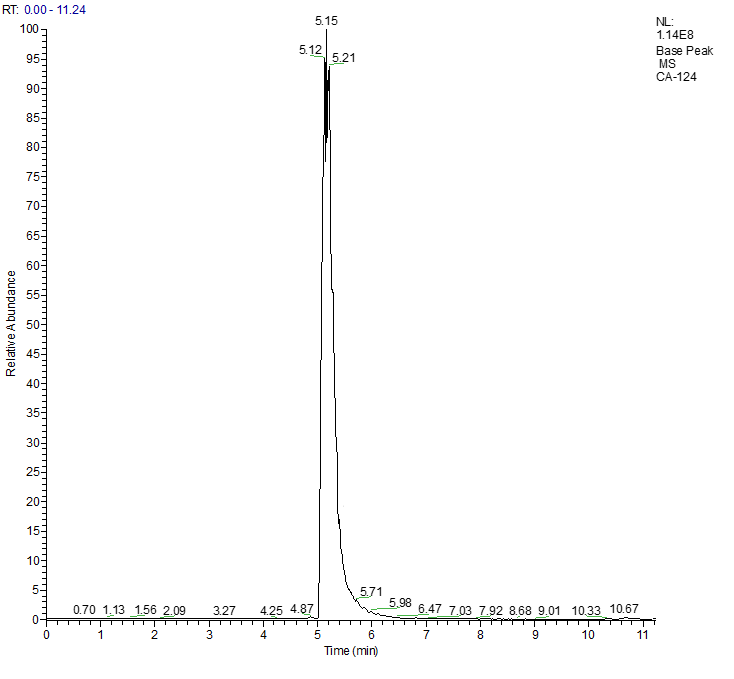

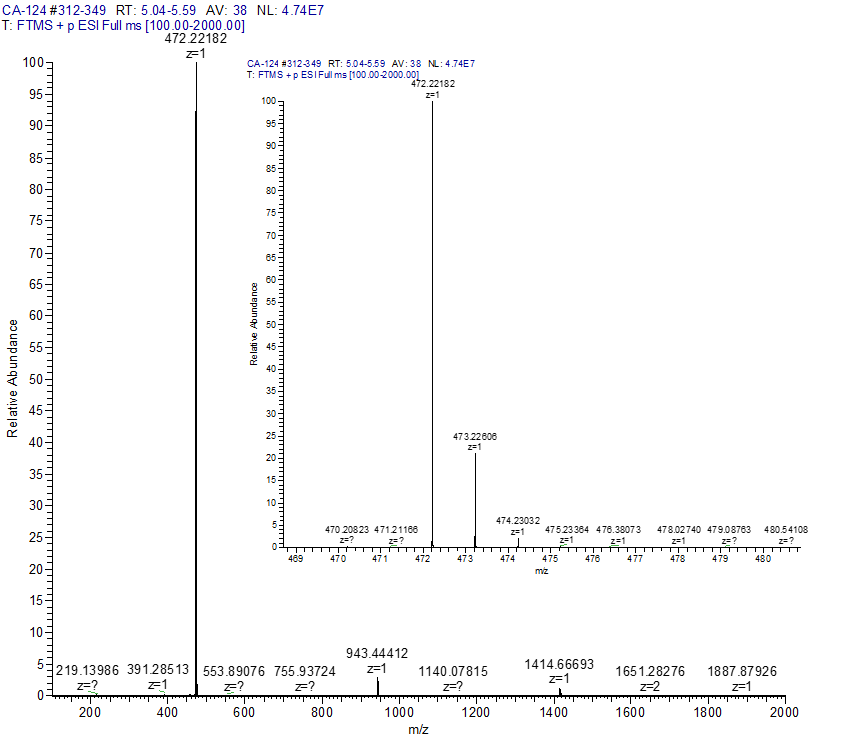

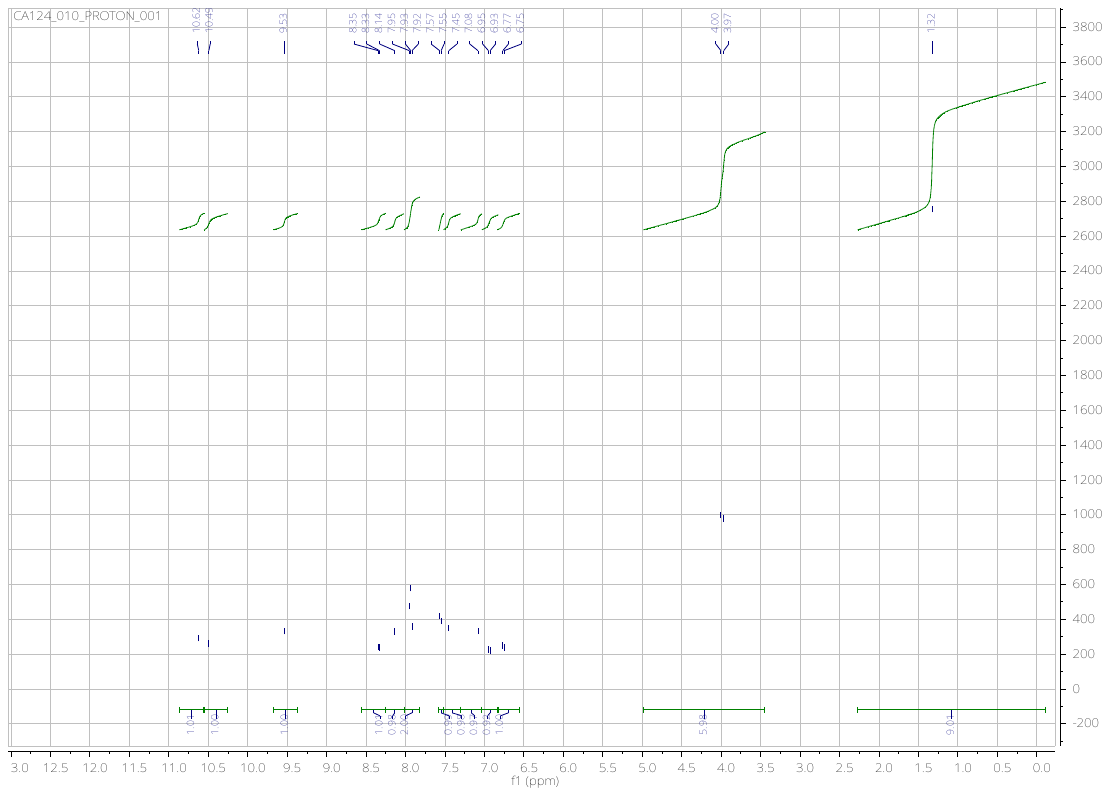

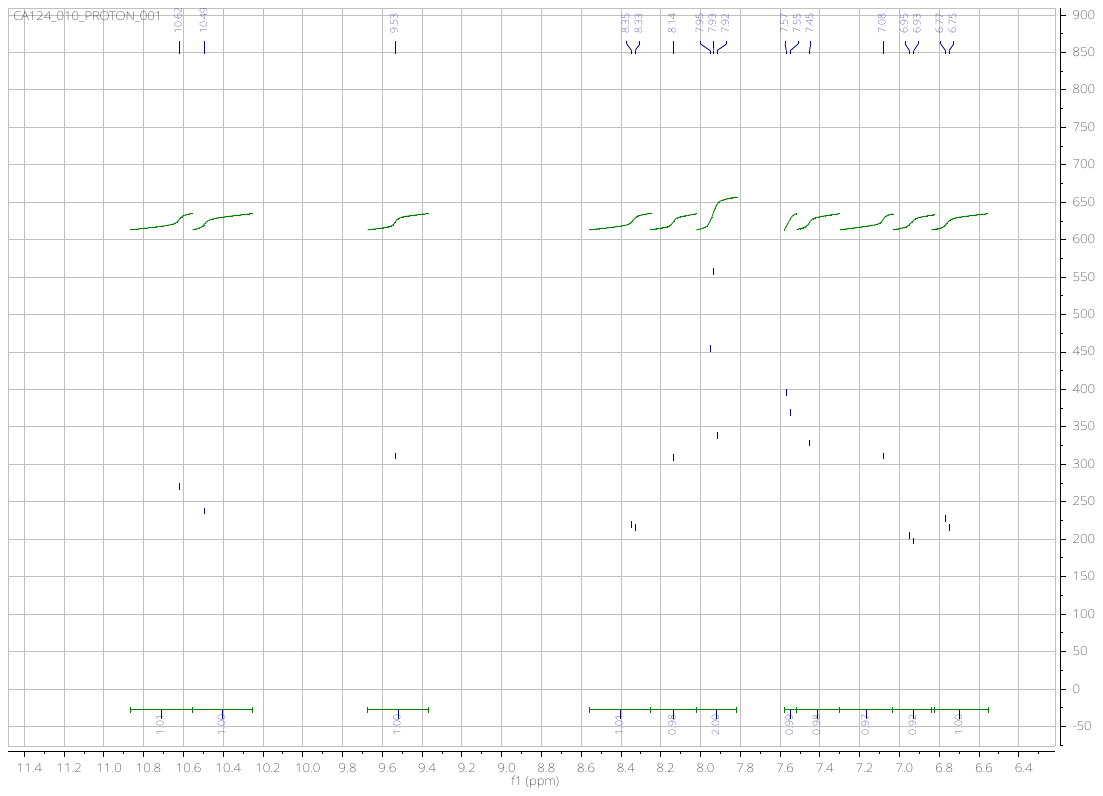

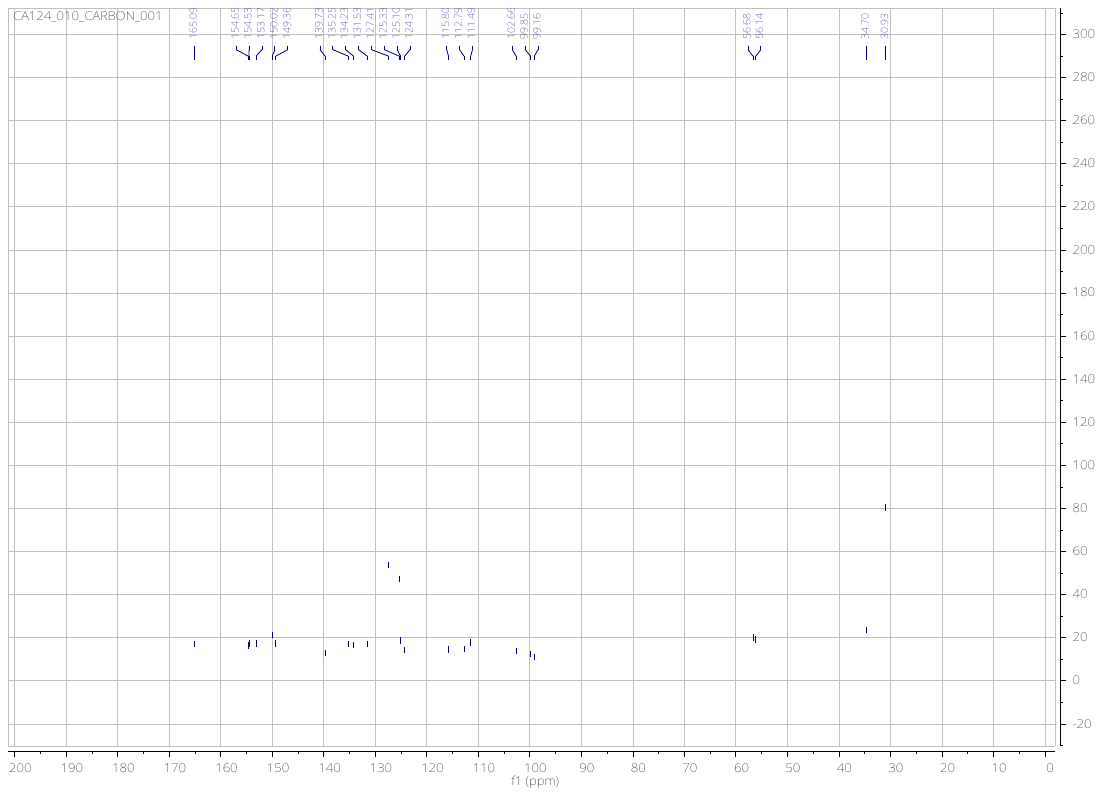

6,7-dimethoxy-*N*-(4-((4-methylbenzyl)oxy)phenyl)quinolin-4-amine (**18**)

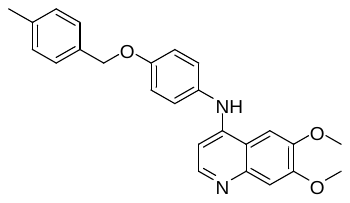

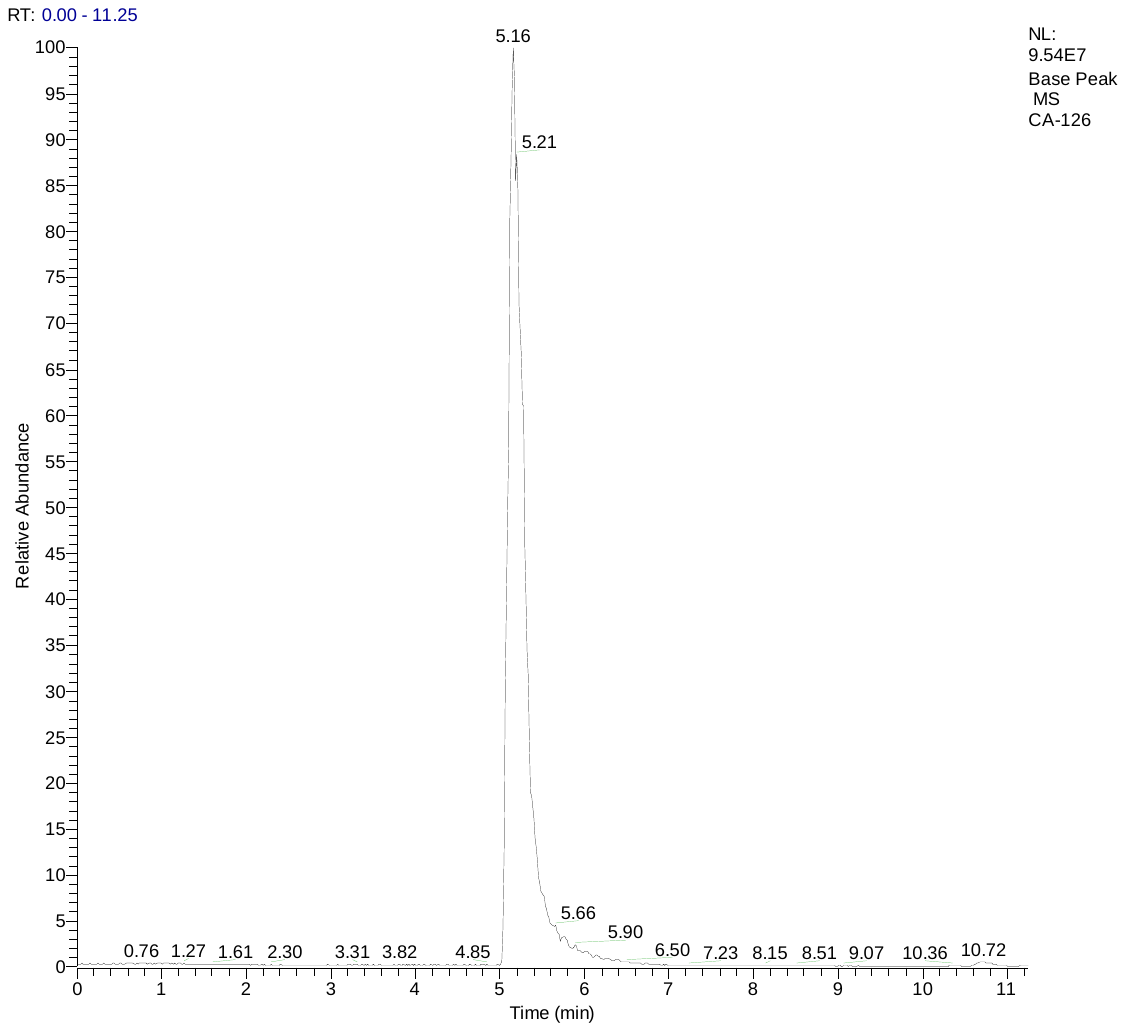

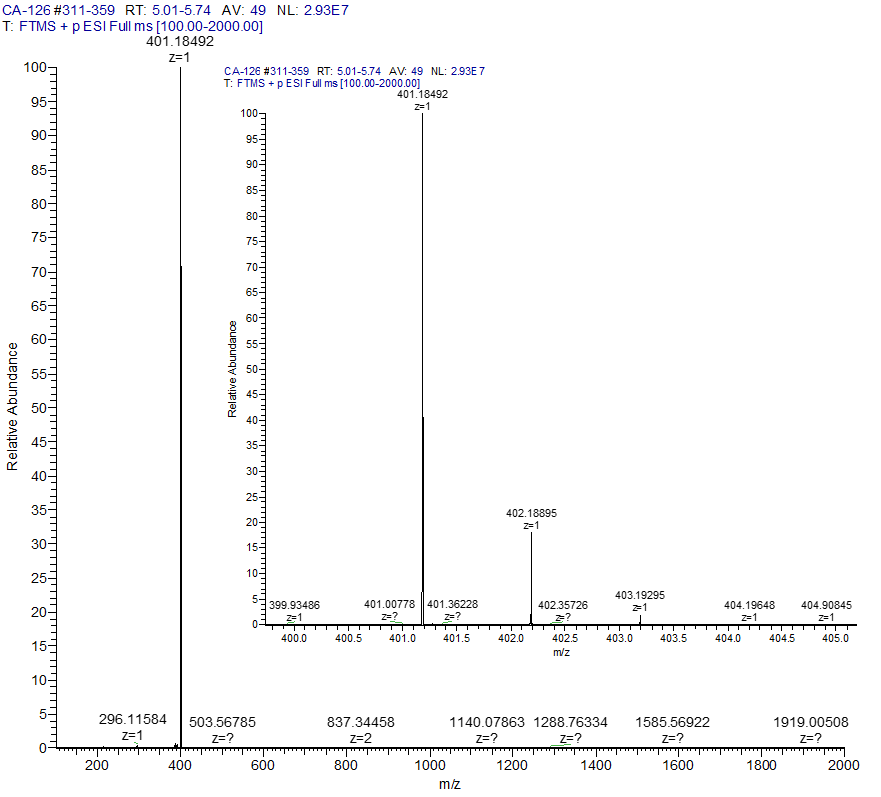

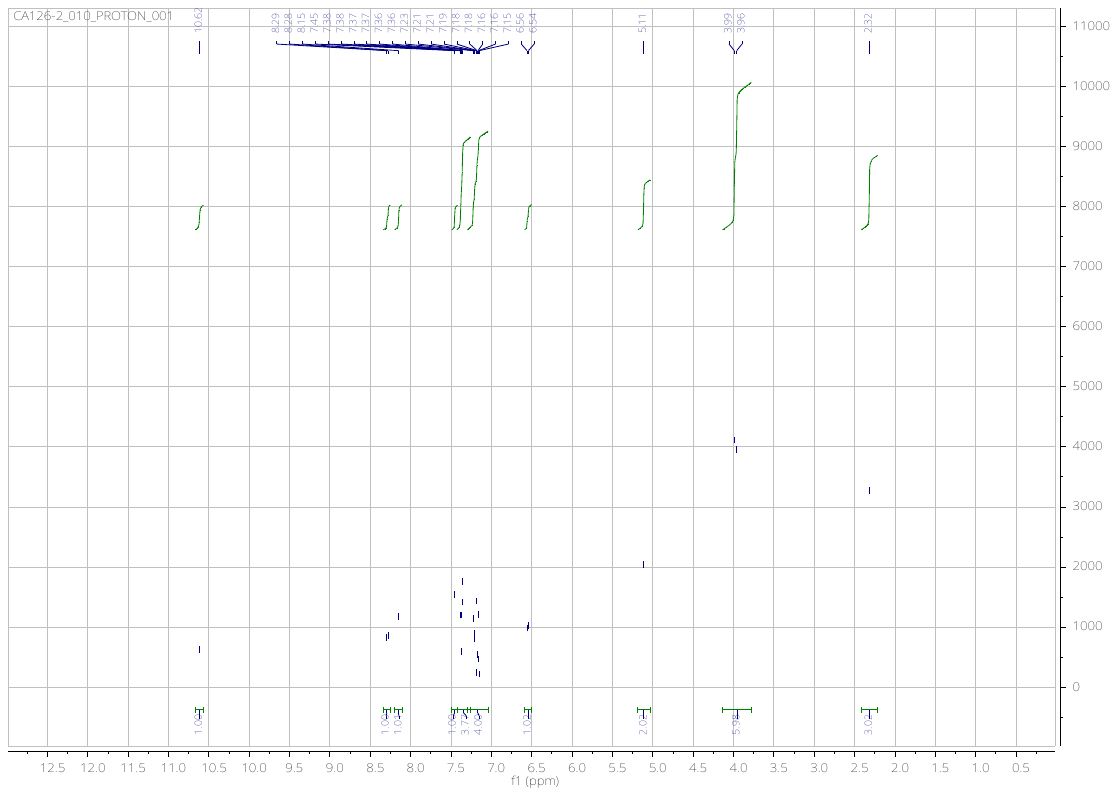

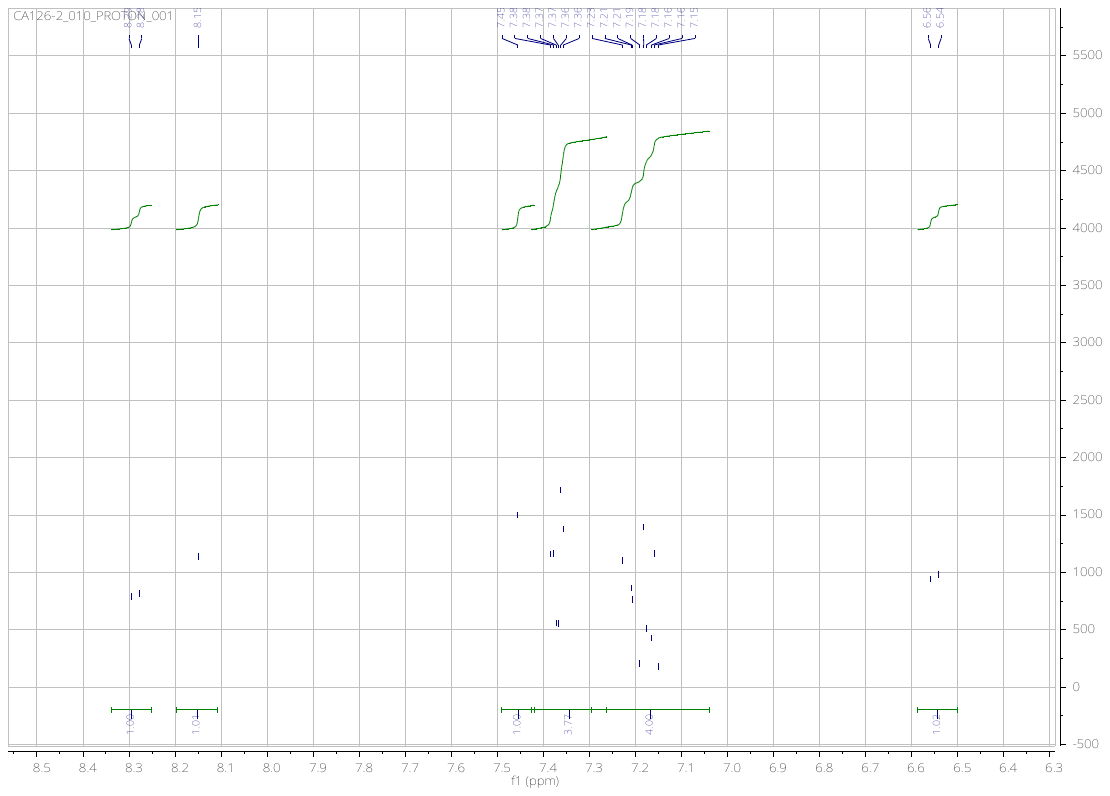

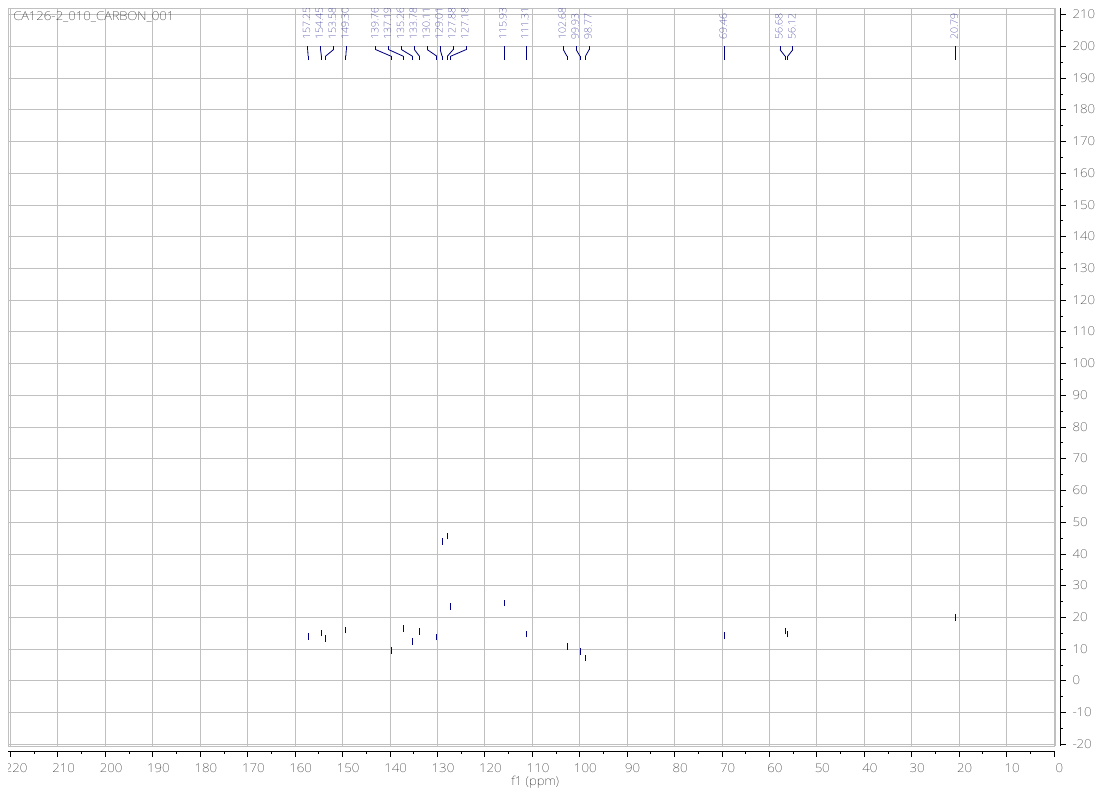

*N*-(4-((6,7-dimethoxyquinolin-4-yl)amino)phenyl)acetamide (**19**)

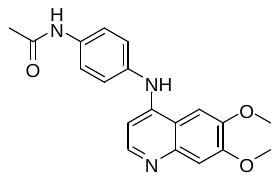

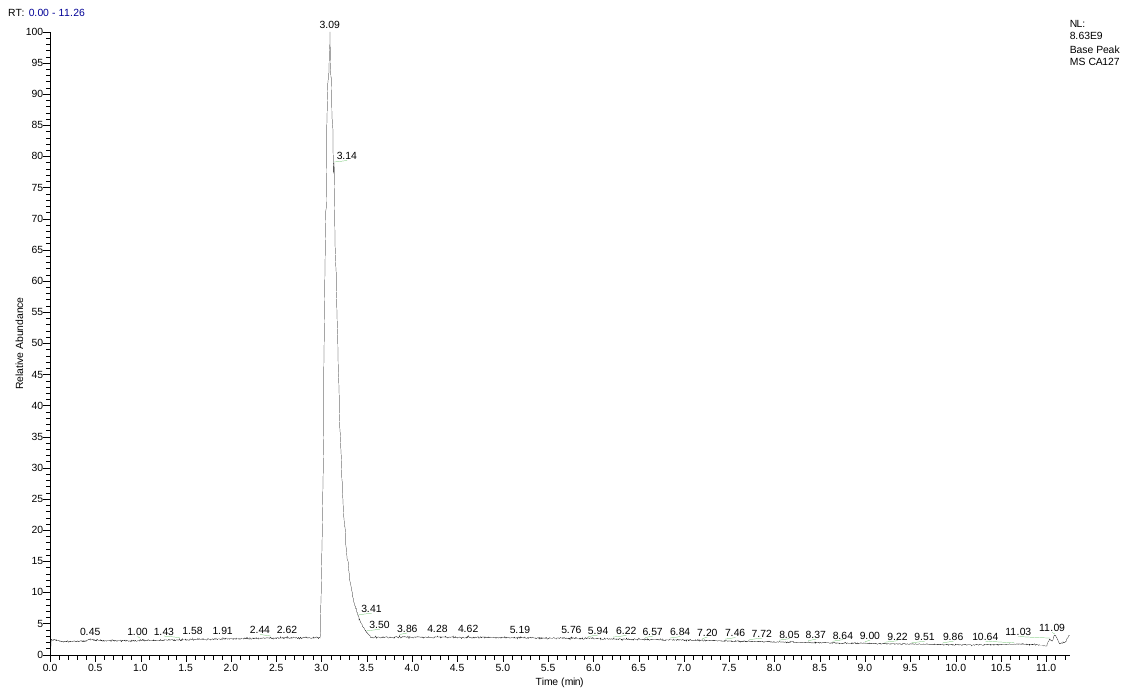

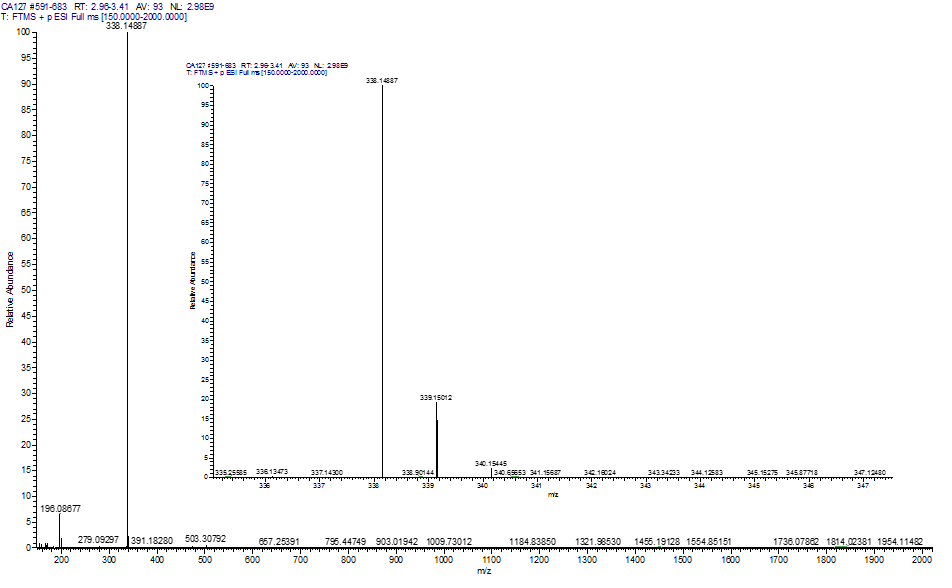

**1.2. Smiles and Labbook codes for final compounds**

| Number | Lab Book | SIMILES |
| --- | --- | --- |
| 1 | CA156 | C#CC1=CC=CC(NC2=CC=NC3=CC(OC)=C(OC)C=C32)=C1 |
| 2 | CA2-158 | C#CC1=CC=CC(NC2=CC=NC3=CC=C(OC)C=C32)=C1 |
| 3 | CA2-159 | C#CC1=CC=CC(NC2=CC=NC3=CC(OC)=CC=C32)=C1 |
| 4 | CA176 | C#CC1=CC=CC(NC2=NC=NC3=CC(OC)=C(OC)C=C32)=C1 |
| 5 | CA204 | COC1=CC2=C(NC3=CC=CC(C#C)=C3)N=CN=C2C=C1 |
| 6 | CA209 | COC1=CC2=NC=NC(NC3=CC=CC(C#C)=C3)=C2C=C1 |
| 7 | CA252 | C#CC1=CC=CC(NC2=C(C#N)C=NC3=CC(OC)=C(OC)C=C32)=C1 |
| 8 | CA251 | N#CC1=C(NC2=CC=CC(C#C)=C2)C3=CC(OC)=CC=C3N=C1 |
| 9 | CA287 | N#CC1=C(NC2=CC=CC(C#C)=C2)C3=CC=C(OC)C=C3N=C1 |
| 13 | CA95 | O=C(NC1=CC=C(NC2=CC=NC3=CC(OC)=C(OC)C=C23)C=C1O)C4=CC=C(C)C=C4 |
| 14 | CA97 | O=C(NC1=CC=C(NC2=CC=NC3=CC=C(OC)C=C23)C=C1O)C4=CC=C(C)C=C4 |
| 15 | CA96 | O=C(NC1=CC=C(NC2=CC=NC3=CC(OC)=CC=C23)C=C1O)C4=CC=C(C)C=C4 |
| 16 | CA2-148 | O=C(NC1=CC=C(NC2=CC=NC3=CC(OC)=C(OC)C=C23)C=C1)C4=CC=C(C)C=C4 |
| 17 | CA124 | O=C(NC1=CC=C(NC2=CC=NC3=CC(OC)=C(OC)C=C23)C=C1O)C4=CC=C(C(C)(C)C)C=C4 |
| 18 | CA126 | CC1=CC=C(C=C1)COC2=CC=C(NC3=CC=NC4=CC(OC)=C(OC)C=C34)C=C2 |
| 19 | CA127 | CC(NC1=CC=C(NC2=CC=NC3=CC(OC)=C(OC)C=C23)C=C1)=O |
| - | Erlotinib | C#CC1=CC=CC(NC2=NC=NC3=CC(OCCOC)=C(OCCOC)C=C32)=C1 |

**1.3. EGFR in cell phosphorylation curves and method**

ProQinase GmbH performed EGFR in-cell assays. Briefly - Compounds were tested for their inhibitory impact on the cellular kinase activity of EGF-R, assessed by the measurement of autophosphorylation. In the cellular EGF-R phosphorylation assay the human epidermoid carcinoma cell line A431 is used, which expresses endogenously a high level of EGF-R. Stimulation of these cells with human epidermal growth factor (EGF) results in receptor tyrosine autophosphorylation. A431 cells were plated in RPMI supplemented with 10% FCS in multi-well cell culture plates. After serum-starvation overnight cells were incubated with compounds in serum-free medium. Raw data were converted into percent substrate phosphorylation relative to high controls, which were set to 100%. IC_50_ values were determined using GraphPad Prism 5 software with constrain of bottom to 0 and top to 100 using a nonlinear regression curve fit with variable hill slope. The equation is a four-parameter logistic equation. IC_50_ curves for indicated compounds on EGF-R mediated autophosphorylation. Raw data (OD450 nM - 540 nM) were converted into percent substrate phosphorylation relative to solvent controls, which were set to 100%. Graphical illustration was performed using GraphPad Prism 5 software.

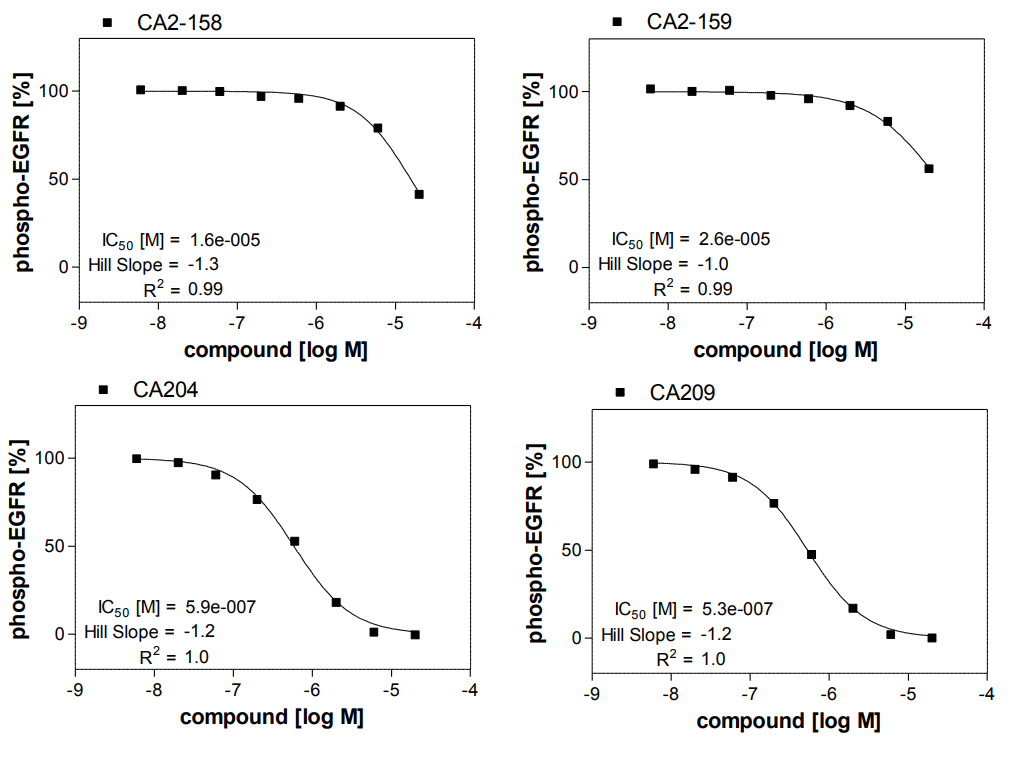

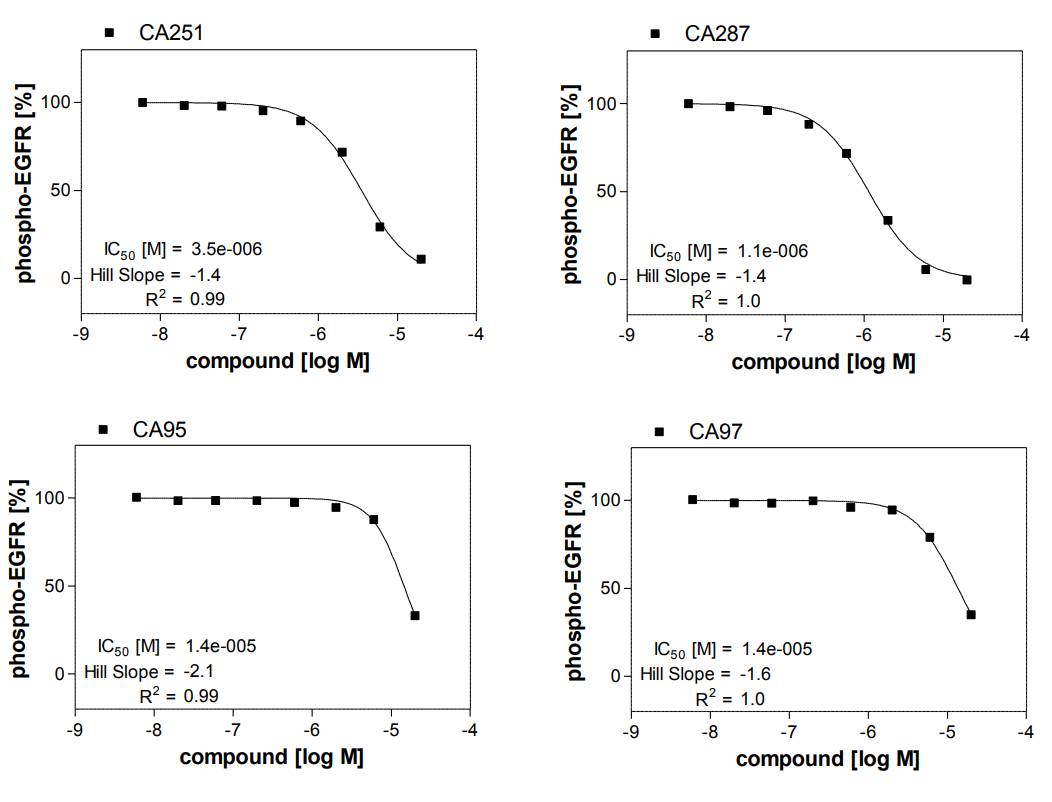

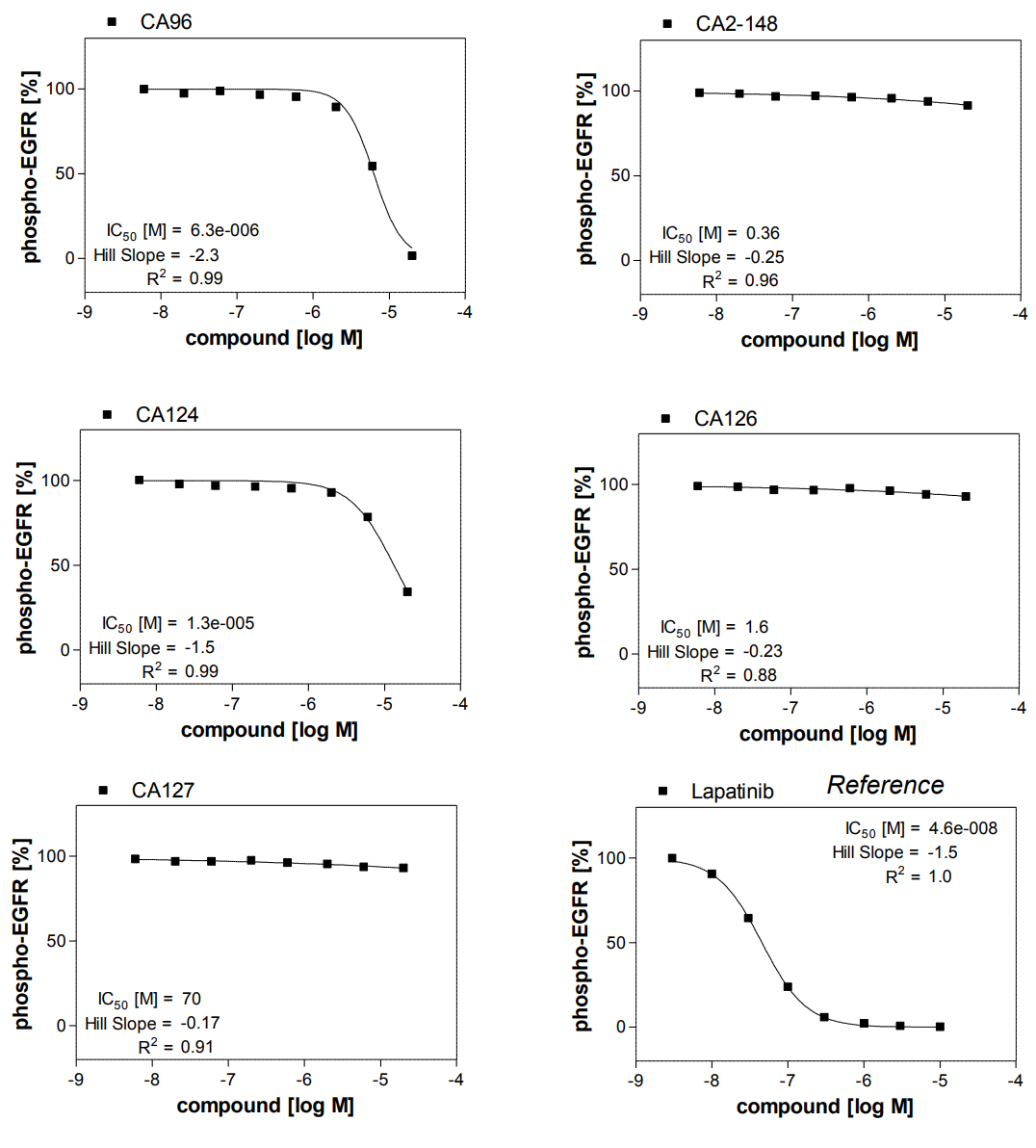

**1.4. IC_50_ curves for A431, UCH-1, UCH-2 and WS-1**

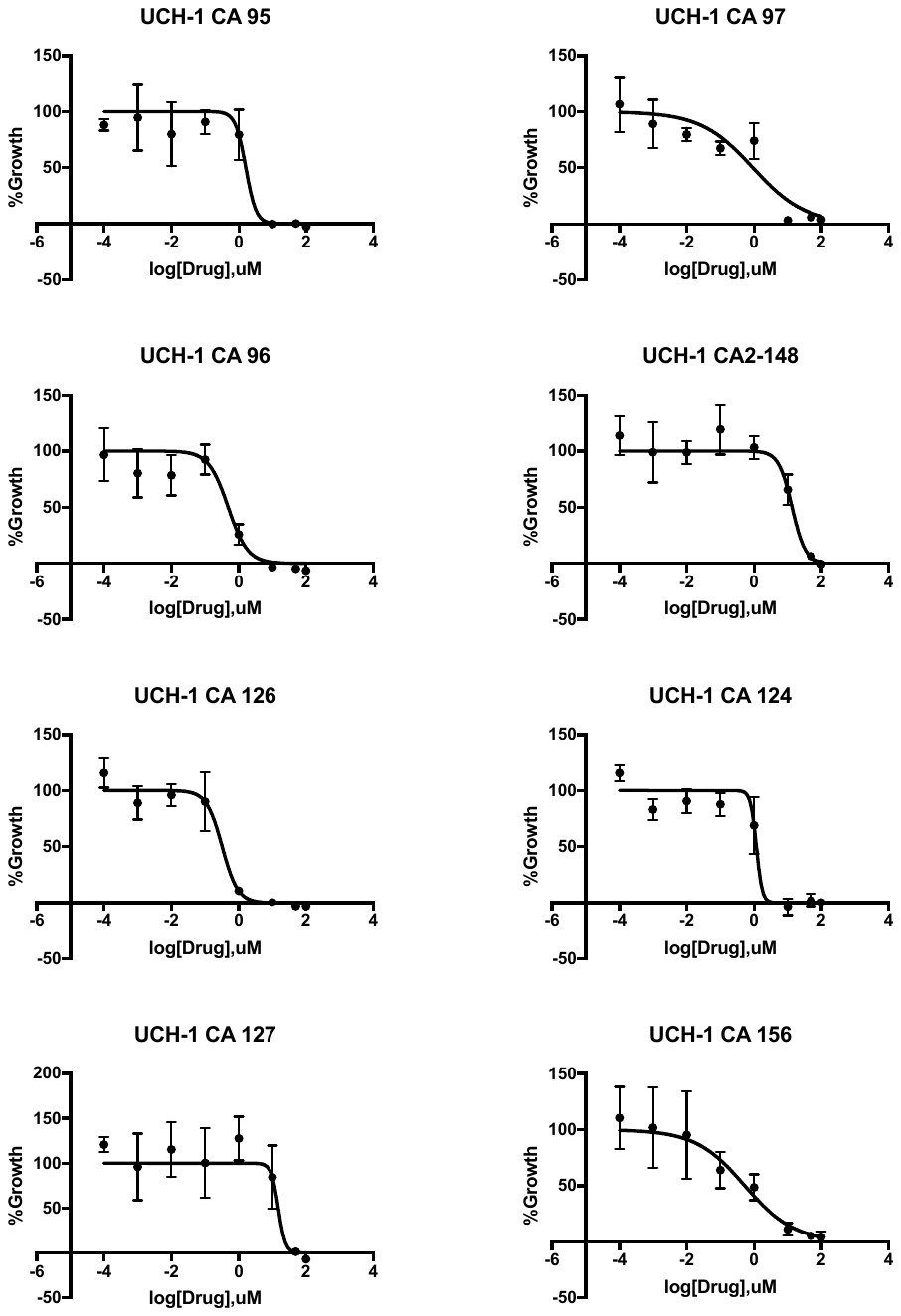

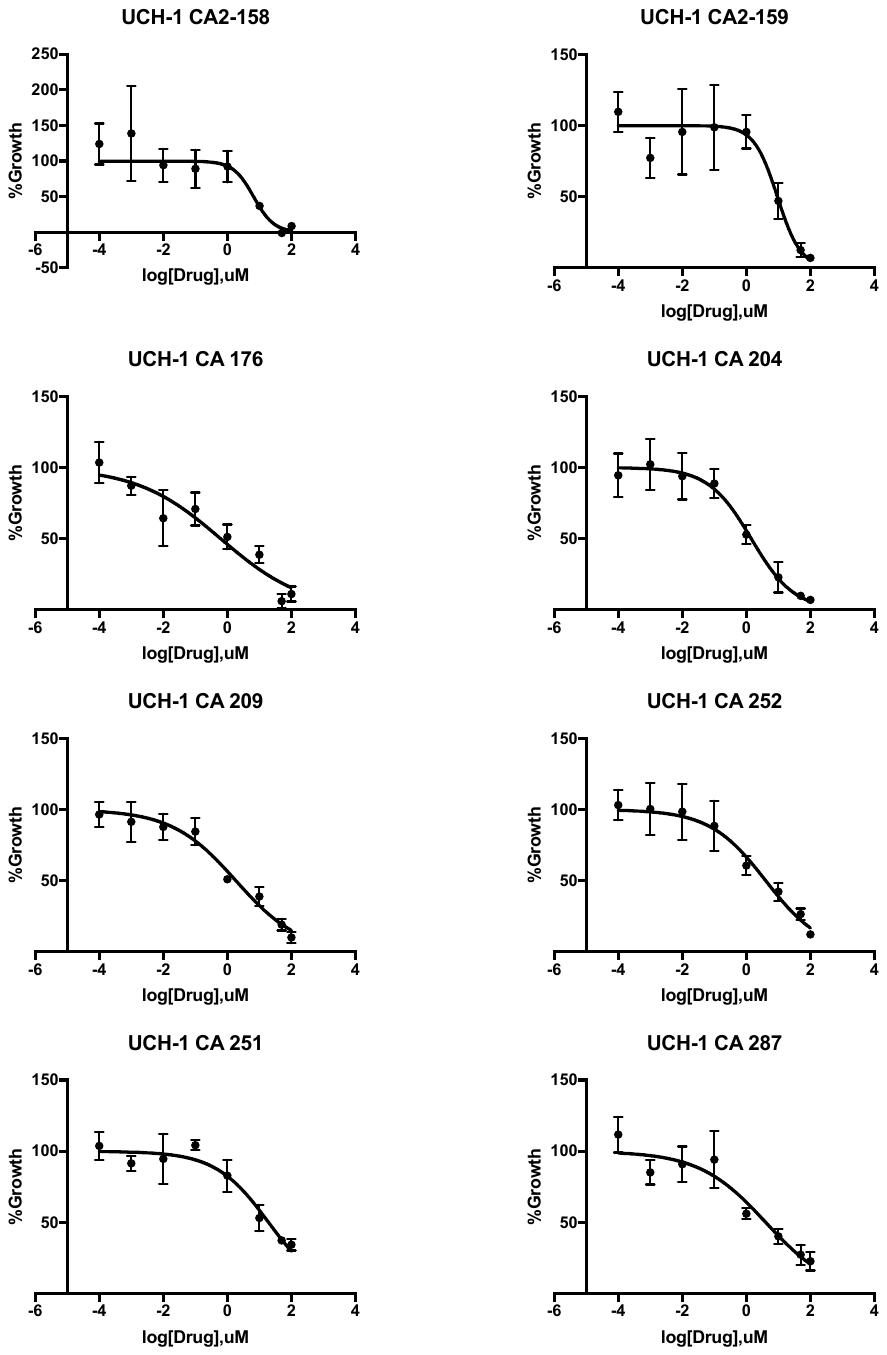

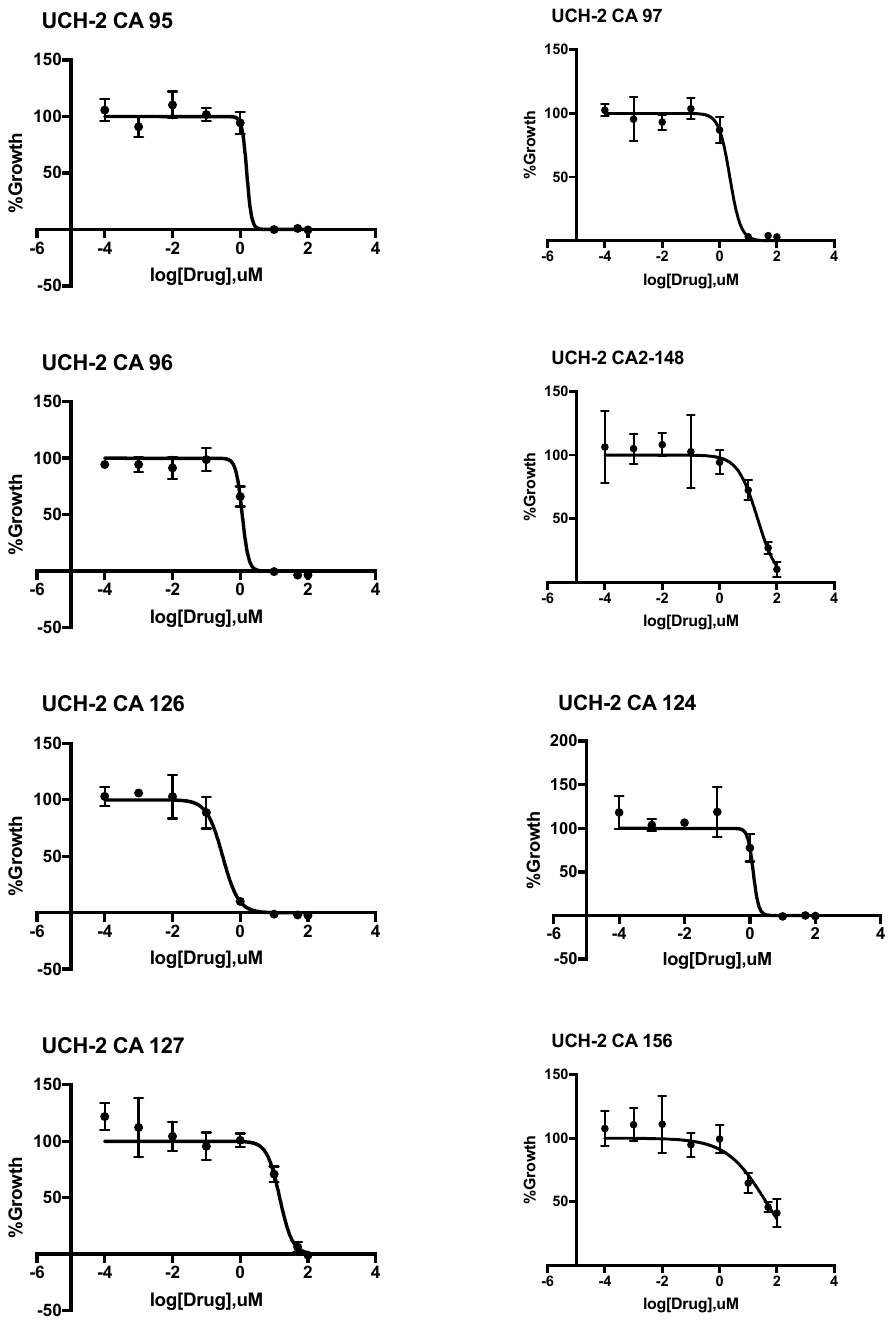

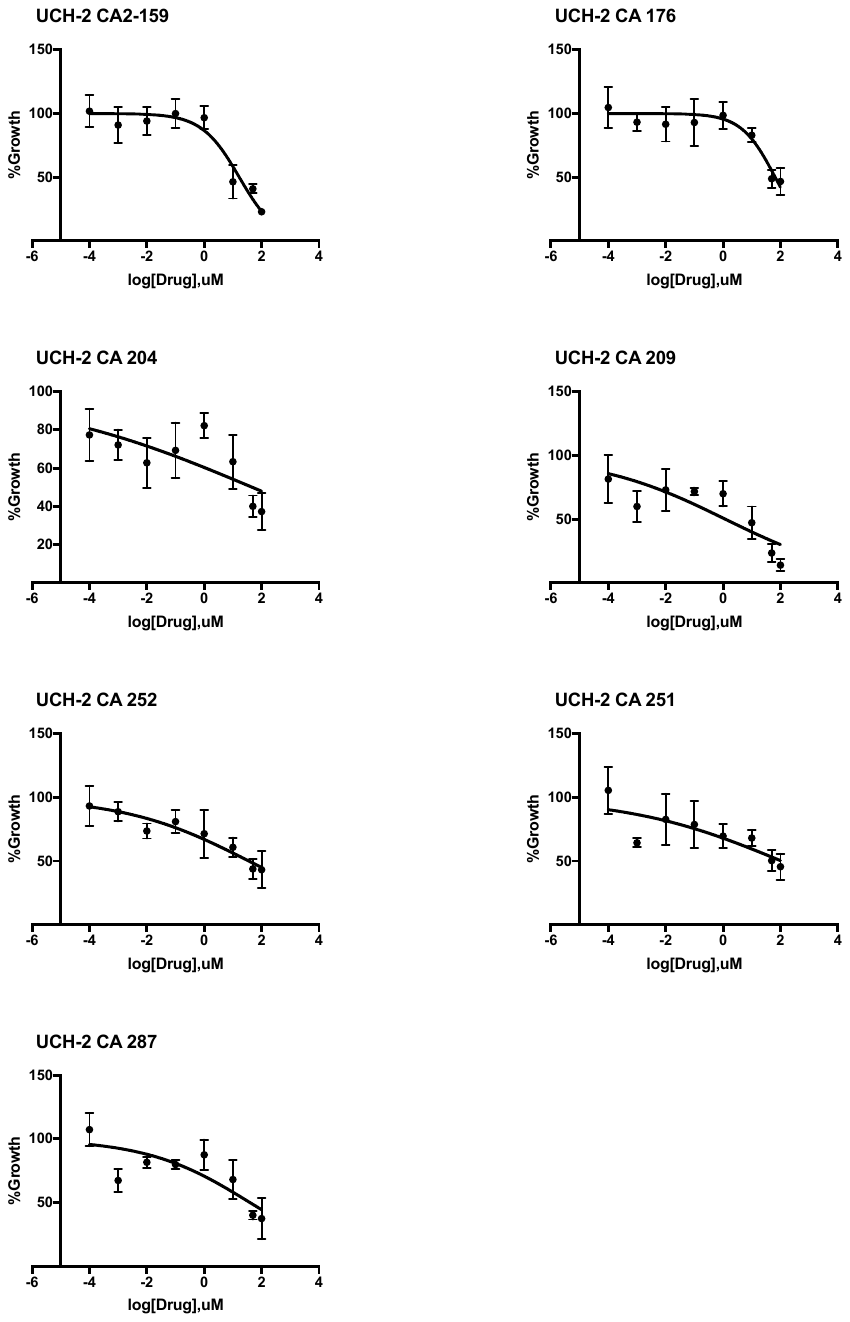

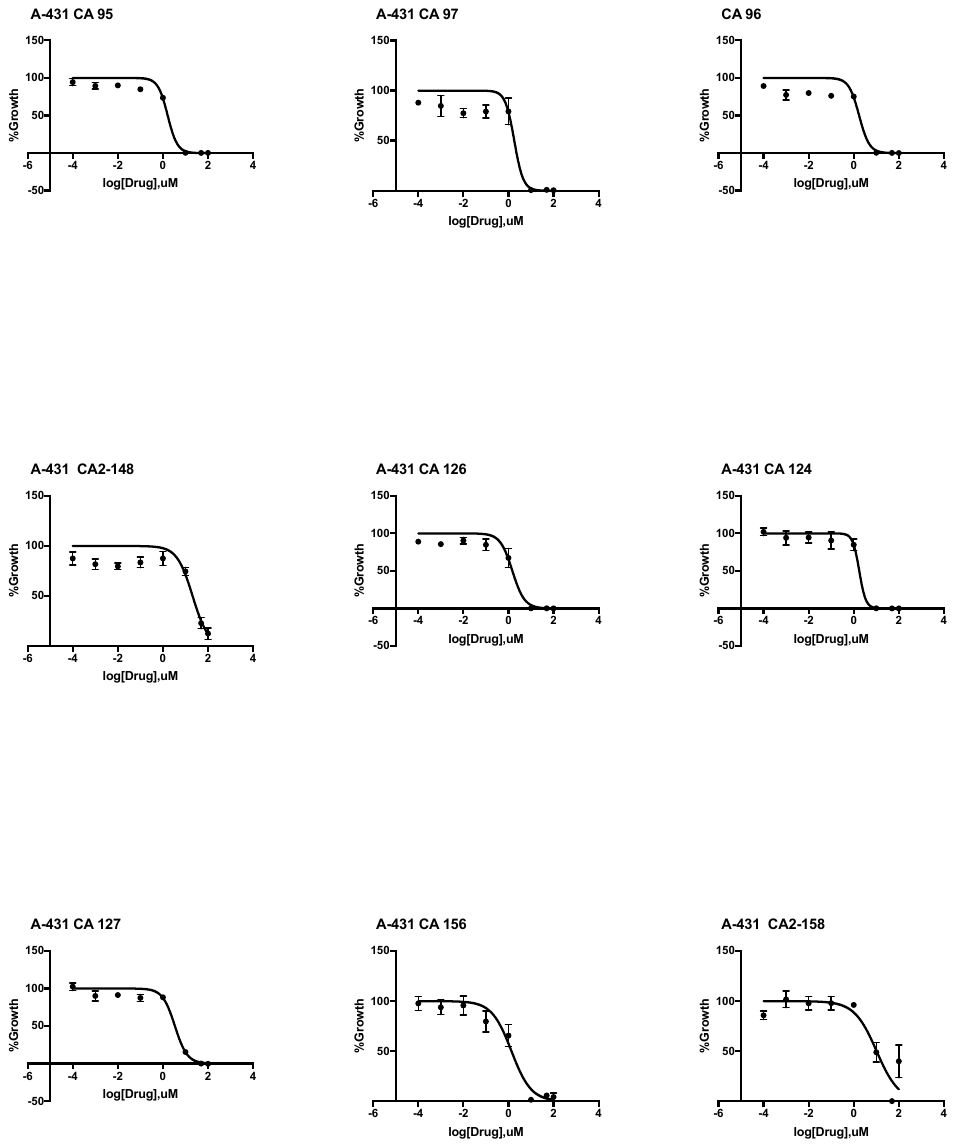

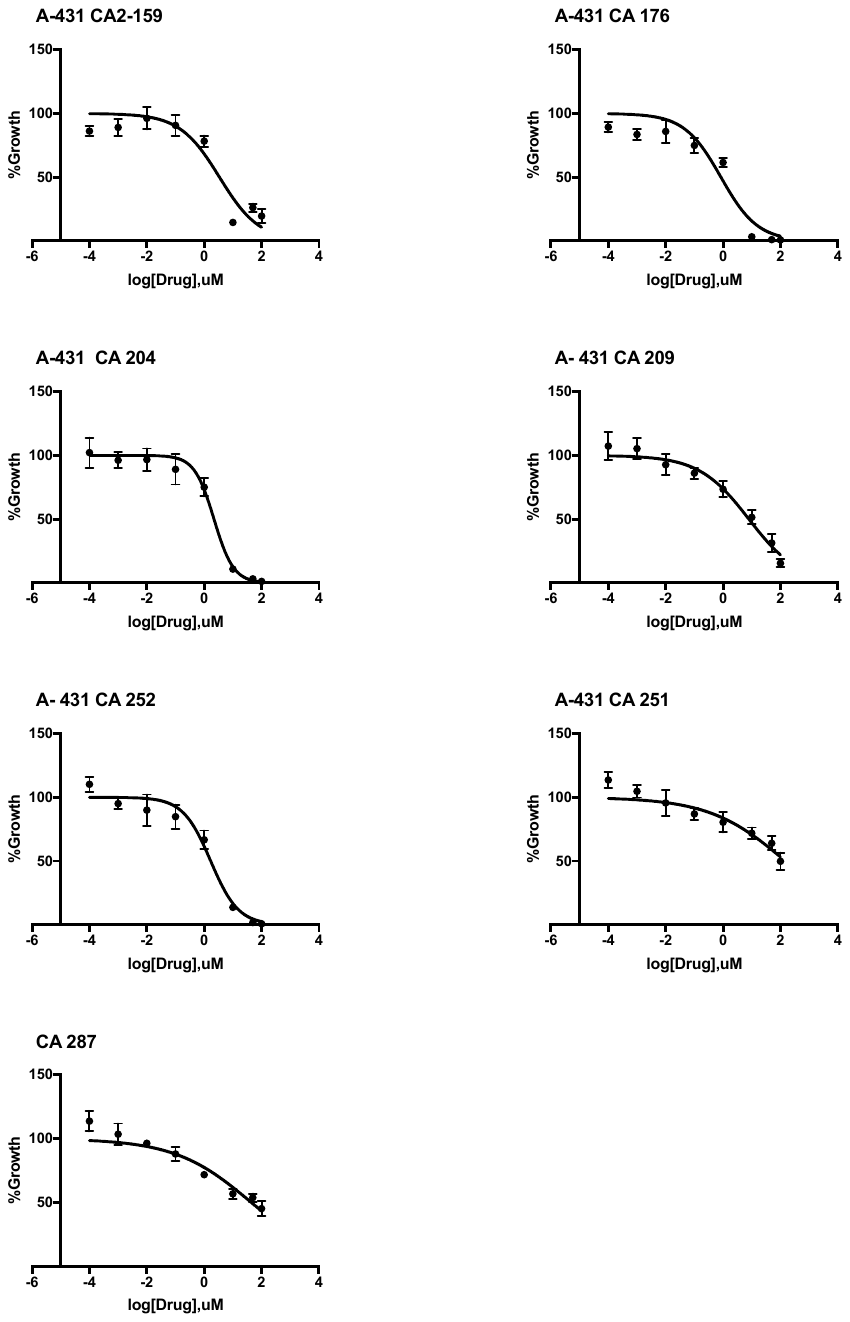

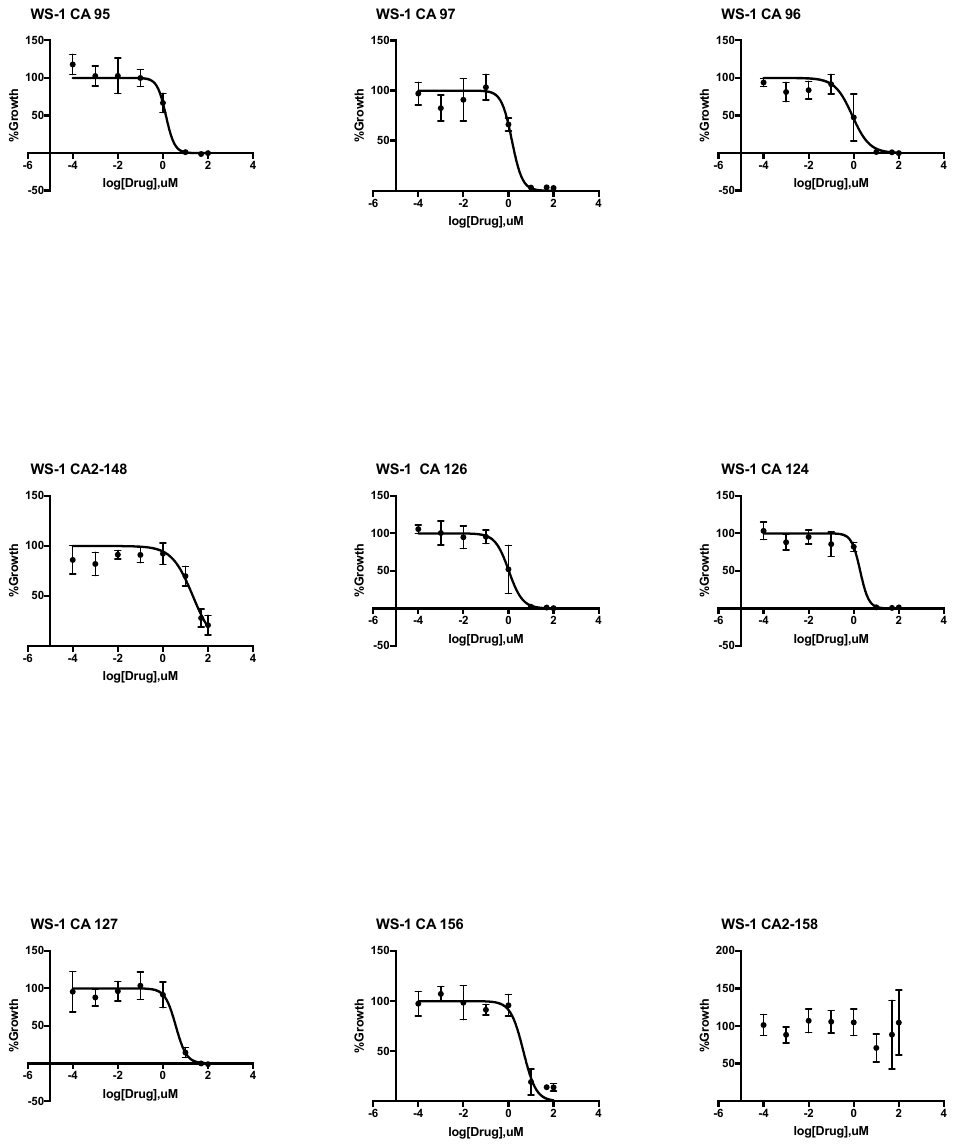

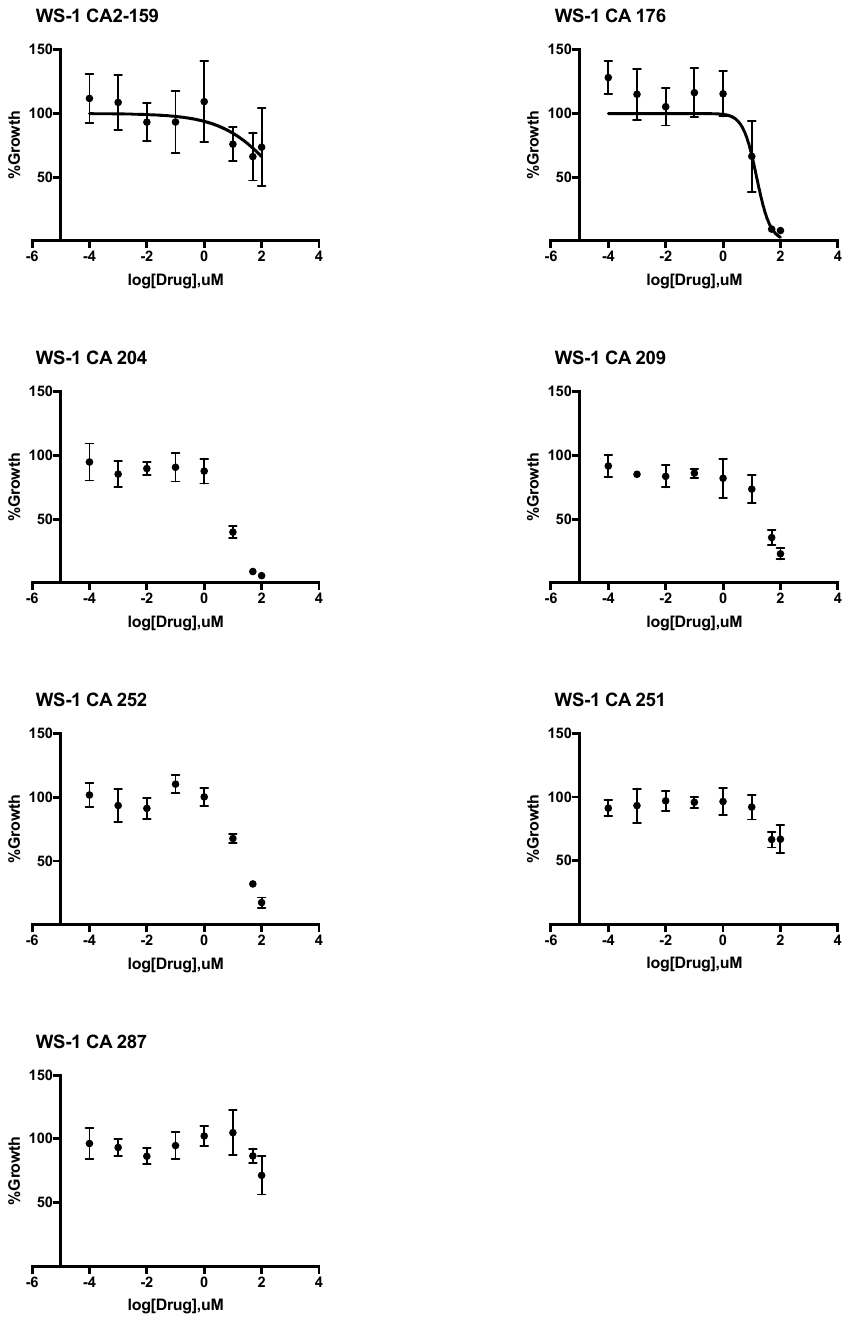

**1.5. Mass Spectrometry method**

Samples were analyzed with a ThermoFisher Q Exactive HF-X (ThermoFisher, Bremen, Germany) mass spectrometer coupled with a Waters Acquity H-class liquid chromatograph system. Samples were introduced via a heated electrospray source (HESI) at a flow rate of 0.6 mL/min. Electrospray source conditions were set as: spray voltage 3.0 kV, sheath gas (nitrogen) 60 arb, auxillary gas (nitrogen) 20 arb, sweep gas (nitrogen) 0 arb, nebulizer temperature 375 degrees C, capillary temperature 380 degrees C, RF funnel 45 V. The mass range was set to 150-2000 m/z. All measurements were recorded at a resolution setting of 120,000.

Separations were conducted on a Waters Acquity UPLC BEH C18 column (2.1 x 50 mM, 1.7 uM particle size). LC conditions were set at 100 % water with 0.1 % formic acid (A) ramped linearly over 9.8 mins to 95 % acetonitrile with 0.1 % formic acid (B) and held until 10.2 mins. At 10.21 mins the gradient was switched back to 100% A and allowed to re-equilibrate until 11.25 mins. Injection volume for all samples was 3 uL.

Xcalibur (ThermoFisher, Breman, Germany) was used to analyze the data. Solutions were analyzed at 0.1 mg/mL or less based on responsiveness to the ESI mechanism. Molecular formula assignments were determined with Molecular Formula Calculator (v 1.2.3). All observed species were singly charged, as verified by unit *m/z* separation between mass spectral peaks corresponding to the ^12^C and ^13^C^12^C_c-1_ isotope for each elemental composition.
